# Fragment-based development of small molecule inhibitors targeting *Mycobacterium tuberculosis* cholesterol metabolism

**DOI:** 10.1101/2024.10.28.620643

**Authors:** Madeline E. Kavanagh, Kirsty J. McLean, Sophie H. Gilbert, Cecilia Amadi, Matthew Snee, Richard B. Tunnicliffe, Kriti Arora, Helena I. Boshoff, Alexander Fanourakis, Maria Jose Rebello-Lopez, Fatima Ortega-Muro, Colin W. Levy, Andrew W. Munro, David Leys, Chris Abell, Anthony G. Coyne

## Abstract

*Mycobacterium tuberculosis* (*Mtb*) is the world’s most deadly infectious pathogen and new drugs are urgently required to combat the emergence of multi-(MDR) and extensively-(XDR) drug resistant strains. The bacterium specifically upregulates sterol uptake pathways in infected macrophages and the metabolism of host-derived cholesterol is essential for *Mtb’s* long-term survival *in vivo.* Here, we report the development of antitubercular small molecules that inhibit the *Mtb* cholesterol oxidases CYP125 and CYP142, which catalyze the initial step of cholesterol metabolism. An efficient biophysical fragment screen was used to characterize the structure-activity relationships of CYP125 and CYP142, and identify a non-azole small molecule **1a** that can bind to the heme cofactor of both enzymes. A structure-guided fragment-linking strategy was used to optimize the binding affinity of **1a**, yielding a potent dual CYP125/142 inhibitor **5m** (K_D_ CYP125/CYP142 = 0.04/0.16 µM). Compound **5m** potently inhibits the catalytic activity of CYP125 and CYP142 *in vitro* (K_I_ values < 0.1 µM), and rapidly depletes *Mtb* intracellular ATP (IC_50_ = 0.15 µM). The compound has antimicrobial activity against both drug susceptible and MDR *Mtb (*MIC_99_ values 0.4 - 1.5 µM) in extracellular assays, and inhibits the growth of *Mtb* in human macrophages (MIC = 1.7 µM) with good selectivity over mammalian cytotoxicity (LD_50_ ≥ 50 µM). The combination of small molecule inhibitors and structural data reported here provide useful tools to study the role of cholesterol metabolism in *Mtb* and are a promising step towards novel antibiotics targeting bioenergetic pathways, which could be used to help combat MDR-TB.

## Introduction

Tuberculosis (TB) is the world’s most deadly infectious disease, killing more than 1.3 million people every year.^1^ Although global TB deaths are declining, there has been an alarming increase in the number and distribution of cases caused by multi-(MDR) or extensively-(XDR) drug resistant strains of the causal bacterium *Mycobacterium tuberculosis* (Mtb). Despite this impending threat, only two drugs (bedaquiline and pretomanid) with new mechanisms of action (MoA) have been approved for the treatment of TB in more than 50 years. Consequently, there is now an urgent need to develop new anti-tubercular drugs, in particular, compounds with activity against recalcitrant *Mtb* populations, such as non-replicating bacteria and MDR-TB.^2^

*Mtb* is a facultative intracellular pathogen with unique metabolic adaptations that enable the bacteria to survive long-term in the harsh, nutrient poor environment of the host macrophage.^3–6^ The development of drugs that specifically target bioenergetic pathways required for intracellular growth has recently emerged as a promising approach that could help address limitations of first and second line drugs^2,7^ For example, bedaquiline, a diarylquinoline that targets the *Mtb* ATP synthase,^8^ is active against both replicating and dormant *Mtb,*^9^ and has improved efficacy against intracellular bacteria, which are typically less sensitive to standard TB drugs.^10,11^ Numerous studies have also demonstrated that the ability of bedaquiline to modulate *Mtb* metabolism helps counteracts drug resistance mechanisms,^12^ synergizes with existing drugs,^13^ and may enhance the antibacterial activity of host macrophages.^2,7,9,13,14^ Since the approval of bedaquiline in 2012, several other compounds targeting bacterial respiration or bioenergetic pathways, including clofazimine,^15^ and the cytochrome bc1 complex inhibitor telacebec (Q203),^16,15^ have entered clinical trials, and are showing promising efficacy against recalcitrant *Mtb* populations, including non-replicating bacteria and MDR-TB.^2,9,17^

Unlike other bacteria, *Mtb* is able to simultaneously utilize diverse carbon sources to support growth *in vivo*.^18^ For example, during infection *Mtb* primarily relies on the metabolism of host-derived fatty acids and cholesterol for energy and biosynthetic building blocks.^19–22^ Specifically the acetyl CoA derived from the breakdown of cholesterol is shuttled into the TCA cycle to produce ATP, while propionyl CoA is incorporated into virulence-associated cell wall lipids.^19^ *Mtb’s* ability to dysregulate sterol homeostasis also modulates the host immune response, producing a more permissive intracellular environment that enables chronic infection.^19,23–25^ Consequently, the development of drugs that inhibit *Mtb* cholesterol metabolism would both target bioenergetic pathways that are required for the bacteria’s long-term persistence, decrease *Mtb* fitness, and support the host immune response.^26,27^

The first step of cholesterol degradation in *Mtb* ― C27-oxidation of the cholesterol/enone side chain ― is catalyzed by a 48 kDa cytochrome p450 enzyme (P450), CYP125.^28–31^ *Cyp125* (*Rv3545c*) is encoded in the *Mtb i*ntracellular *gr*owth (*igr*) operon,^32^ which is widely conserved across actinomycetes and is essential for *Mtb* survival in macrophages^33^ and mice.^5^ The expression of CYP125 is upregulated during infection or when *Mtb* is cultured in the presence of cholesterol,^25,29^ and *Mtb* CYP125 knockout (Δ*Cyp125)* are unable to grow on cholesterol as the sole source of carbon.^25^ Furthermore, Δ*Cyp125 Mtb* mutants are unable to grow on rich media supplemented with cholesterol, because of the accumulation of the toxic CYP125 substrate cholestenone.^21,25,29,31^ Interestingly, certain strains of *Mtb*, including the common laboratory model *Mtb* H37Rv, express an second P450 enzyme CYP142, which can also oxidize cholesterol, and rescues the growth of *ΔCYP125 Mtb*.^25,28,34^ The catalytic efficiency CYP125 and CYP142 is similar, however, they synthesize the opposite stereoisomers of 26-hydroxycholes-4-en-3-one and are only distantly evolutionarily related.^29,34,35^ This partial functional redundancy in the *Mtb* genome highlights the importance of maintaining the integrity of the cholesterol metabolic pathway, and also presents challenges for the development of CYP125/142 inhibitors.

Despite the role of cholesterol metabolism for *Mtb* virulence being identified more than 15 years ago,^19^ there has been little progress in the development of drugs to inhibit this pathway.^36^Imidazole-containing antifungal drugs (e.g. econazole, clotrimazole), which target the fungal P450 CYP51, bind tightly to the heme-cofactor of several *Mtb* P450s,^28,35–40^ and inhibit the growth of *Mtb.*^41^ However, the anti-tubercular activity of these drugs is not dependent on cholesterol, and they have comparatively weak binding affinity to CYP125/142 (*K_D_* values > 1 µM) compared to other essential *Mtb* P450s, which is likely due to the relatively narrow active site channel of the cholesterol oxidases.^28,35^ In addition, the imidazole antifungals are generally not considered suitable for treating TB because of their susceptibility to *Mtb* azole efflux transporters,^42^ and potential to cause adverse side effects and drug-drug interactions due to inhibition of mammalian P450s.^43^ Our lab previously reported preliminary results from a fragment-based screening campaign targeting CYP125, which yielded several hit fragments that were validated to bind CYP125 by differential scanning fluorimetry (DSF) and ligand-observed NMR.^44^ However, no further optimization of the compounds was attempted because we could not obtain high quality X-ray crystal structures of ligand-bound CYP125.

Here, we report the development of dual CYP125/142 inhibitors, which inhibit the growth of *Mtb* in extra- and intracellular assays. We initially leverage an efficient biophysical screening strategy to characterize the CYP125/142 fragment binding profile and identify a non-imidazole hit **1a** that might be more potent, selective, and less susceptible to azole efflux transporters than the anti-fungal drugs. We subsequently employ CYP142 as a structural proxy to guide hit-to-lead optimization of dual CYP125/142 inhibitors, which have low nanomolar binding affinity and inhibit CYP125/142 catalytic activity *in vitro*. Finally, we demonstrate that these novel CYP125/142 inhibitors have antimicrobial activity against extracellular *Mtb* (including MDR-TB), and *Mtb* in human macrophages. The combination of small molecule inhibitors and structural data reported here, provides a promising step towards the development of chemical probes to study the role of cholesterol metabolism for *Mtb* virulence *in vivo*, and the development of novel antibiotics to combat MDR-TB. This research has also supported the development of a subsequent series of CYP125 inhibitors with antitubercular activity.^45^

## Results

### Fragment screening identifies preferred CYP125/142 ligand

A focused library of eighty fragments was assembled in order to characterize the preferred heme-binding chemotype of CYP125 and CYP142, and to identify a common chemical scaffold that could be used for the development of a dual CYP125/142 inhibitor (Table S1).^39^ Each fragment in the library contained an aliphatic or aromatic amine, however, imidazole-containing fragments were specifically deprioritized (≤10%), because of the promiscuity of this functional group for binding to both human and microbial P450s, and sensitivity to azole efflux transporters.^42,46,47^ This library was screened against a panel of purified *Mtb* P450s (Fig. 1a), including CYP125 and CYP142, by UV-vis spectroscopy to identify fragments that induced a red-shift in the *λ_max_* of the enzymes absorbance spectrum. All P450s have a unique absorbance spectrum that reflects the co-ordination environment of heme iron, and small molecules that co-ordinate directly to ferric heme using a strong field ligand (e.g., nitrogen) cause a red shift in the *λ_max_*, which typically indicates stabilization of the inactive, low spin state of the enzyme.^48^ This spectral property makes UV-vis a highly efficient method to identify small molecules with the potential to inhibit P450 activity, and to infer their binding orientation, in the absence of structural data.

**Figure 1.**
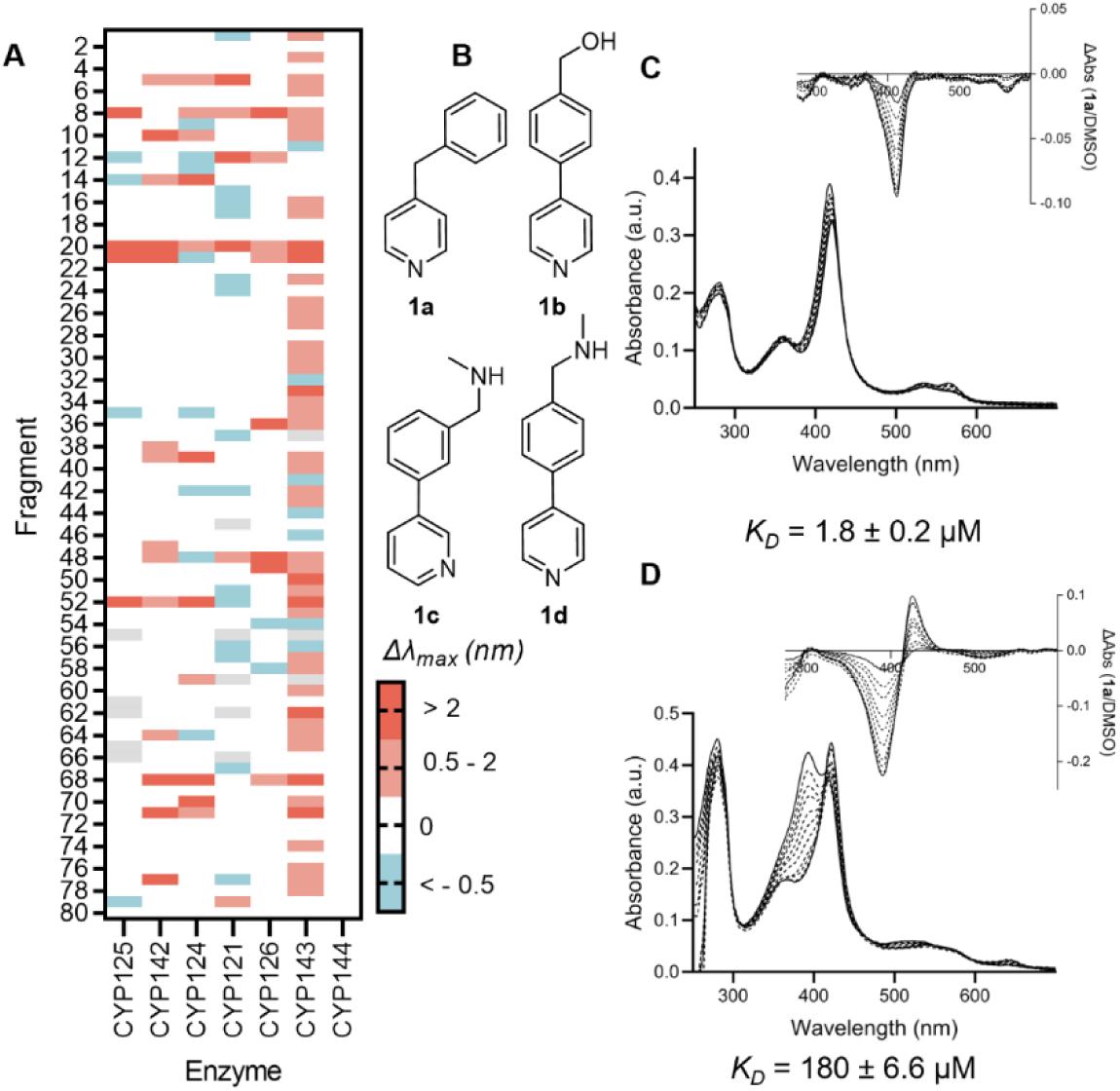
Identification of fragment hit 1a. **(A)** Heat map from screening focused fragment library (1 mM) against 7 *Mtb* P450s (4-6 µM) by UV-Vis spectroscopy. Color indicates the shift in the maximum wavelength of each enzyme’s absorbance spectrum (Δλ_max_, nm) relative to DMSO. Grey = not tested. **(B)** Structures of CYP125 hits. **(C, D)** Absorbance spectra from dose-response titration of fragment **1a** binding to CYP125 (C) or CYP142 (D) (each 5 μM). Insets – Difference spectra verse enzyme-DMSO complex. Data are mean of n = 3 titrations. K_D_ values were by fitted data to a hyperbolic model.

Only 5% of fragments in the focused library produced a red shift in the *λ_max_* of the CYP125 absorbance spectrum, which was surprisingly few compared to other *Mtb* and bacterial P450s that we have previously analyzed.^39,44,49^ All 4 of the CYP125 hit compounds contained a pyridine ring as the putative heme binding motif and a biphenyl or benzylpyridine structure (Fig. 1b). In contrast, 15% of fragments in the library produced a red shift in the *λ_max_* of the CYP142 spectrum, including 3 of the 4 fragments that were identified as hits for CYP125 (**1a, c, d**). The binding affinity (K_D_ value) of fragments **1a-d** to CYP125 and CYP142 was subsequently determined by optical titration,^50^ leading to the identification of benzylpyridine **1a** as the most potent compound (Fig. 1c, d). The low micromolar binding affinity of **1a** to both CYP125 (K_D_ = 180 ± 7 µM) and CYP142 (K_D_ = 1.8 ± 2.0 µM), good ligand efficiency (LE = 0.4 – 0.6), synthetically tractable chemical structure, and non-azole heme binding group, resulted in its selection for structural characterization and hit-to-lead optimization chemistry.

### Structural characterization of CYP142 bound to 1a

As CYP125 and CYP142 have a similar biochemical function,^25,28,34,35^ fragment-binding profile,^39^ and binding mode to **1a** (inferred from UV-vis spectroscopy) (Fig. 1c, d), we hypothesized that a structure of **1a** in complex with CYP142 might provide a suitable surrogate for CYP125, and help guide the development of dual CYP125/CYP142 inhibitors. A 1.7 Å X-ray crystal structure of **1a** in complex with CYP142 (Fig. 2a) was obtained by soaking fragment solutions into CYP142 crystals that were prepared by sitting-drop vapor diffusion. Interestingly, the structure revealed two molecules of **1a** bound per enzyme active site: one coordinated directly to the heme iron via the pyridine-N (**1a**-**i**), and the second located near the entrance of the active site channel (**1a**-**ii**). This CYP142-**1a** structure was aligned with that of CYP125 in complex with econazole (PDB 3IW2),^51^ as this was the only structure available of CYP125 bound to a type II, “inhibitor-like” ligand (Fig. 2b). Like **1a**, econazole increases the *λ_max_* of the CYP125 (and CYP142) optical spectrum and induces a shift EPR g-values consistent with direct co-ordination of the imidazole ring to ferric heme, as observed for CYP142-**1a-i**.^28^ However, as for CYP142-**1a-ii**, electron density for econazole in complex with CYP125 was only resolved near the near the entrance of the active site channel. These similarities made the CYP125-econazole structure suitable for comparison with CYP142-**1a**, and suggested that both heme co-ordination and hydrophobic interactions near the entrance of the P450 active site channel might constitute binding “hotspots”, which could be exploited to optimize the affinity of dual CYP125/142 inhibitors.^52,53^

**Figure 2.**
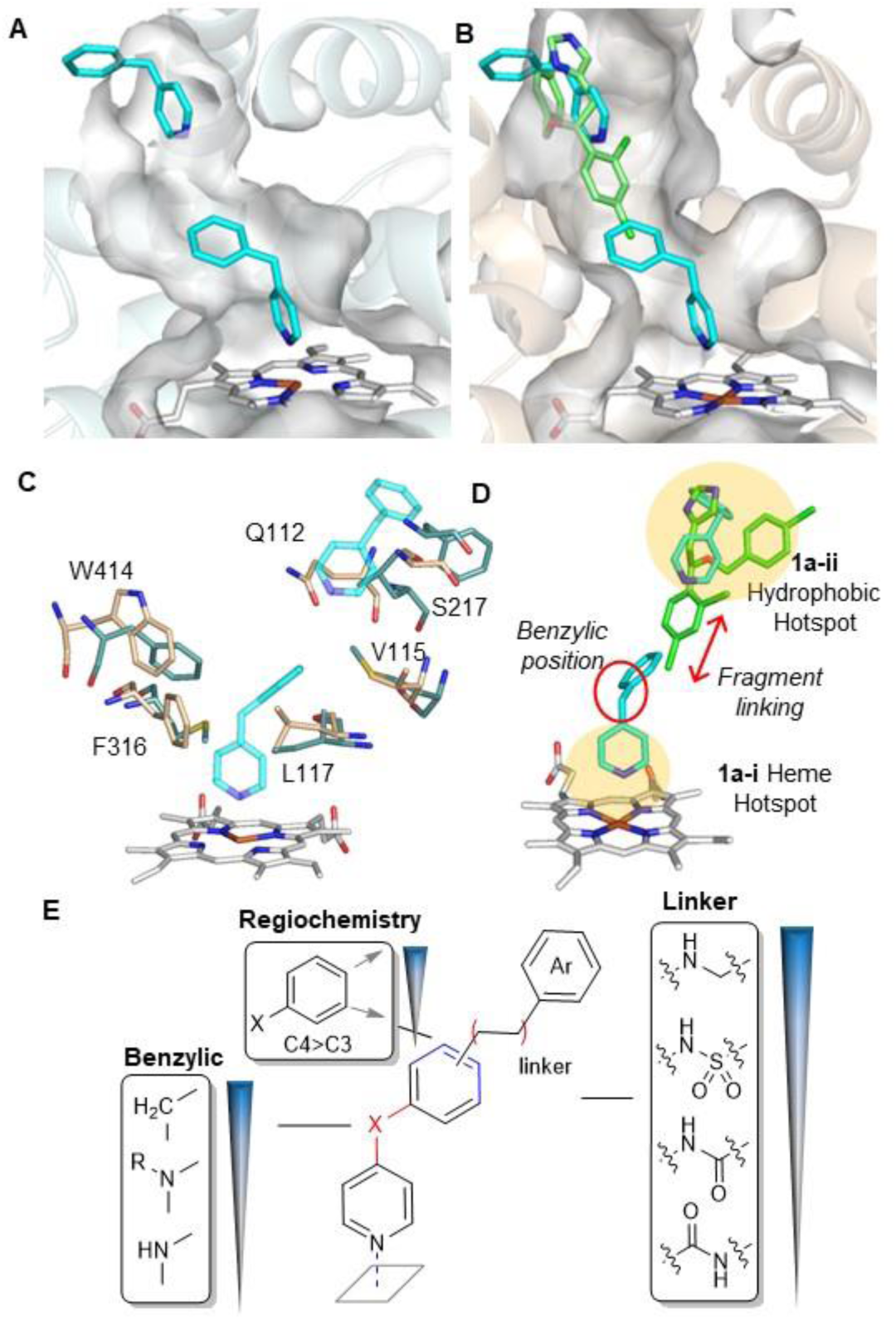
Structure-based design of dual CYP125/142 inhibitors. (A) X-ray crystal structure of CYP142-**1a** (cyan sticks) complex. Active site surface (grey) (PDB 8S53). (B) Overlay of CYP142-**1a** (cyan sticks) and CYP125-econazole (green sticks) (PDB 3IW2) structures. CYP125 cartoon (wheat) and active site surface (grey). (C) Key amino acids that differ between active site of CYP125 (wheat) and CYP142 (blue). Amino acid numbers refer to CYP125. Fragment **1a** (cyan). (D) Ligand design strategy, highlighting binding hotspots, and key motifs for SAR exploration. (E) CYP125 SARs established from screening a library of **1a** analogues that contained diverse functional groups at the benzylic position, and varied the functional group and substitution pattern of the **1a-i** – **1a-ii** “linker” (see Table S2).

The aligned structures indicated that the binding mode of **1a-i/ii** to CYP142 could be accommodated within the conformation of the econazole-bound CYP125 active site (Fig. 2b), and enabled the identification of key active site residues that differed between CYP125 and CYP142 (Fig 2c). The most notable of these included replacement of several aromatic residues near the CYP125 heme cofactor and **1a-i** benzylic-CH_2_ group (aka “*benzylic position*”), with smaller and/or aliphatic amino acids in CYP142 (e.g., ^125^F316 > ^142^M280, ^125^W414 > ^142^F380, and ^125^L117 > ^142^I76), and extensive differences in the F/G helices and B-C loop; including substitution of several residues located between the phenyl ring of **1a-i** and pyridine of **1a-ii** (aka *“linker”* region) (e.g., ^125^Q112 > ^142^L72, ^125^S217 > ^142^F179, ^125^V115 > ^142^Met74).^35^ As these variations could produce different SARs, the initial synthetic optimization of a dual CYP125/142 inhibitor focused on generating a library of analogues with diverse substituents at the “*benzylic*” and “*linker*” positions of the **1a** scaffold (Fig. 2d).

### Synthetic optimization of dual CYP125/142 inhibitors

A library of **1a** analogues was synthesized and screened by UV-vis spectroscopy to determine the SAR contributing to CYP125/142 binding affinity and selectivity. In the first iteration of compounds (**2a-h**), the effect of the “*benzylic*” functional group of **1a-i** was analyzed, and in the second iteration (**3a-g, 4a-i**), the functional group and substitution pattern of the “*linker”* used to join the **1a-i** phenyl ring with the **1a-ii** hydrophobic hotspot was varied (Fig 2d-e, Table S2). In brief, compounds containing different functional groups at the benzylic position were synthesized by either acid or copper-catalyzed arylation of aniline, phenol, or benzene sulfinic acid with a halopyridine (**2a-e**); or the condensation of 4-picoline with benzyladehyde (**2g-h**) (Scheme S1). Compounds synthesized to study the SAR of the “linker” were based on the scaffold of either **1a**, or the benzylic amine analogue **2a** (which showed improved binding to CYP142), and incorporated a wide range of functional groups at either C3- or C4-of the phenyl ring (Schemes S2, S3).

As observed in the original fragment screen (Fig. 1a, Table S1), CYP125 SAR were more stringent than CYP142 (Fig. 2e, Table S2). For example, only compounds with a tertiary amine (**2b**, **c**) or aliphatic group (**2g**, **h**) at the *benzylic* position caused a significant red shift in the CYP125 *λ_max_*, while CYP142 additionally bound to benzylic secondary amines (**2a**) or ethers (**2d**). Neither enzyme tolerated a polar group, such as a sulfone (**2e**) or carbonyl (**2f**), at the benzylic position. The addition of a “*linker*” substituent to the phenyl ring of either **1a-i**, or the 4-aminophenyl pyridine analogue **2a**, typically improved binding, however, SAR were again more stringent for CYP125 than CYP142. Amines (**3d**-**e, 4a**-**b**), amides (**3f**), or sulfonamides (**3g**) derivatives, substituted at C4 of the **1a-i** phenyl ring were preferred by CYP125, while CYP142 broadly tolerated amine, ether (**3a-b**), and ester (**4c-d**) substituents at either C3 or C4, but bound to sulfonamides comparatively weakly (**3g**). Both enzymes disfavored carboxylic acids (**4e-f**) or alcohol (**4g**) substituents, but bound more strongly to fragments containing a 3- or 4-bromo substituent (**4h**, **i**), highlighting the potential to significantly improve binding affinity by elaborating the **1a-i** scaffold to increase hydrophobic interactions with the **1a-ii** hotspot in the upper active site channel (Fig. 2d).

These SAR guided the synthesis of a second generation of **1a** analogues (**5a-p**), which were designed to optimize binding affinity for CYP125 and CYP142 by linking together the heme-binding and hydrophobic hotspots accommodated by **1a-i** and **1a-ii**, respectively. Each compound contained a methylene or amine at the *benzylic* position, and either an anilide, carboxamide, or sulfonamide *linker* of 3-4 bond lengths. The general synthesis of key compounds in this library is described in Scheme 1. In brief, Suzuki-Miyaura cross-coupling of pyridine boronic acid with a functionalized benzyl chloride (e.g., **7a-d**) afforded compounds with a methylene group at the benzylic position and either a carboxamide-linked aromatic group (**5e**, **5f**, **5h**, **5i**) or nitro substituent on the phenyl ring (**3c**). Reduction of the nitro group with tin(II) chloride yielded primary amines (**3d**, **3e**), which were subsequently coupled with carboxylic acids (**5a**, **5d**) or sulfonyl chlorides (**5j**, **5k**) using carbodiimide chemistry or base, respectively, or functionalized with aromatic substituents by reductive amination with a benzylaldehyde (**5l**, **5m**, **5p**) or acetophenone (**5o**). Compounds with a secondary amine at the benzylic position were synthesized by acid catalyzed coupling of 4-chloropyridine with a functionalized aniline to yield **4c** or **6b**, followed by alkylation with methyl iodide to yield the tertiary amine **8a**. Ester and nitro groups were hydrolyzed or reduced, respectively (**4b**, **4e**, **8b**), and then used to synthesize anilide (**5b**, **5c**), carboxamide (**5g**), or benzylamine (**5n**)-linked aromatic substituents.

The binding affinity (K_D_ values) of the resulting compounds (**2a**, **3f**, **3g**, **5a-p**) to CYP125 and CYP142 validated the SARs established during the initial iterations of fragment optimization (Table 1). For example, replacing the benzylic methylene group with a secondary or tertiary amine translated to an approximate 20-fold loss in CYP125 binding affinity (e.g., **5a** vs **5c**, **5e** vs **5g**, **5m** vs **5n**), while varying the C4/C3-substitution pattern on the **1a-i** phenyl ring resulted in 10-100-fold difference in K_D_ value (e.g., **5e** vs **5f**, **5h** vs **5i**, **5l** vs **5m**). In contrast, the binding affinity of most compounds to CYP142 was similar (K_D_ ∼ 1 µM). However, a 20-fold improvement was achieved by replacing either the amide (e.g. **5a**) or sulfonamide (e.g. **5j**) linker with a methyl amine (e.g., **5m**). Combining the SAR favored by CYP125 and CYP142 yielded a potent dual inhibitor **5m**, which had K_D_ values of 40-160 nM for both enzymes, and good ligand efficiency (LE) (> 0.4) due to the high group efficiency (GE) of the benzylamine (GE = 0.20-0.50).

**Table 1.**
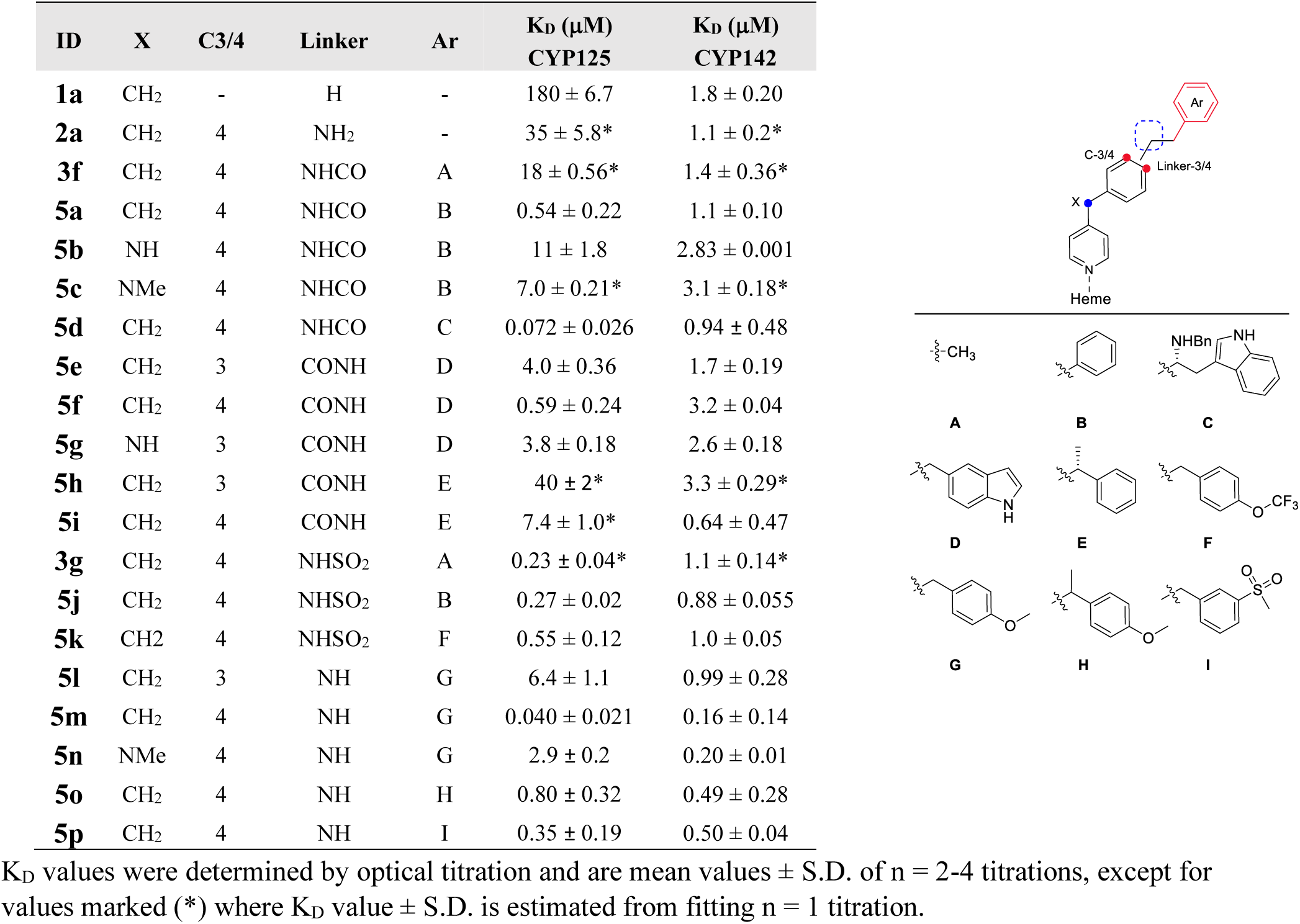
Structure and binding affinity of dual CYP125-142 inhibitors. K_D_ values were determined by optical titration and are mean values ± S.D. of n = 2-4 titrations, except for values marked (*) where K_D_ value ± S.D. is estimated from fitting n = 1 titration.

| ID | X | C3/4 | Linker | Ar | K<sub>D</sub> (μM)<br>CYP125 | K<sub>D</sub> (μM)<br>CYP142 |
| --- | --- | --- | --- | --- | --- | --- |
| **1a** | CH<sub>2</sub> | - | H | - | 180 ± 6.7 | 1.8 ± 0.20 |
| **2a** | CH<sub>2</sub> | 4 | NH<sub>2</sub> | - | 35 ± 5.8* | 1.1 ± 0.2* |
| **3f** | CH<sub>2</sub> | 4 | NHCO | A | 18 ± 0.56* | 1.4 ± 0.36* |
| **5a** | CH<sub>2</sub> | 4 | NHCO | B | 0.54 ± 0.22 | 1.1 ± 0.10 |
| **5b** | NH | 4 | NHCO | B | 11 ± 1.8 | 2.83 ± 0.001 |
| **5c** | NMe | 4 | NHCO | B | 7.0 ± 0.21* | 3.1 ± 0.18* |
| **5d** | CH<sub>2</sub> | 4 | NHCO | C | 0.072 ± 0.026 | 0.94 ± 0.48 |
| **5e** | CH<sub>2</sub> | 3 | CONH | D | 4.0 ± 0.36 | 1.7 ± 0.19 |
| **5f** | CH<sub>2</sub> | 4 | CONH | D | 0.59 ± 0.24 | 3.2 ± 0.04 |
| **5g** | NH | 3 | CONH | D | 3.8 ± 0.18 | 2.6 ± 0.18 |
| **5h** | CH<sub>2</sub> | 3 | CONH | E | 40 ± 2* | 3.3 ± 0.29* |
| **5i** | CH<sub>2</sub> | 4 | CONH | E | 7.4 ± 1.0* | 0.64 ± 0.47 |
| **3g** | CH<sub>2</sub> | 4 | NHSO<sub>2</sub> | A | 0.23 ± 0.04* | 1.1 ± 0.14* |
| **5j** | CH<sub>2</sub> | 4 | NHSO<sub>2</sub> | B | 0.27 ± 0.02 | 0.88 ± 0.055 |
| **5k** | CH<sub>2</sub> | 4 | NHSO<sub>2</sub> | F | 0.55 ± 0.12 | 1.0 ± 0.05 |
| **5l** | CH<sub>2</sub> | 3 | NH | G | 6.4 ± 1.1 | 0.99 ± 0.28 |
| **5m** | CH<sub>2</sub> | 4 | NH | G | 0.040 ± 0.021 | 0.16 ± 0.14 |
| **5n** | NMe | 4 | NH | G | 2.9 ± 0.2 | 0.20 ± 0.01 |
| **5o** | CH<sub>2</sub> | 4 | NH | H | 0.80 ± 0.32 | 0.49 ± 0.28 |
| **5p** | CH<sub>2</sub> | 4 | NH | I | 0.35 ± 0.19 | 0.50 ± 0.04 |

### Structural characterization of dual CYP125/142 inhibitors

A combination of improvements made to CYP125 expression and crystallization conditions during the synthetic optimization of **1a**, in addition to the tight binding affinity of the dual CYP125/142 inhibitors, enabled us to obtain high resolution X-ray crystal structures of compounds **5j** and **5m** in complex with CYP125 (Fig. 3a, 3c, 3h) and CYP142 (Fig. 3b, 3d, 3h), and compound **5g** in complex with CYP125 (Fig. S1, S2 and Table S3). In all 4 structures, the pyridine-nitrogen of the inhibitor directly coordinated to the P450 heme iron, consistent with their type II optical spectra and the binding mode of **1a-i** to CYP142 (Fig 2a). The CYP125-**5m** (Fig. 3a) and CYP142-**5m** (Fig. 3b) structures also illustrate that the 4-methoxybenzylamine substituent accurately recapitulates the binding mode of **1a-ii** in the hydrophobic hotspot (Fig. 3e), validating the fragment-linking strategy used to optimize inhibitor binding affinity. In contrast, the conformation of the sulfonamide linker in compound **5j** directs the phenyl substituent away from the hydrophobic hotspot in both CYP125 (Fig. 3c) and CYP142 structures (Fig. 3d), introducing disfavorable steric interactions with Met74 in CYP142 (Fig. 3f). This conformation might account for the weaker binding affinity of compounds that contain a sulfonamide (e.g. **3g**, **5k**) or amide linker (e.g., **5a-d**) relative to their benzylamine analogues (e.g., **5l-p**), and is consistent with the comparatively weak affinity of CYP142 to fragments containing a sulfonamide substituent, which was noted in the original SAR screen (e.g., **3g**, Table S2).

**Figure 3.**
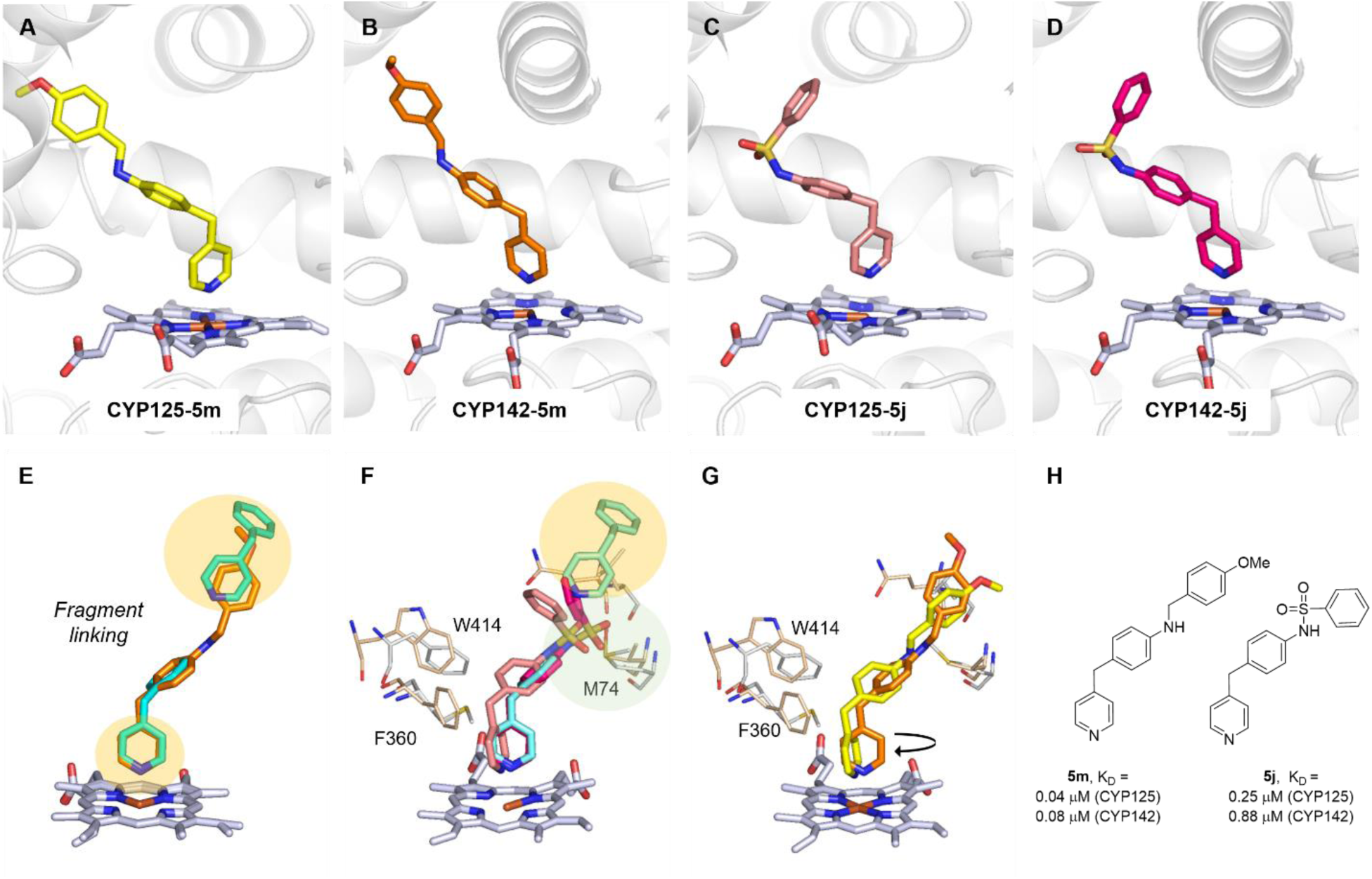
Structural characterization of CYP125-142 inhibitors. X-ray crystal structures of CYP125 (A, C) or CYP142 (B, D) in complex with **5m** (A, B) or **5j** (C, D). Heme cofactor is shown as grey sticks and protein secondar structure as grey cartoon. (E) Overlaid structures of CYP142**-5m** (orange) and CYP142-**1a** (blue), highlighting heme and hydrophobic hotspots (yellow). (F) Overlaid structure of CYP125-**5j** (salmon), CYP142-**5j** (magenta), and CYP142-**1a** (blue), highlighting key active site residues as wheat (CYP125) or grey (CYP142) sticks. Hydrophobic hotspot (yellow) and sulfonamide clash (green) show in colored spheres. (G) Overlaid structures of CYP125-5m (yellow) and CYP142-5m (orange). Arrows indicate rotated orientation of CYP125-bound pyridine and proximity of benzylic position to aromatic active site residues. Red lines indicate requirement of C4 substitution pattern for binding to CYP125. (H) Chemical structure and K_D_ values of compound **5m** and **5j** binding to CYP125 or CYP142, as determined by optical titration.

In all CYP125 co-crystal structures, the pyridine ring of the inhibitor was rotated 90 degrees relative to that observed for the same inhibitor in complex with CYP142 (Fig. 3g), and the orientation of the **1a-i** pyridine ring in the original CYP142-**1a** structure (Fig. 2a). This orientation better accommodates the aromatic residues Phe360 and Trp414 in the CYP125 active site and likely contributes to the sensitivity of CYP125 to bulky or polar substituents at the *benzylic* position of **1a-i** (Fig. 2e, Table 1, Table S2). Rotation of the pyridine ring could also account for the preference of CYP125 for a C4 substitution pattern on the **1a-i** phenyl ring, as unlike CYP142, only elaboration from C4 provides direct alignment with the hydrophobic hotspot.

The structure of compound **5g** in complex for CYP125 (Fig. S1) revealed a surprising “substrate-like” shift in the enzyme active site, whereby amino acids in the F- and I-helices move inwards to attain a hybrid orientation that is midway between the previously reported substrate-(**cholestenone**) and inhibitor-(**econazole**) bound CYP125 complexes.^31^ Furthermore, Glu271 extends across the heme cofactor to hydrogen bond with the benzylic amine of **5g**. This unusual orientation could account for the weak type II optical spectra generated by **5g**, and other compounds containing an amine at the *benzylic* position and/or C3 phenyl substitution pattern.

### Inhibition of CYP125/142 catalytic activity *in vitro*

The ability of the elaborated **1a** analogues to inhibit CYP125/142 catalytic activity was assessed *in vitro* using an LC-MS-based substrate turnover assay to monitor the conversion of cholest-4-en-3-one (Fig 4a). Experiments were performed as previously described,^28,35^ using recombinantly expressed and purified CYP125 or CYP142, and an exogenous electron transport chain consisting of spinach ferrodoxin/ferrodoxin reductase coupled to a glucose-6-phosphate/glucose-6-phosphate dehydrogenase-NADP(H) regenerating system. All compounds that were tested in this assay inhibited CYP125 catalytic activity at concentrations that correlated with the K_D_ values determined from optical titrations (R^2^ = 0.82) (Fig 4b, c), and 4 compounds (**5d**, **5j**, **5k**, **5m**) were calculated to have inhibition constants (K_I_ values) less than 1 µM (Fig. 4b, d, Table S4). A subset of the most potent CYP125 inhibitors was subsequently tested against CYP142 (K_I_ values between 0.05 – 1.1 μM) and found to correlate with the relative potency against CYP125 (R^2^ = 0.89) (Fig. 4b-d). The most potent dual CYP125/142 inhibitor **5m** (K_I_ = 0.10 μM (CYP125), 0.05 μM (CYP142)), was subsequently selected as the lead candidate for biological profiling. No obvious oxidation of the CYP125/142 inhibitors themselves could be detected in the biochemical assays. However, a detailed analysis of all reaction products was not performed.

**Figure 4.**
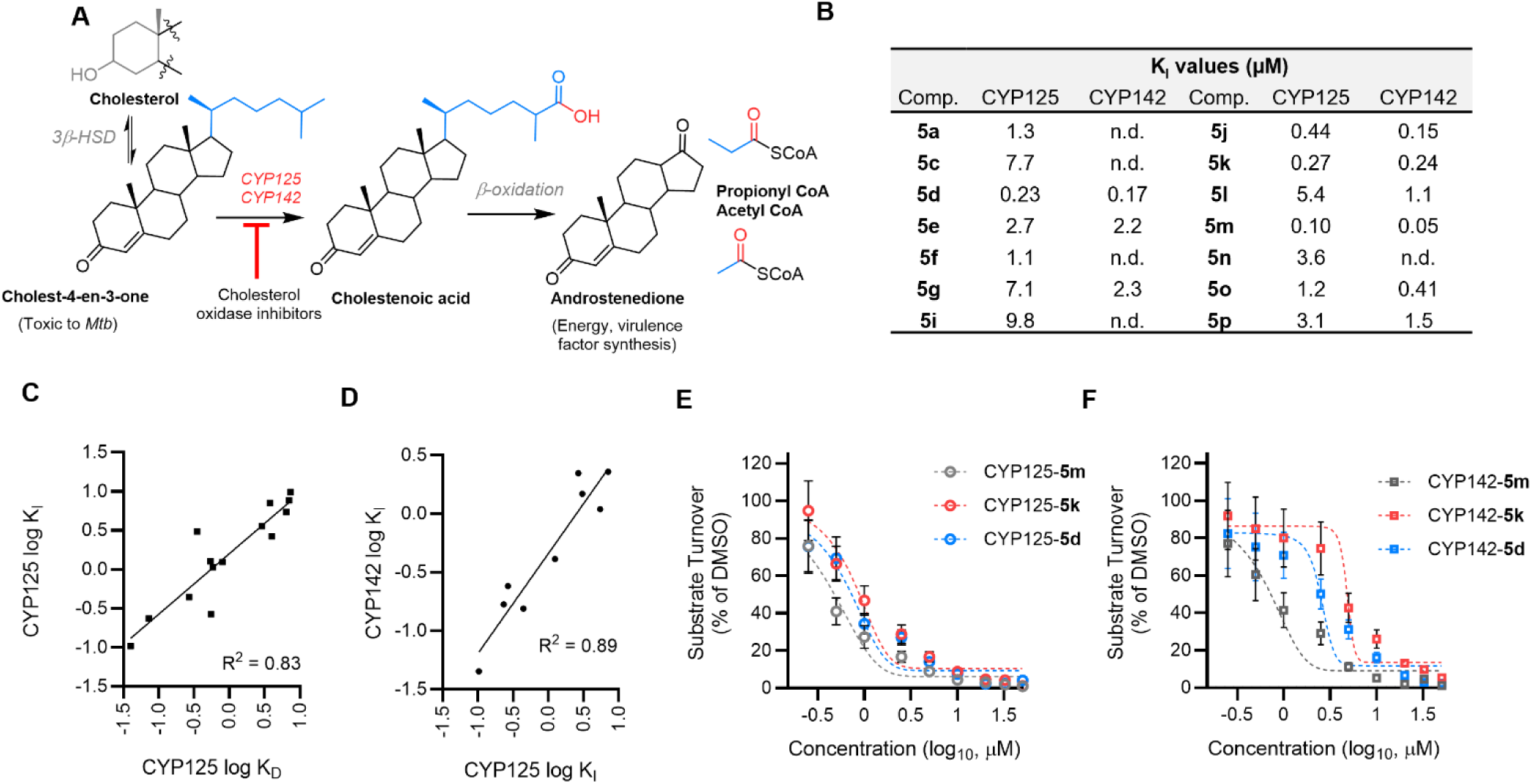
Inhibition of CYP125-142 cholestenone metabolism *in vitro*. (A) *Mtb* cholesterol metabolism: CYP125 and CYP142 catalyze C27-oxidation of cholest-4-en-3-one to yield cholestenoic acid, which is subsequently degraded to androstendione, acetyl CoA, and propionyl CoA. 3β-HSD - 3β-hydroxysteroid dehydrogenase; (B) Inhibition equilibrium constants (K_I_) values for the turnover of cholest-4-en3-one (5 μM) by CYP125 (0.5 μM) or CYP142 (1 μM). K_I_ values estimated by Cheng-Prusoff equation, cholest-4-3-one K_m_ CYP125 (2.1 μM), K_m_ CYP142 (0.36 μM); (C) Binding affinity (K_D_) and inhibition constants (K_I_) of **1a** analogues for CYP125 correlates (R^2^ = 0.83, P <0.0001); (D) Inhibition (K_I_ values) of CYP125 and CYP142 by **1a** analogues correlates (R^2^ = 0.89, P = 0.001); (E, F) Inhibition of CYP125 (0.5 μM) (E) or CYP142 (1 μM) (F) catalyzed turnover of cholest-4-en-3-one (5 μM) by **5d**, **5k**, and **5m**. Data are mean ± SD of n=3 replicates.

### Antimicrobial activity of dual CYP125/142 inhibitors against extracellular *Mtb*

We initially assessed the antimicrobial activity of the CYP125/142 inhibitors against extracellular *Mtb* (H37Rv) that was cultured in media containing cholesterol as the sole source of carbon, as the genetic disruption of CYP125, or CYP125 and CYP142, inhibits the growth of *Mtb* under these conditions.^21,25,29,35^ The concentration of compound required to completely inhibit *Mtb* growth (MIC_99_) was calculated 2 weeks post-compound treatment from the reduction of resazurin (MABA)^54^ relative to DMSO-treated controls (Table 2, Table S5). The most potent dual CYP125/142 inhibitor **5m**, was found to also have the strongest antimicrobial activity (MIC_99_ = 1.5 µM, ∼ 0.46 µg/mL), and several other compounds also inhibited *Mtb* growth with modest MIC_99_ values of between 12.5 – 25 µM, including amides **5d**, **5e**, **5g**, sulfonamide **5k**, and benzylamines structurally related to **5m** (**5l**, **5o**, **5p**).

**Table 2.**
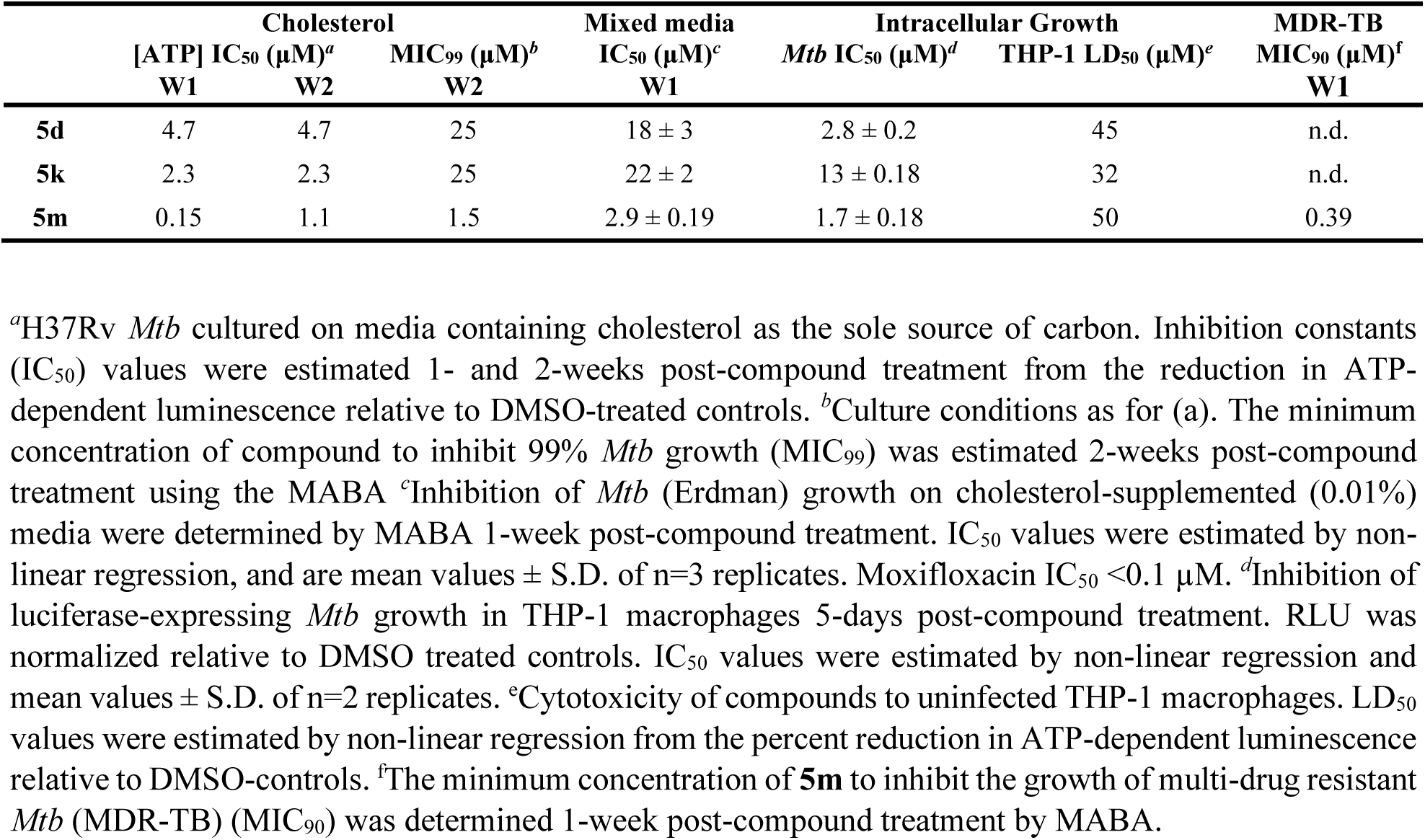
Antitubercular activity of CYP125/142 inhibitors.

Encouraged by these results, we repeated this experiment and used an ATP luminescence assay to provide a more direct measure of the effect of the CYP125/142 inhibitors on *Mtb* metabolism over the 2-week treatment period (Table 2, Table S5). These independent experiments confirmed that **5m** (IC_50_ = 0.15 µM), and benzylamines **5l**, **5o**, **5p** (IC_50_ values 0.15 - 1.5 µM) potently depleted intracellular ATP concentrations 1-week post-compound treatment (Fig. 5a, Table S5). Several other compounds including **5d**, **5g**, and **5k**, also had IC_50_ values < 5 µM, which is consistent with the reliance of *Mtb* on cholesterol metabolites to drive ATP generation under these growth conditions,^19^ and parallels the activity of other compounds with target *Mtb* metabolism.^8,9^ The IC_50_ values of all compounds increased between 1-week and 2-week measurements, (e.g., 2-week IC_50_ **5m** = 1.2 µM), suggesting that MIC_99_ values recorded in initial experiments might improve with repeated compound dosing, or measurement 1-week post-compound treatment.

**Figure 5.**
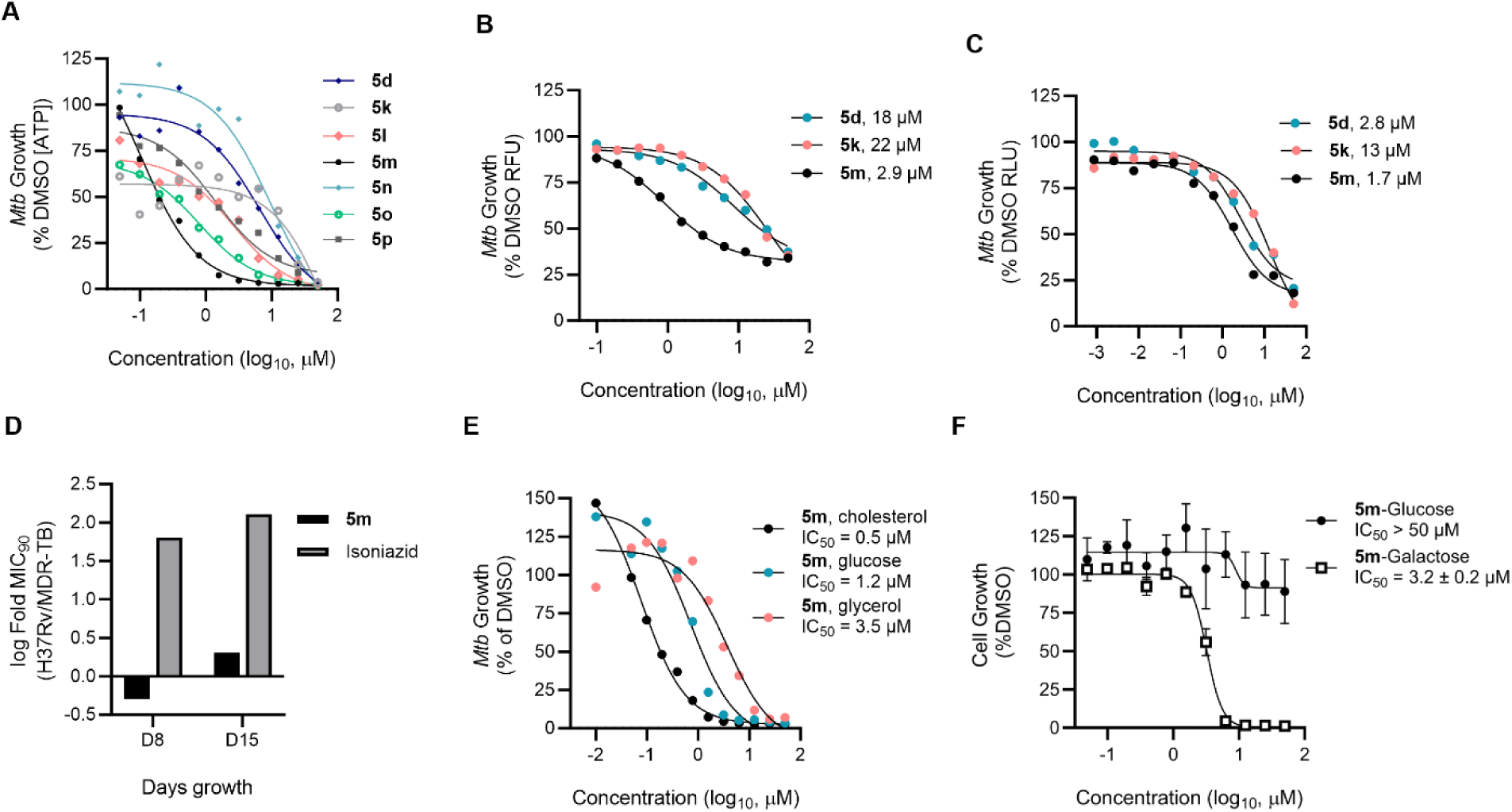
Antitubercular activity of CYP125/142 inhibitors. (A) CYP125/142 inhibitors deplete intracellular ATP when *Mtb* (H37Rv) is cultured on cholesterol as the sole carbon source. ATP concentration was assessed 1-week post-compound treatment and RLU was normalized as a percent of the DMSO-treated control. (B) Inhibition of extracellular *Mtb* (Erdman) growth on media supplemented with 0.01% cholesterol, determined by MABA 1-week post-compound treatment. RFU was normalized as a percent of DMSO-treated controls, and data are mean values ± S.D. of n=3 replicates. (C) Inhibition of *Mtb* (H37Rv:pATB45luc) growth in THP-1 macrophages, quantified from RLU 5-days post-compound treatment. Data are represented as a percent of the DMSO treated control are mean values ± S.D. of n=2 replicates. (D) Lead compound **5m** retains inhibitory activity against multi-drug resistant *Mtb* (MDR-TB). MIC_90_ (μM) values of **5m** and isoniazid were determined against extracellular H37Rv and MDR *Mtb* cultured on standard media 1- and 2-weeks post-compound treatment, and used to calculate log-fold change (MIC_90_ H37Rv/MIC_90_ MDR-TB). (E) Comparison of **5m** potency against extracellular *Mtb* (H37Rv) cultured on media containing either cholesterol, glucose, or glycerol as the sole source of carbon. Inhibition of *Mtb* growth relative to DMSO-treated controls was determined from ATP concentration (cholesterol and glucose) or MABA (glycerol) after 7- or 10-days post-compound treatment, respectively. (F) Selective cytotoxicity of **5m** against HepG2 cells cultured on galactose-containing media. Growth inhibition was determined from ATP concentration (RLU). Data are represented as a percent of the DMSO-treated controls, and are mean values ± S.D. of n=2 replicates.

As the accumulation of the toxic CYP125/142 substrate cholestenone has been shown to inhibit the growth of ΔCyp125/142 *Mtb*,^21^ we also assessed the antimicrobial activity of a representative subset of the CYP125/142 inhibitors (**5d**, **5k**, **5m**) against bacteria that were cultured on standard media supplemented with low concentration cholesterol (0.01% w/v) (Table 2, Fig. 5b). For these assays, inhibition of *Mtb* (Erdman) growth was quantified ∼1-week post-compound treatment using the MABA. All compounds retained inhibitory activity, with compound **5m** estimated to have an IC_50_ value of 2.9 µM. These data demonstrate that the antimicrobial activity of the CYP125/142 inhibitors could extend to an environment with more diverse nutrient availability, as found *in vivo*.

### Antimicrobial activity of CYP125/142 against *Mtb* in human macrophages

Encouraged by the activity of the CYP125/142 inhibitors against extracellular *Mtb*, we proceeded to assess the ability of a representative subset of compounds to inhibit the growth of luciferase expressing *Mtb* (H37Rv pATB45luc) in human macrophage-like THP-1 cells.^55^ *Mtb* growth inhibition was determined 5-days post-compound treatment (infection) from the reduction in luminescence signal intensity relative to DMSO-treated controls. The potential cytotoxicity of the CYP125/142 inhibitors to uninfected THP-1 macrophages was assessed in parallel experiments using an ATP glow assay. Both benzylamine **5m** (IC_50_ = 1.7 µM) and amide **5d** (IC_50_ = 2.8 µM) potently inhibited the growth of intracellular *Mtb*, while sulfonamide **5k** was considerably less active (13 µM) (Fig. 5c, Table 2). Compound **5m** also showed good selectivity over mammalian cytotoxicity, with no effect on THP-1 cell viability at concentrations up to 50µM. These results illustrate that **5m**, and related benzylamine compounds, have suitable properties for use in cell-based assays, which could help better understand the role of CYP125/142 in *Mtb* pathology.

### CYP125/142 inhibitors are active against MDR-TB

Finally, we assessed the ability of benzylamine **5m** to inhibit the growth of a MDR strain of *Mtb* (K26b00MR 113), which is insensitive to the first line anti-TB drugs isoniazid and rifampicin.^56^ The antimicrobial activity of **5m** and isoniazid were assessed in parallel against drug sensitive H37Rv *Mtb* and MDR-TB by MABA, 1- and 2-weeks post-compound treatment. While **5m** retained a similar MIC_90_ value against both strains of *Mtb* (H37Rv = 0.78 µM, MDR-TB = 0.39 µM), the IC_50_ value of isoniazid was > 50-150-fold higher against MDR-TB (Table 2, Fig. 5d, Table S6). Considering the increasing prevalence of MDR-TB across the globe, and limited pipeline of novel anti-TB drugs, the potency of **5m** in these experiments supports further exploration of the benzylpyridine chemotype as anti-TB agents.

### Biological mechanism and safety profiling

Confident that **5m** inhibits the growth of *Mtb* on cholesterol, we subsequently explored the scope of the compound’s antimicrobial activity by testing a subset of the CYP125/142 inhibitors against extracellular *Mtb* cultured on other defined carbon sources. These experiments revealed a notable 8-fold decrease in the potency of **5m** against *Mtb* grown on glucose (IC_50_ = 1.2 µM) compared to cholesterol (IC_50_ = 0.15 µM), and > 20-fold decrease in potency against *Mtb* grown on minimal media supplemented with glycerol (IC_50_ = 3.5 µM) (Fig. 5e, Table S7). Similarly, there was a 2 - 16-fold increase in the IC_50_ value of benzylamines **5l**, **5o** and **5p**, and sulfonamide **5k** against *Mtb* grown on glucose verse cholesterol, while in contrast, compound **5d** had a similar IC_50_ value regardless of media composition (Table S7).

This carbon-dependent trend in the antimicrobial activity of **5m** is consistent with a mechanism of action that is, at least in part, dependent on inhibiting cholesterol metabolism. However, these results also indicated that the compound may have an additional mechanism(s) of action. A panel of reporter assays were subsequently used to detect whether the compound interacted with biological pathways targeted by existing TB drugs (Fig. S3). As treatment of *Mtb* with compound **5m** did not induce the upregulation of *iniB*, or *recA* and *radA* reporters, our results imply that the compound is unlikely to inhibit *Mtb* cell wall synthesis, or induce DNA damage, respectively.^57^ However, data from several other reporter assays were inconclusive due to the presence of BSA (Fig. S4) and further mechanistic characterization should be performed in future.

The good selectivity of **5m** for bacterial cytotoxicity verse mammalian cells that was observed when the compound was tested against THP-1 macrophages (Table 2), was independently verified in experiments using HepG2 cells cultured on standard (glucose-containing) media (LD_50_ >50 µM). However, notably, **5m** showed significant cytotoxicity against HepG2 cells cultured on media containing galactose (LD_50_ = 3.2 µM), indicating that the compound may inhibit oxidative metabolism (Fig. 5f).^58^

Finally, to provide insight into whether CYP125/142 inhibitors based on the benzylpyridine scaffold might cause drug-drug interactions or off-target activity when used *in vivo*, **5m** was screened against a panel of human drug metabolizing P450s (Table S8). As several P450 isoforms were inhibited as concentrations < 5 µM, the compound has potential to cause drug-drug interactions and further optimization of the benzyl pyridine scaffold to improve CYP125/142 selectivity would be required for in vivo applications. Despite this, the good activity of **5m** against intracellular *Mtb* and MDR-TB, and low mammalian cytotoxicity, should make the compound a useful tool to study CYP125/142 in cell-based assays, and promising that further optimization could yield a novel class of anti-TB compounds.

### Discussion

The reliance of intracellular *Mtb* on host-derived cholesterol for long-term survival and virulence, makes cholesterol metabolism a compelling biological target for the development of novel antibiotics. ^19–22,26,27,59^ Despite this, few compounds have been developed to specifically inhibit key enzymes involved in *Mtb* cholesterol metabolism.^60–62^ Here, we have described an efficient fragment- and structure-guided approach to develop small molecule inhibitors of the P450 enzymes CYP125 and CYP142, which catalyze the first committed step of cholesterol degradation in *Mtb*.^28–31,34,35^ The lead compounds developed in this study have activity against both intracellular *Mtb* and MDR-TB, low toxicity to human macrophages, and drug-like chemical properties, which should make them useful tools to study *Mtb* sterol metabolism, and are promising step towards the development of novel drugs that could help combat the global TB pandemic.

Our earlier attempts to develop CYP125 inhibitors were hindered by a lack of structural data to guide hit-to-lead optimization,^44^ and further complicated by the discovery that CYP142 – an enzyme with low sequence or structural similarity to CYP125 – could rescue the growth of *ΔCyp125 Mtb* on cholesterol.^25,34,63^ To overcome these technical and biological challenges, we leveraged fragmented screening by UV-vis spectroscopy to efficiently sample chemical space,^64^ characterize the SARs shared by CYP125 and CYP142, and identify a chemical scaffold with suitable drug-like properties that could be used for the development of a dual CYP125/142 inhibitor. By designing a tailored heme-binding fragment library and employing UV-vis spectroscopy, instead of more commonly used biophysical screening techniques,^65^ we were able to rapidly identify fragments that not only bound to CYP125 and CYP142, but also functionally stabilized the low-spin or inactive state of the enzymes (Fig 1a-d).^48^ UV-vis spectroscopy also provided detailed understanding of the binding site and orientation of the fragment hits, which helped guide hit-to-lead optimization chemistry, and enabled us to use the co-crystal structure of **1a** in complex with CYP142 as a structural proxy for CYP125 (Fig 2b, d).

The significant improvement in binding affinity that was achieved through synthetically linking together fragment **1a-i** and **1a-ii** to yield compound **5m** (K_D_ **1a**/**5m**: CYP125 > 1000-fold, CYP142 > 10-fold), illustrates how fragment screening can be used to identify energetic hotspots that may contribute disproportionately to binding affinity.^66^ Furthermore, as observed previously, our data highlights the importance of optimizing the properties of the chemical linker to ensure that the original fragments can maintain an optimal binding orientation (Fig. 3a-d, and Table 1).^67,68^ The lead compound development through this fragment linking approach (**5m**) binds to both CYP125 and CYP142 with comparable affinity to the enzyme’s endogenous substrates (K_D_ ∼ 100 nM),^34,63^ has excellent ligand efficiency (LE > 0.4), and potently inhibits the enzyme’s catalytic activity *in vitro* (K_I_ ∼ 0.05 – 0.10 µM).

Throughout the fragment screening and inhibitor optimization campaign we noted that structure-activity relationships (SAR) for binding to CYP125 were significantly more sensitive than CYP142. For example, varying the substitution pattern or chemistry of the **1a-i**-**1a-ii** linker resulted in up to 100-fold difference in CYP125 binding affinity, while CYP142 K_D_ values varied <10-fold (Table 1, Fig. 2e). These results reflect the structural differences in upper active site of CYP125 and CYP142,^63^ and support hypothesis that CYP142 may have evolved to metabolize a more diverse pool of sterol substrates than CYP125.^69^ While CYP125 is encoded within the conserved *igr* operon, and has been functionally characterized as a cholesterol oxidase in multiple species of actinobacteria,^27–30^ CYP142 is encoded within a cluster of lipid metabolizing genes and shares greater structural similarly with *Mtb* CYP124;^63^ an enzyme that is thought to primarily oxidize fatty acids and vitamin D.^70,71^ Building on the data reported here, could help develop isoform-selective CYP125 or CYP142 inhibitors that would facilitate research into the enzyme’s independent roles in *Mtb* sterol metabolism. For example, the significantly larger proportion of CYP125-**5m** binding affinity that can be attributed to forming interactions with the **1a-ii** hotspot (ΔΔG [**5m**-**1a**] = 0.5) compared to CYP142 (ΔΔG [**5m**-**1a**] = 0.2), suggests that hydrophobic interactions near the entrance of the P450 active site contribute disproportionality to CYP125 ligand recognition. In contrast, CYP142 binding affinity to the **1a** compound series appears to be more strongly driven by pyridine-heme co-ordination. Removing or attenuating the potency of heme binding pyridine might favor CYP125 selectivity or could be used to reduce off-target interactions with other P450s, while modifying the linker to exploit differences in the distal active site of CYP125/142 might yield CYP142-selective compounds. The ligands and SAR reported here may also help guide the development of inhibitors for the human cholesterol oxidases CYP27A1 and CYP46A1, both of which currently lack high chemical probes. As CYP27A1 and CYP46A1 play important roles in bile acid biosynthesis and the elimination of cholesterol from the brain, respectively,^72–74^ profiling the activity of the dual CYP125/142 inhibitors against human CYP27A1 and CYP46A1, as well as a broad spectrum of other human P450s, will be an important consideration when assessing their further optimization as anti-TB compounds.

The antimicrobial activity of the dual CYP125/142 inhibitors against *Mtb* grown on cholesterol (Fig. 5a, Table 1, Table S4), or cholesterol-supplemented rich media (Fig. 5b, Table S5), is consistent with previous studies in which either *Cyp125*, or *Cyp125* and *Cyp142*, were genetically disrupted.^21,25,29,31^ In addition, the weaker activity of **5m**, and related benzylamine compounds, against *Mtb* that was cultured on either glucose or glycerol as a sole source of carbon supports a mechanism of action that is, at least in part, dependent on cholesterol utilization (Fig. 5e, Table S7). In contrast, the potency of control compounds (e.g., 4-aminosalicylic acid, isoniazid) was similar regardless of media composition. Although nutrient availability can alter *Mtb* growth rate, we did not observe any intrinsic differences in fitness across experimental conditions, and the MIC values of control compounds (e.g., 4-aminosalicylic acid, isoniazid) was similar regardless of carbon source (Table S5 & S7).

Despite this, the ability of **5m** to inhibit the growth of extracellular *Mtb* in the absence of cholesterol, suggests that either CYP125 and CYP142 have important, uncharacterized physiological functions, or that **5m**, and related analogues, have a secondary mechanism of action. Preliminary biological profiling indicated that **5m** is unlikely to induce DNA damage or inhibit *Mtb* cell wall synthesis, which are the mechanisms of some existing first line TB drugs (e.g. isoniazid, fluoroquinolones), however, other reporter assays were inconclusive (Fig. S3). The ability for both bedaquiline and **5m** to potently decrease intracellular ATP, and the enhanced potency of both compounds against *Mtb* grown on lipids compared to standard glucose or glycerol media, suggests an overlap in their mechanism(s) of action at the level of oxidative phosphorylation.^7,75^ However, further mechanistic characterization is required. Despite this, the potent activity of **5m** against both drug susceptible and MDR-*Mtb* cultured under a variety of conditions provides promise that further optimization of the benzylpyridine scaffold could yield compounds that retain antitubercular activity *in vivo*, where *Mtb* can access more heterogenous carbon sources.

*Mtb’s* unique metabolic adaptions to survive in human macrophages contributes to the bacteria’s reduced sensitivity to first-line TB drugs, and the need to identify compounds which specifically have activity against intracellular *Mtb*.^7,76^ As the utilization of cholesterol is required to establish a long-term, chronic infection, and is one of the primary nutrients available to non-replicating *Mtb,*^3,19,21,77^ we anticipate that drugs targeting CYP125/142 could help to specifically address recalcitrant bacterial populations. Our study demonstrates that compound **5m** inhibits the growth of *Mtb* in recently infected human macrophage-like cell lines, and we anticipate that, like other drugs targeting *Mtb* metabolism, **5m** may also have activity against dormant *Mtb*.

The low cytotoxicity of compound **5m** to both THP-1 macrophages and HepG2 cells cultured on glucose is consistent with evidence that THP-1 cells are primarily glycolytic,^78^ and should enable the compound to be used as a chemical tool to help study the role of CYP125/142 during infection. In addition, as shifting macrophage metabolism towards aerobic glycolysis correlates with a more effective immune response,^79^ **5m** might synergistically decrease *Mtb* fitness and have a beneficial immunomodulatory effect. For example, hydroxycholesterol metabolites, such as those synthesized by CYP125/142, have been reported to polarize macrophages towards a more tolerogenic M2 phenotype.^80,81^ As such, it would be intriguing in future studies to analyze whether CYP125/142 inhibition alters macrophage cytokine profiles. Furthermore, as carbon liberated from cholesterol metabolism is used to synthesize virulence-associated lipids such as phthiocerol dimycocerosate (PDIM),^19,20^ future studies should evaluate the effect of CYP125/142 inhibition on *Mtb* cell wall integrity and immunogenicity.

In contrast, compound **5m** was selectively cytotoxic to HepG2 cells cultured on galactose media, suggesting that the compound might inhibit mammalian mitochondrial function.^58^ Interestingly, the FDA-approved *Mtb* ATP-synthase inhibitor bedaquiline has also recently been shown to inhibited the growth of tumor-initiating cancer stem cells through interfering with mammalian mitochondrial function.^82^ As such, determining the potential mammalian targets of the CYP125/142 inhibitors is also important for future research.

Many drug discovery campaigns that are initiated from a target-centric or in vitro approach fail due to a lack of cellular activity, often as a result of inadequate drug permeability or susceptibility to efflux.^83,84^ Our approach attempted to address these challenges from the outset by screening a tailored fragment library that was biased away from azoles, which are common efflux substrates,^42,43^ and by selecting a ligand efficient hit fragment with a distinct structure to existing drugs.^85^ The good anti-tubercular activity of **5m** against both extra- and intracellular *Mtb* suggests that the compound is able to adequately penetrate both mammalian cells and the complex mycobacterial cell wall, however, a direct analysis of intracellular exposure was not performed (Table 2, Fig. 5). Furthermore, we anticipated that like bedaquiline, and other compounds that deplete cellular ATP, **5m** should inherently decrease *Mtb* efflux transporter activity, thus potentially increasing the efficacy of other antimicrobial drugs.^7,12^

In summary, we have reported an efficient fragment-based approach to develop the first cell active dual CYP125/142 inhibitors. The potency of these compounds against priority *Mtb* populations, including intracellular and MDR bacteria, low toxicity towards human macrophages, and distinct chemical scaffold from existing compounds are promising for their further optimization as chemical tools or antibiotics. The anti-tubercular activity of the CYP125/142 inhibitors exemplifies that expanding the scope of biological pathways considered for drug development offers potential for the development of antibiotics with new mechanisms of action. In this respect, (host)-microbial metabolism is ripe with potentially druggable targets that await exploitation.

## Acknowledgements

M.E.K. was supported by a Commonwealth (University of Cambridge) Scholarship awarded in conjunction with the Cambridge Commonwealth Trust and Cambridge Overseas Trust. A.G.C. and K.J.M. were supported by grants from the BBSRC (grant no. BB/I019669/1 and BB/I019227/1) and M.S. was supported by BBSRC (grant no. BB/M011208/1). This work was funded in part by the Division of Intramural Research of the NIAID/NIH and we acknowledge the Diamond Light Source and the staff of the beamlines i02, i04, and i24 (proposal mx8997, mx17773, and mx24447) for assistance that contributed to the results presented here.

## Author Information

### Authors

Anthony G. Coyne – Yusuf Hamied Department of Chemistry, University of Cambridge, Lensfield Road, Cambridge, CB2 1EW, UK;

Sophie H. Gilbert - Department of Chemistry, University of Cambridge, Lensfield Road, Cambridge, CB2 1EW, UK. Present address: ProQR Therapeutics, Leiden, The Netherlands:

Cecilia Amadi – Centre for Synthetic Biology of Fine and Specialty Chemicals (SYNBIOCHEM), Manchester Institute of Biotechnology, University of Manchester, 131 Princess Street, Manchester, M1 7DN, UK. *Present address*: Department of Experimental Pharmacology & Toxicology, University of Port Harcourt, East/West Road, PMB 5323 Choba, Rivers State, Nigeria.

Matthew Snee – Centre for Synthetic Biology of Fine and Specialty Chemicals (SYNBIOCHEM), Manchester Institute of Biotechnology, University of Manchester, 131 Princess Street, Manchester, M1 7DN, UK. *Present address:* Wellcome Trust Centre for Cell-Matrix Research, School of Biological Sciences, University of Manchester, Oxford Road, Manchester M13 9PT, UK

Richard B. Tunnicliffe – Centre for Synthetic Biology of Fine and Specialty Chemicals (SYNBIOCHEM), Manchester Institute of Biotechnology, University of Manchester, 131 Princess Street, Manchester, M1 7DN, UK.

Kriti Arora – Tuberculosis Research Section, Laboratory of Clinical Infectious Diseases, National Institute of Allergy and Infectious Disease, National Institutes of Health, Bethesda, Maryland, USA. Present address: Novartis Biomedical Research, 5959 Horton Street, Emeryville, CA-94608, USA.

Helena I. Boshoff – Tuberculosis Research Section, Laboratory of Clinical Infectious Diseases, National Institute of Allergy and Infectious Disease, National Institutes of Health, Bethesda, Maryland, USA.

Alexander Fanourakis – Yusuf Hamied Department of Chemistry, University of Cambridge, Lensfield Road, Cambridge, CB2 1EW, UK. *Present address:* Department of Chemistry, University of Chicago, 5735 S Ellis Ave, Chicago, IL 60637, USA.

Maria Jose Rebello-Lopez - TB Profiling Biology, Global Health R&D, GSK, Severo Ochoa, 2, 28760 Tres Cantos, Spain. Present contact

Fatima Ortega-Muro - Global Health DMPK, Global Health R&D, GSK, Severo Ochoa, 2. 28760 Tres Cantos, Spain. Present contact

Colin W. Levy – Manchester Protein Structure Facility (MPSF), Manchester Institute of Biotechnology, University of Manchester, Manchester, M1 7DN, UK.

Andrew W. Munro – Centre for Synthetic Biology of Fine and Specialty Chemicals (SYNBIOCHEM), Manchester Institute of Biotechnology, University of Manchester, 131 Princess Street, Manchester, M1 7DN, UK

David Leys - Department of Chemistry, Manchester Institute of Biotechnology, University of Manchester, 131 Princess Street, Manchester, M1 7DN, UK.

Chris Abell – Yusuf Hamied Department of Chemistry, University of Cambridge, Lensfield Road, Cambridge, CB2 1EW, UK

---

**Scheme 1.**
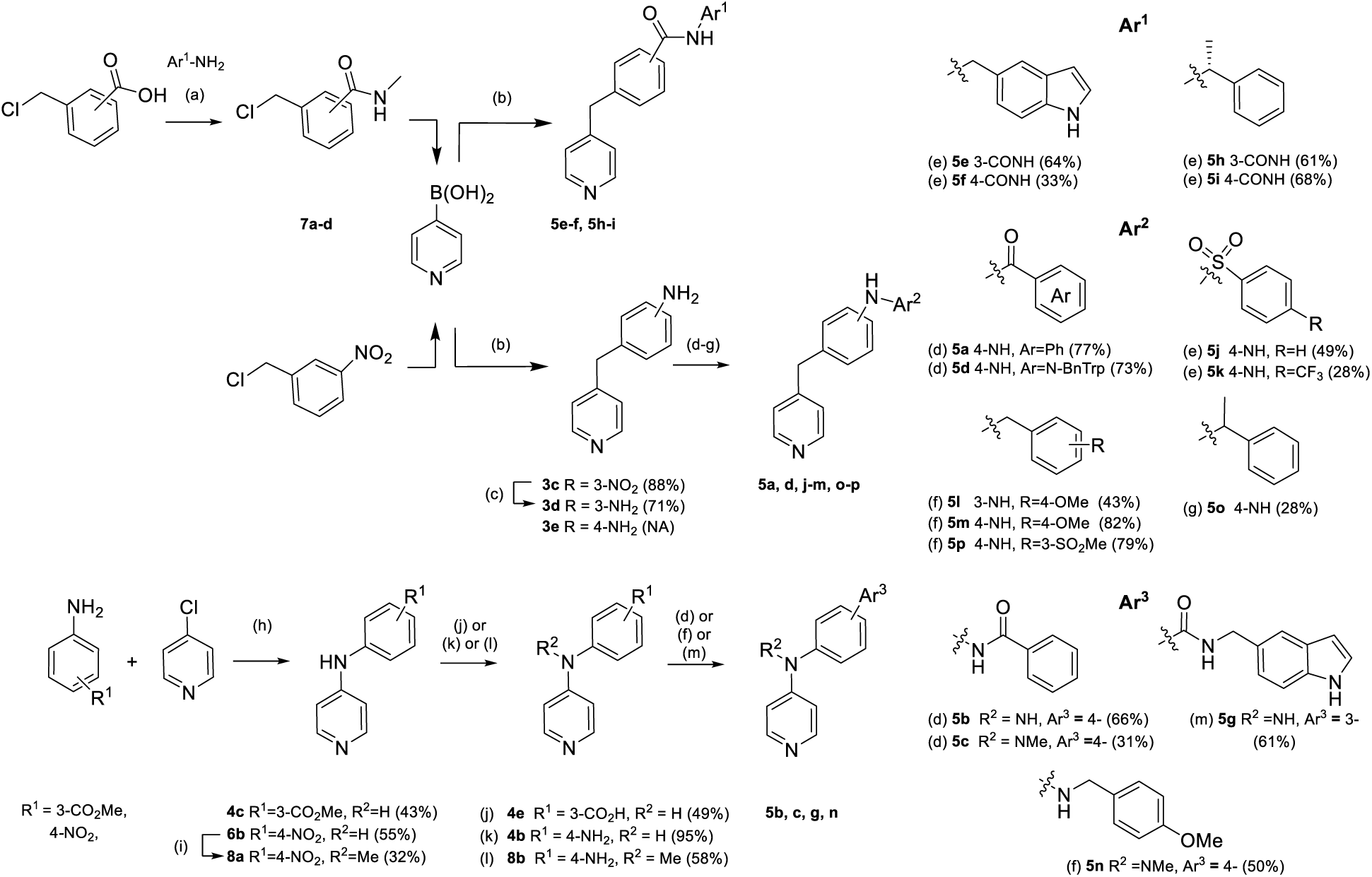
(a) HBTU, Et_3_N, DMC:DMF (7:1), r.t., 24 h; (b) Pd(PPh_3_)_4_, Na_2_CO_3_, DME:H_2_O (2:1), 100 °C, 4 h; (c) Pd/C, N_2_H_4_.xH_2_O, EtOH, 90 °C, 2 h; (d) ArCO_2_H, HATU, DIPEA, DCM, 0 °C−r.t., 24 h (**5a**, **5b**, **5c**); or PyBOP, NMM, DCM, DMF, r.t., 5 h (**5d**); (e) ArSO_2_Cl, pyridine, r.t., 20 h (**5j**); or ArSO_2_Cl, Et_3_N, DCM, r.t., 20 h (**5k**); (f) RCOH, AcOH, NaCNBH_3_, MeOH, r.t., 20 h (**5l**, **5m**, **5p**, **5n**); (g) RCOMe, TiCl_4_, DCM, 0 °C, 3 h; then Na(CN)BH_3_, MeOH, r.t., 24 h (**5o**); (h) HCl (37%), EtOH, 90 °C, 20 h; (i) NaH, DMF, MeI, 0 °C-r.t., 8 h; (j) **4c**, LiOH.H_2_O, MeOH:H_2_O:THF, r.t., 4 h; (k) **6b**, SnCl_2_.2H_2_O, HCl (37%), EtOH, 0-80 °C, 1 h; (l) **8a**, Zn(s), NH_4_Cl, DMF, r.t., 24 h; (m) EDC.HCl, HOAt, DIPEA, DMF:DCM (1:10).

## Table of contents graphic

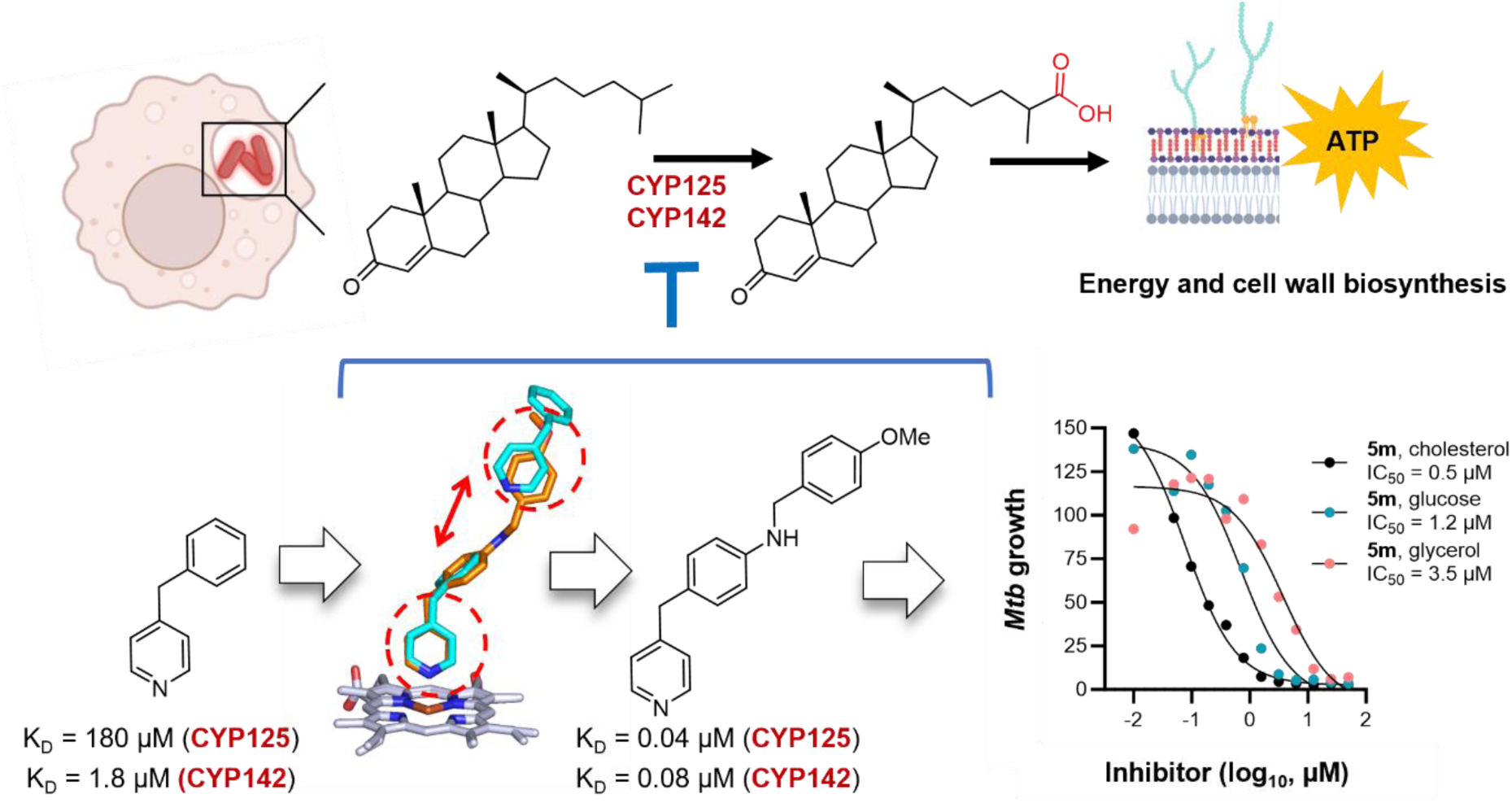

### Synopsis

New drugs are urgently required to treat tuberculosis. Here, we develop small molecules that inhibit *Mtb* cholesterol metabolism, and the growth of intracellular and multi-drug resistant bacteria.

