## Supplementary material for "Fragment-based development of small molecule inhibitors targeting *Mycobacterium tuberculosis* cholesterol metabolism": Materials and Methods

[1]Yusuf Hamied Department of Chemistry, University of Cambridge, Lensfield Road, Cambridge, CB2 1EW, UK. [2]Centre for Synthetic Biology of Fine and Specialty Chemicals (SYNBIOCHEM), Manchester Institute of Biotechnology, University of Manchester, 131 Princess Street, Manchester, M1 7DN, UK. [3]Tuberculosis Research Section, Laboratory of Clinical Infectious Diseases, National Institute of Allergy and Infectious Disease, National Institutes of Health, Bethesda, Maryland, USA. [4]Global Health R&D, GSK, Severo Ochoa, 2, 28760 Tres Cantos, Spain. [5]Manchester Protein Structure Facility (MPSF), Manchester Institute of Biotechnology, University of Manchester, Manchester, M1 7DN, UK. [6]Department of Chemistry, Manchester Institute of Biotechnology, University of Manchester, 131 Princess Street, Manchester, M1 7DN, UK

[#]corresponding authors

.

### Biophysical Methods

#### Protein expression and purification

All experiments, except for the crystal structures obtained of CYP125A1 in complex with inhibitors **5g**, **5j** and **5m**, and CYP142A1 in complex with inhibitor **5j** and **5m**, were performed using *Mtb* CYP125A1 and *Mtb* CYP142A1 proteins that were expressed and purified as previously described.[1,2] In brief, *Cyp125A1* (*Rv3545c*), encoding residues 1-433, and *Cyp142A1* (*Rv3518c*), encoding residues 1-398, were expressed as N-$His_6$-tagged constructs from pET15b vectors in *E. coli* C41(DE3) cells. Bacteria were cultured in 2xYT medium supplemented with ampicillin (100 mg/L) at 37 °C until an OD600 of 0.8. The temperature was then reduced to 23 °C and isopropyl ß-D-1-thiogalactopyranoside (150 µM) was added to induced protein expression, along with 5-aminolevulinic acid (100 µM) to enhance heme incorporation. Bacteria were cultured for a further 18-24 hours and then harvested by centrifugation (9000 g, 4 °C, 20 minutes) and stored at -80 °C until purification. Cell pellets were thawed on ice, resuspended in 50 mM potassium phosphate buffer (pH 8.0), containing 250 mM KCl, 10% v/v glycerol, DNase, lysozyme, protease and phosphatase inhibitors (cOmplete EDTA-free protease inhibitor cocktail tablets, Roche (1 tablet/50 mL), PMSF (1 mM), and benzamidine hydrochloride (1 mM)), and lyzed by sonication. Supernatants were clarified by centrifugation (40,000 g, 4 °C, 30 minutes) and then purified by His-tag affinity chromatography (Ni-NTA (Qiagen) or HisTrap FF (GE Healthcare) eluting with up to 55 m imidazole (CYP142A1) or 200 mM imidazole (CYP125A1). Protein containing fractions were pooled and dialyzed overnight into 50 mM Tris-HCl, pH 7.2, containing 1 mM EDTA and 50 mM KCl, and then purified by anion exchange chromatography (Resource-Q or Q-sepharose, GE Healthcare), eluting with 50-500 mM KCl. Protein containing fractions were pooled and dialyzed into 50 mM Tris-HCl, pH 7.2, containing 1 mM EDTA, concentrated, and purified by gel filtration chromatography (Sephacryl S-200, GE Healthcare). Protein purity and concentration was determined by SDS-PAGE and UV-visible spectroscopy, using the previously established extinction coefficients for CYP125 ($\varepsilon_{449-490}$ = 91 $mM^{-1}$ $cm^{-1}$)[3] and CYP142 ($\varepsilon_{418}$ = 140 $mM^{-1}$ $cm^{-1}$)[1], then aliquots were snap frozen and stored at -80 °C until further use.

Crystal structures of CYP125A1 and CYP142A1 in complex with lead compounds **5g**, **5j**, and **5m** were generated using N-terminally truncated constructs. Cyp125A1 (Rv3545c), encoding residues 18-433, and Cyp142A1 (Rv3518c), encoding residues 2-398, were cloned into a pET21a vector downstream of an T7 leader sequence, and a TEV-cleavable Twin-Strep hexa-histidine dual affinity tag. Proteins were expressed as described above, except that media was supplemented with 250 µM 5-aminolevulinic acid and 200 µM IPTG. Harvested cells were lysed by sonication in 50 mM KPi pH 8.0, 200 mM KCl, 10 % v/v glycerol, supplemented with protease and phosphatase inhibitors, DNase, and lysozyme, and then clarified by centrifugation (42,000 ×g, 4 °C, 1 h). Proteins were purified from supernatants by gravity affinity chromatography using Strep-Tactin XT high capacity resin (IBA Lifesciences), eluting with buffer supplemented with 1x BXT Strep-Tactin elution buffer. The Twin-Strep $His_6$-tag was removed by overnight incubation with Tobacco etch virus protease (TEV) (1:20, TEV:P450), followed by incubation with Nickel-NTA (Qiagen) or Nickel-EXCEL (GE Healthcare) resin for 1 hour. The tag-free protein was collected by gravity filtration, concentrated to 1 mL, and purified by size exclusion chromatography, typically using a Superdex 200 (GE Healthcare) column equilibrated with 20 mM HEPES or Tris-HCl buffer, pH 7.5, containing 200 mM KCl, and 1 mM TCEP. Protein purity and concentration was determined as above, and either directly used in crystallography experiments, or flash frozen.

**Compound screening by UV-visible spectroscopy**

Compounds (1–100 mM) were prepared as stock solutions in $d_6$-DMSO and added as a 2 μL aliquot to solutions of P450 proteins (4–6 μM, 198 μL), or to buffer alone, to achieve a final concentration 1-100 μM compound and 1% v/v $d_6$-DMSO. Samples were either analyzed in quartz cuvettes with a 1 cm path length using a CARY400 UV-vis spectrophotometer (Varian, UK), or in UV-star® 96-well microplates (Greiner Bio-one, UK) using a CLARIOstar microplate reader (BMG Labtech, Germany) in absorbance mode. Spectra were recorded continuously between 800–250 nm at 25 °C. Spectra of the compound in buffer alone were subtracted from protein-containing spectra to account for any inherent UV absorbance of the small molecule. Difference spectra were generated by subtracted the spectrum of an inhibitor-free protein sample from test samples. The magnitude of change in the maximum wavelength of the Soret band of the enzyme's absolute absorbance spectrum ($\lambda_{max}$) relative to a DMSO control (CYP125 $\lambda_{max}$ = 392.5 nm, CYP142 $\lambda_{max}$ = 418 nm), and the change in absorbance between the maximum and minimum wavelengths of the enzymes difference spectrum (ΔAbs), were used to identify P450 ligands. Spectral perturbations that caused a red-shift in the enzyme Soret band ($\lambda_{max}$) was typically classified as Type-II, "inhibitor-like", while those that caused a blue shift were classified as Type I, "substrate-like" interactions.[1–3] As CYP125 is predominantly high spin (HS) at resting state,[4] $\Delta\lambda_{max}$ was calculated for both the HS and low spin (LS) enzyme populations represented in the specta, and the LS/HS ratio was used to further evaluate the degree of LS stabilization, or "inhibition. Perturbations of $\Delta\lambda_{max} < \pm 1$ nm using the CARY400 spectrophotometer, or $< \pm 1.5$ nm using the CLARIOstar microplate reader were considered within experimental error. All UV-vis spectra were generated using Origin software (OriginLab, Northampton, MA) or MARS Data Analysis Software (BMG Labtech). Data were processed using Microsoft Excel (Microsoft Office, 2010).

**Optical titrations to determine dissociation constants**

Optical titrations were performed using a Varian Cary 400 UV-vis spectrophotometer (Varian, CA, USA) according to a previously described procedure.[5] Assays were performed in reduced volume (200 μL) quartz cuvettes with a path length of 1 cm (Starna, Essex, UK). Ligands were prepared as $d_6$-DMSO stock solutions (0.25 mM−500 mM) and proteins (4−6 μM) were prepared in the appropriate buffer. Aliquots (0.2 μL) of ligand stock solutions were added directly to cuvettes containing either protein solutions, or buffer alone. The final $d_6$-DMSO concentration did not exceed 1% v/v of the assay solution. Spectra were recorded between 800−250 nm at 25 °C after the addition of each aliquot of ligand. Buffer control spectra were subtracted from protein spectra to account for any inherent absorbance of added ligands/solvent. Difference spectra were generated by subtracting the initial ligand-free protein spectrum from each successive titration spectrum. The maximum change in absorbance for each difference spectrum was then plotted against ligand concentration and fitted using a one-site binding model hyperbolic/Michaelis-Menten equation (Eq. 1), the Hill function for cooperative binding (Eq. 2) or a modified version of the Morrison equation (Eq. 3) for tight binding inhibitors[6].

Equation 1. $A_{obs} = (A_{max} \times L)/(K_D + L)$

Equation 2. $A_{obs} = (A_{max} \times (L)^n)/((K_D)^n + (L)^n)$

Equation 3**.** $A_{obs} = (A_{max}/2Et) \times ((L + Et + K_D) - (((L + Et + K_D)^2) - (4 \times L \times Et))^{0.5})$

In Equations 1−3, $A_{obs}$ is the observed change in absorbance, $A_{max}$ is the maximum absorbance change at saturation, Et is the enzyme concentration, L is the concentration of ligand, n is the extent of cooperativity and $K_D$ is the dissociation constant for the P450-ligand complex. Data were processed using Microsoft Excel (Microsoft Office, 2013). Data fitting and analysis were performed using Origin software (OriginLab, Northampton, MA) or GraphPad Prism 5.01 (GraphPad Software, San Diego, USA).

**X-Ray crystallography**

The CYP142A1-**1a** structure was obtained using the N-$His_6$-tagged construct. All other structures were obtained using the truncated, tag-free constructs of CYP125A1 and CYP142A1. Crystallization was performed using the sitting-drop vapor diffusion method, at 15-20 mg/mL protein, using a Mosquito nanolitre pipetting robot (TTP labtech). Crystals of CYP125 were obtained in 0.1 M MES buffer, pH 6.5, containing 1.5 - 2.1 M ammonium sulfate. Crystals of CYP142A1 used compound **5j** and **5m** were obtained in 0.1 M sodium acetate, pH 4.5, containing 0.1 M potassium bromide, 8% PEG 20,000, and 8% PEG 550 MME. Compounds were prepared as saturated DMSO stocks and diluted with crystallization mother liquor to 2.5% of total volume. These soaking solutions were pipetted onto drops containing crystals at 4°C for at least 24 hours, then crystals were harvested, cryoprotected using paratone oil, and frozen in liquid nitrogen for data collection. The structure of CYP142A1 in complex with fragment **1a** was obtained using CYP142A1 (15 mg/mL) crystallized in 0.1 M sodium acetate, pH 4.8, containing 0.1 M potassium thiocyanate, 8% PEG 200 and 10% PEG 550 MME. Crystals were back-soaked with 24% PEG 550 MME to remove PEG200 and then soaked with 4 mM fragment **1a**. Diffraction datasets were collected at Diamond light source in Oxfordshire at various beamlines. Data was integrated using the DIALS pipeline,[4] with scaling and merging performed using aimless.[5] Crystallographic models were solved using molecular replacement using the published ligand-free enzyme structures (3IW0 for CYP125 and 2XKR for CYP142). Model building was performed using COOT[6] with ligand restraints generated using ACEdrg.[7] Refinement was performed using PHENIX.refine.[8] Data tables and statistics are provided in **S.I. Table 3**, and ligand density maps are in **S.I. Figure 1**. All structures have been deposited in the Protein Data Bank (http://www.rcsb.org/pdb/) under the accession codes: CYP142-**1a** (8S53), CYP142-**5j** (7QQ7), CYP142-**5m** (7P5T), CYP125-**5j** (7ZGL), CYP125-**5m** (7ZIC), CYP125-**5g** (8S4M). Images of crystal structures were generated using an academic version of the PyMOL Molecular Graphics System, Version 1.3, 2010, Schrödinger, LLC.

#### Biological Activity

##### Safety Statement

All experiments using *M. tuberculosis* strains H37Rv, Edrman, and CDC1551, and luciferase-modified variants, carry some risk of infection and were performed using appropriate safety protocols in BSL3 certified laboratories. The protocols described herein do not pose a high risk for aerosolization.

No other unexpected or unusually high safety hazards were encountered in chemical or biological methods.

##### Substrate turnover assay

Substrate turnover and inhibition assays were set up using either CYP125 (0.5 μM) or CYP142 (1 μM), 10 μM spinach ferredoxin and 1.5 μM spinach ferredoxin reductase in 50 mM KPi buffer, pH 7.5, containing 150 mM KCl, and 0.05% Tween-20 (KPi buffer). Cholest-4-en-3-one (10 mM) was prepared in 45% (v/v) HPCD, and compound stock solutions were diluted in DMSO to 25-100 mM. CYP-ferredoxin mixtures were preincubated with cholest-4-en-3-one (5 μM) and compounds (0-100 μM) for 30 mins at 25 ºC, and then substrate turnover was initiated by the addition of an NADPH regeneration system consisting of 1 mM NADPH, 10 mM glucose-6-phosphate and 2 U glucose-6-phosphate dehydrogenase in KPi buffer. Reactions were allowed to proceed for between 0 - 45 minutes with shaking at 750 rpm at 30 °C, then quenched by the addition of an equal volume of acetonitrile, followed by shaking at 900 rpm for 10 minutes. Samples were then filtered through protein precipitation plates (Phenomenex) under vacuum into mass spectrometry plates. Turnover of cholest-4-en-3-one was monitored by LC-MS using an Agilent 6545XT Advance Bio LC/Q-TOF, equipped with a 2.1 x 100 mm, 1.8 μM Agilent Eclipse Plus C18 column and an elution gradient of 0.1% formic acid in water to 0.1% formic acid in acetonitrile. Samples were quantified with reference to an androstenedione internal standard and a cholest-4-en-3-one calibration curve. Reactions were performed at a range of substrate concentrations, with 5 μM being selected as optimal for calculating $IC_{50}$ values for this compound series. Control reactions were also performed in the absence of NADPH. Data (n=3) were analyzed using Agilent MassHunter Quantification software and resulting $IC_{50}$ curves were fitted in OriginLab graphing software. $IC_{50}$ values were converted to $K_I$ values using the Cheng-Prusoff Equation, (cholestenone $K_m$ CYP125 = 2.1 μM, $K_m$ CYP142 = 0.36 μM).

##### Inhibition of extracellular *Mtb* (H37Rv) growth on defined carbon sources

*M. tuberculosis* (*Mtb*) H37Rv was grown in Middlebrook 7H9 broth medium (Difco) supplemented with 0.3 g/L casitone, 0.81 g/L NaCl, 0.05% (v/v) tyloxapol, and either 97 mg/L cholesterol, 4 g/L glucose, or 0.2% glycerol. For inhibitor assays, a 10-fold serial dilution of the test compounds was made in the desired medium in duplicate rows of a 96-well plate. *Mtb* cells were then added to all the wells at the final concentration of 1 x $10^4$ CFU, and plates were incubated at 37 °C for up to 15 days. Minimum inhibitory concentrations ($MIC_{99}$) were determined using the Microplate Alamar Blue Assay (MABA) on day 15. In brief, resazurin reagent (1:10 dilution of Alamar Blue reagent, Invitrogen) was added to the MIC plates and the cultures were incubated for 24 hours at 37 °C. The concentration of compound required to completely inhibit resorufin fluorescence was determined visually.

Depletion of intracellular ATP concentration by 50% ($IC_{50}$) values were determined as described above except that measurements were made on day 8 and day 15. BacTitre Glo reagent (Promega) was added to microtiter plates (1:10 dilution) and luminescence was recorded after 15 minutes of incubation at room temperature. $IC_{50}$ values were determined to be the concentration of compound required to reduce luminescence by 50% relative to DMSO-treated controls. Isoniazid or *p*-amino salicylic acid were used as positive control compounds for all experiments. Reported MIC and $IC_{50}$ values are the mean of duplicate treatments.

**Inhibition of extracellular MDR-TB growth**

MDR-TB strain K26b00MR 113, which is resistant to isoniazid and rifampicin, was cultured on glucose-casitone media (as described above) and treated with DMSO, compound **5m** (50 – 0.05 µM), or isoniazid. *Mtb* growth was monitored on day 7 and day 14 post-compound treatment by MABA and is reported as the minimum concentration required to inhibit 90% growth ($MIC_{90}$). Assays were performed in duplicate and data are mean values.

**Inhibition of extracellular *Mtb* (Erdman) growth**

*Mtb* (Erdman) was maintained in Middlebrook 7H9 broth medium containing 2% v/v glycerol, 5% w/v BSA, 2 g/L dextrose, and 3 mg/L catalase; supplemented with 2% w/v glucose. Three days prior to the assay, the Erdman strain was pre-adapted to cholesterol by switching the glucose containing 7H9 media to that supplemented with 0.01% w/v cholesterol. Cultures were treated with DMSO or compounds (50 - 0.098 µM), and growth inhibition was assessed 7 days post-compound treatment by the addition of resazurin. Plates were incubated for 48 hours and then fluorescence was recorded. Assays were performed in triplicate and all plates contained a Moxifloxacin (0.005 - 2.5 µM) treated controls, which corresponded to 100% inhibition of *Mtb* growth. Percent growth in each well was calculated relative to maximum signal intensity in uninhibited wells, and $IC_{50}$ values were estimated by non-linear regression (3-parameter), using GraphPad Prism v10.0.1.

**HepG2 Cytotoxicity assay**

Compounds were tested against HepG2 cells for their ability to inhibit ATP production as a measure of cytotoxicity. HepG2 cells were cultured in DMEM supplemented with 10% (v/v) FBS, Hepes, L-glutamine and glucose or galactose. Compounds were serial diluted in microtiter plates (as described above) and HepG2 cells were added at a final concentration of 20,000 cells per well. Inhibition of ATP levels was noted by adding CellTitre Glo (Promega) reagent at a 1:10 dilution and luminescence was noted after 10 minutes incubation at room temperature.

**Mechanisms of action reporter assays**

Assays to determine compound mechanisms of action were performed using bioluminescent transcriptional reporter *Mtb* strains as described previously.[7] In brief, compounds were prepared as a 2-fold serial dilution in 96-well plates (50 – 0.05 µM) and *Mtb/iniB*, *Mtb/recA*, or *Mtb/radA* (27572410) was added to final concentration of 1 x $10^6$ cells per well. The plates were incubated at 37 °C for 1 week and luminescence was recorded on days 1, 2, 4 and 7 post-compound treatment. Signal intensity was normalized as a % of the maximum signal intensity induced by control compounds which inhibit cell wall synthesis (SQ109, top concentration = 100 μM), or induce DNA damage (moxifloxacin, top concentration = 25 μg/mL), respectively. Assays were performed in duplicate, and data are shown as mean values ± SD.

**Intracellular growth assay**

Intracellular growth assays were performed as previously described,[8] with minor modifications. *Compounds* – Compounds were prepared as DMSO stock solutions and 50 µL was dispensed into assay plates as an 11-point 3-fold serial dilution using a HP Dispenser D300e Control, V3.3.1, Device 2.69.0.0 (Tecan). The final DMSO concentration was 0.5% and all compounds were tested in duplicate.

*THP-1 cell culture* – THP-1 cells (ATCC TIB-202) were maintained in RPMI-1640 (Sigma R5886), supplemented with 10% FBS (FBS SOUTH AMERICAN (CE), Gibco #10270,), 1 mM sodium pyruvate (Sigma, #S8636) and 2 mM L-glutamine (Sigma, #G2150) at 37º C, 5% $CO_2$, and 95% humidity. Cells were handled according to GSK policies for management of human biological samples.

*Preparation of Mtb single bacteria suspensions.* – The luminescent strain *Mtb* H37Rv pATB45luc grown at 37 °C in Middlebrook 7H9 medium (Difco) supplemented with 0.2% glycerol, 0.5% bovine albumin fraction, 0.2% dextrose, 0.003% catalase (Becton Dickinson), 0.05% tyloxapol (Merck). Hygromycin B was added to the medium at a final concentration of 50 µg/mL. All experimental work with live *Mtb* H37Rv was carried out following standard operating procedures in compliance with Biosafety Level 3 regulations (BSL3). A single bacteria suspension of *Mtb* H37Rv pATB45luc was prepared prior to infection. 25 mL of bacterial culture grown to $OD_{600}$= 0.6 (log phase in our conditions is between $OD_{600}$ from 0.05 - 1) was centrifuged at 2230*g* for 10 min. After removal of the supernatant, bacteria were dispersed by vigorously shaking with sterile glass beads 4MM (201-0278 VWR) for 2 min. Dispersed bacteria were then re-suspended in 35 mL of RPMI medium and left to decant for 5 min at room temperature. 30 mL of the supernatant were centrifuged at 308g for 5 min. Supernatant was collected and its $OD_{600}$ was measured. OD/mL was converted to CFU/mL ($OD_{600}$ 0.125 ~ $10^8$ CFUs/mL).

*Infection of THP-1 cells with Mtb –* 1 x $10^6$ THP-1 cells were simultaneously differentiated with phorbol myristate acetate (PMA, 40 ng/mL, Sigma, #P1585) and infected with a single cell suspension of *Mtb* H37Rv pATB45luc in a roller bottle at a MOI of 1:10. Cells were incubated for 4 hours at 37° C at 1.5 rpm. After incubation, infected cells were washed five times by centrifugation at 308g for 5 min to remove extracellular bacilli and re-suspended in fresh RPMI medium. In the last wash, a Falcon cell strainer 40 µm (Corning) was used to remove cells clumps. The infected cells were re-suspended in RPMI medium supplemented with 10% FBS, 2 mM L-glutamine and 1 mM sodium pyruvate at a concentration of 2 x $10^5$ cells/mL. 50 µL of this cell suspension (10,000 cells) were dispensed into 384-well plates containing compounds. Plates were incubated at 37° C, 5% $CO_2$ and 90% relative humidity for 5 days. On day 5, luminescence, which is proportional to bacterial load, was determined by using the BrightGlo™ Luciferase Assay System (Promega, # E2650) according to the manufacturer's protocol, except that 20 µL of Bight-Glo™ mix was used instead of 50 µL. Plates were read using an Envision Multilabel Plate Reader (PerkinElmer) using the 384-plate Ultra-Sensitive luminescence mode, with a measurement time of 200 ms per well. All plates were assayed in duplicate.

*Data Analysis* – All assay plates contained a DMSO-treated column which correspond to 100% bacterial growth, and a Rifampicin (5 µM) treated column, which corresponds to 100% inhibition of *Mtb* growth, which were used to assess assay quality (Z' ≥ 0.4) and to normalize data on a per-plate basis. Growth (%) for each well was calculated relative to the maximum signal intensity in the uninhibited samples. $IC_{50}$ values for each compound was estimated by non-linear regression (3-parameter) using GraphPad Prism 10.0.01

**THP-1 cytotoxicity**

THP-1 cells (ATCC TIB-202) were maintained in RPMI-1640 (Sigma R5886), supplemented with 10% FBS (FBS SOUTH AMERICAN (CE), Gibco #10270,), 1 mM sodium pyruvate (Sigma, #S8636) and 2 mM L-glutamine (Sigma, #G2150) at 37º C, 5% $CO_2$, and 95% humidity. For cytotoxicity assays, THP-1 monocytes were seeded at 5 x $10^5$ cells/mL and treated with 40 ng/mL phorbol myristate acetate (PMA) (Sigma, # P1585) to differentiate for 4 hours. The cells were then harvested, washed with complete medium, adjusted to 2 x $10^5$ cells/mL, and 50 µL (10,000 cells) was transferred to each well of clear-bottomed, sterile 384-well plates (Greiner, #781095), which already contained 250 µL/well of DMSO/compounds diluted in media. Diluted compounds were prepared as a 12-point serial dilution (1:3) from a top concentration 50 µM. Compounds were assayed in duplicate in each assay plate. Each assay plate contained a column of DMSO negative controls which correspond to 100% growth, and a column of doxorubicin (Sigma, #D1515) positive controls which correspond to 100% inhibition of growth, which were used to monitor assay quality (Z' > 4). Hygromycin (Sigma) was then added at the final concentration of 0.1 mg/ml, and the plates were incubated for 5 days at 37 ºC, 5% CO2 and 95% humidity. Luminescence was measured on day 5 using the ATPLite 1-step kit (Perkin Elmer, #6016739 ). Briefly, 25 µL of reconstituted substrate solution was added to each well, the plate was shaken for 1 minute in the dark, and then luminescence was measured using EnVision Multilabel Reader (PerkinElmer), using measurement time 0.1 s. Percent growth inhibition was calculated as: %Inhibition = 100 × [(data – DMSO)/(doxorubicin – DMSO)]. The concentration of the compound necessary to inhibit 50% of THP-1 cell growth ($LD_{50}$) and was calculated by fitting %inhibition data by nonlinear regression (GraphPad Prism).

#### Chemical synthesis and characterization

**General.** All reagents were commercially sourced unless otherwise specified. All reactions were conducted under the positive pressure of a dry nitrogen atmosphere. Anhydrous solvents were either freshly distilled (DCM and MeOH over $CaH_2$, THF over $CaH_2$ and $LiAlH_4$) or purchased from commercial sources. Reactions were monitored by liquid chromatography mass spectrometry (LCMS) or thin layer chromatography (TLC), using Merck glass-backed silica (Kieselgel 60 F254 0.25 mm) plates. TLC plates were visualized under UV (254/365 nm) and retention factors (Rf) are provided for the noted solvent system. Flash column chromatography was performed using an IsoleraTM Spektra One/Four purification system and either a GraceResolv™ LOK flash cartridge containing silica gel (40 μm) (Grace Discovery Sciences, USA) or Biotage SNAP column containing KP-silica gel (50 μm). Solvents are reported as volume/volume (v/v) eluent mixture. Proton ($^1$H) and carbon ($^{13}$C) nuclear magnetic resonance (NMR) spectra were recorded at 300 K using either a Bruker 400 MHz AVANCE III HD Smart Probe, 400 MHz QNP cryoprobe or 500 MHz DCH cryoprobe spectrometer. Chemical shifts are given in parts per million (ppm) (δ), relative to residual protonated solvent peak of the deuterated solvent indicated, and the relative integral, multiplicity, and coupling constants (*J* Hz) of the peaks is noted. Assignment of $^1$H-NMR and $^{13}$C-NMR spectra was assisted by DEPT, and 2D NMR experiments (COSY, edited $^1$H−$^{13}$C-HSQC and 1H−13C HMBC) where necessary. Infrared (IR) absorption spectra were recorded on a Spectrum One™ FT-IR (Perkin Elmer) spectrometer by attenuated total reflectance (ATR). Data are reported as vibrational frequency ($v_{max}$, cm$^{-1}$) and peak intensity - strong (s), medium (m), weak (w) or broad (br). LCMS was carried out using an AQUITY UPLC H-class system (Waters, Manchester UK). Samples were either run under acidic conditions on an ACQUITY UPLC HSS C-18 column, eluting with a gradient of 95-5% v/v water (containing 0.1% formic acid) in MeCN, or under basic conditions on an ACQUITY UPLC BEH130 C18 column, eluting with a gradient of 95-5% v/v water (containing 10 mM $NH_4OAc$) in MeCN over a period of 3.5 minutes. Small molecule high resolution mass spectrometry (HRMS) was carried out using a Micromass Quadrapole-Time-of-flight (Q-Tof) mass spectrometer, Waters Xevo G2-XS QTof mass spectrometer or a ThermoFinnigan Orbitrap Classic LCMS spectrometer attached to a Dionex Ultimate 3000 HPLC. The mass to charge ratio (*m/z*) of the molecular ion and difference from calculated mass (*δ* ppm) have been quoted. All final compounds used in protein binding or cell-based assays had a purity of > 95% by LCMS analysis.

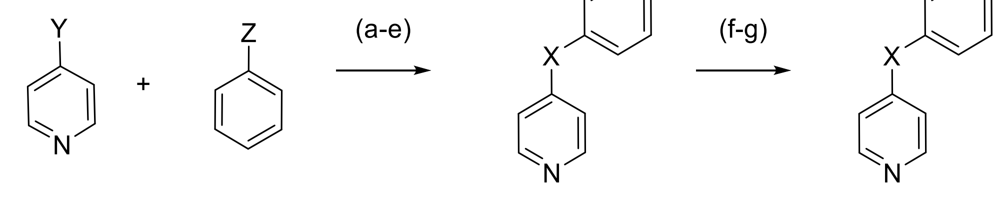


Y = Cl, I, Br, $CH_3$

Z = $NH_2$, NHPh, OH, $SO_3Na$, COH

**2a** X = NH (a, 53%)
**2c** X = NPh (b, 6%)
**2d** X = O (c, 4%)
**2e** X = $SO_2$ (d, 2%)
**2g** X = *E*-CH=CH (e, 39%)

**2b** X = NMe (f, 75%)
**2h** X = $CH_2CH_2$ (g, 94%)

**S.I. Scheme 1.** Synthesis of **1a-i** analogue to explore SAR of benzylic position. *Reagents and conditions:* (a) Y=Cl.HCl, Z=$NH_2$, HCl (37%), EtOH, 90 °C, 20 h; (b) Y=Br.HCl, Z=NHPh, K$^t$OBu, $Pd(OAc)_2$, *rac*-BINAP, toluene, 70 °C, 16 h; (c) Y=Cl.HCl, Z=OH, Cu(s) powder, $Cs_2CO_3$, DMF, 100 °C, 18 h; (d) Y=I, Z=$SO_3$Na, *L*-proline sodium salt, CuI, DMSO, 80 °C, 44 h; (e) Y=$CH_3$, Z=COH, $Ac_2O$, 140 °C, 24 h; (f) **2a**, MeI, K$^t$OBu, DMF, r.t. 19 h; (g) **2g**, $H_2$(g), Pd/C, EtOH, r.t., 20 h.

**N-Phenylpyridin-4-amine (2a)**

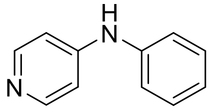

Aniline (273 μL, 3.00 mmol) was added to a solution of 4-chloropyridine hydrochloride (450 mg, 3.00 mmol) in EtOH (15 mL), followed by a catalytic amount of conc. HCl (37%, 4 drops), and the reaction was heated at 90 °C for 20 hours. The reaction was then concentrated to approximately 7 mL and quenched with saturated NaHCO3. The aqueous phase was extracted with EtOAc (4 x 20 mL) and the organic fractions were combined, dried over anhydrous Na2SO4 and the solvent was removed under reduced pressure. The crude product was purified by flash chromatography (0-10% v/v MeOH in DCM) to yield compound 2a as a white solid (286 mg, 1.58 mmol, 53%). Rf 0.07 (10% v/v MeOH in DCM); 1H-NMR (400 MHz, CDCl3) δ 8.26 (d, J = 6.5 Hz, 2H), 7.36 (app. t, J = 7.9 Hz, 2H), 7.20 (d, J = 7.6 Hz, 2H), 7.13 (t, J = 7.4 Hz, 1H), 6.83 (d, J = 6.6 Hz, 1H), 6.47 (br s, 1H) ppm; 13C-NMR (100 MHz, CDCl3) δ 151.0, 150.0, 139.6, 129.7, 124.4, 121.8, 109.6 ppm; IR (solid) vmax 3054-2700 (w, br, N-H), 2896, 2838 (w, C-H), 1610 (m, C=C), 1585 (s, pyridine CC/CN), 1524 (m, C=C, N-H), 1491 (s, N-H), 1448 (m, C=C), 1349 (m, C-N), 1334 (s, C-N), 1236 (w, C-H), 1217 (m, C-H), 994 (s, C-H), 893 (w, C-H), 806 (m, C-H), 749, 694 (s, C-H), 638 (w, C=H) cm-1; LCMS (+ESI) m/z 171.1 [M+H]+, retention time 1.73 min, (100%); HRMS (+ESI) m/z (Calcd. C11H11N2 [M+H]+, 171.0917), Obs. 171.0913 (δ 2.1 ppm).

***N*-Methyl-*N*-phenylpyridin-4-amine (2b)**

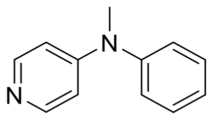

Potassium *tert*-butoxide (53 mg, 0.47 mmol) was added as a single portion to a solution of *N*-phenylpyridin-4-amine **2a** (20 mg, 0.12 mmol) in anhydrous DMF (3 mL) at room temperature. Methyl iodide (30 µL, 0.47 mmol) was added and the reaction was stirred at room temperature for 19 hours, before being diluted with water (5 mL). The product was extracted into $Et_2O$ (3 x 10 mL), the combined organic fractions were dried over anhydrous $Na_2SO_4$ and then the solvent was removed under reduced pressure. The crude product was purified by flash chromatography (0-10% v/v MeOH in DCM) to yield compound **2b** as a yellow amorphous solid (17 mg, 0.09

mmol, 75%). $^1$H-NMR (500 MHz, $CDCl_3$) $\delta$ 8.17 (d, $J$ = 6.3 Hz, 2H), 7.44-7.40 (m, 2H), 7.27 (dd, $J$ = 7.4, 1.2 Hz, 1H), 7.19 (d, $J$ = 7.4 Hz, 2H), 6.54 (dd, $J$ = 5.1, 1.5 Hz, 2H), 3.31 (s, 3H) ppm; $^{13}$C-NMR (125 MHz, $CDCl_3$) $\delta$ 154.3, 149.1, 146.1, 130.3, 126.9, 126.8, 108.4, 39.7 ppm; IR (solid) $\nu_{max}$ 3035, 2921, 1642, 1604, 1584, 1493, 1361, 1224, 985, 808 cm$^{-1}$; LCMS (+ESI) *m/z* (Calcd. $C_{12}H_{13}N_2$ $[M+H]^+$, 185.1), *Obs.* 185.2.

***N,N*-Diphenylpyridin-4-amine (2c)**

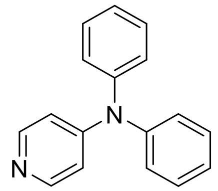

Diphenylamine (217 mg, 1.3 mmol), $Pd(OAc)_2$ (11 mg, 0.05 mmol), *rac*-BINAP (31 mg, 0.05 mmol) and potassium *tert*-butoxide (321 mg, 2.9 mmol) were added to a stirred suspension of 4-bromopyridine hydrochloride (207 mg, 1.1 mmol) in toluene (9 mL). The reaction was heated at 70 °C for 16 hours, then cooled to room temperature and the product was extracted into $Et_2O$ (20 mL). The combined organic fractions were washed with brine (3 x 30 mL), dried over anhydrous $Na_2SO_4$ and the solvent was removed under reduced pressure. The crude product was purified by flash chromatography (0-15% v/v EtOAc in DCM) to yield compound **2c** as a brown solid (15 mg, 0.06 mmol, 6%). $^1$H-NMR (400 MHz, $CDCl_3$) $\delta$ 8.21 (d, $J$ = 6.2 Hz, 2H), 7.34 (app. t, $J$ = 7.9 Hz, 4H), 7.26-7.15 (m, 6H), 6.72 (dd, $J$ = 5.0, 1.5 Hz, 2H) ppm; $^{13}$C-NMR (100 MHz, $CDCl_3$) $\delta$ 153.9, 150.4, 145.4, 130.0, 126.9, 125.8, 113.0 ppm; IR (solid) $\nu_{max}$ 2988, 2902, 1575, 1483, 1450, 1338 1075, 812, 694 cm$^{-1}$; LCMS (+ESI) *m/z* (Calcd. $C_{17}H_{15}N_2$ $[M+H]^+$, 247.1), *Obs.* 247.0.

**4-Phenoxypyridine (2d)**

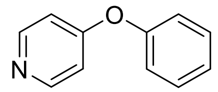

Phenol (141 mg, 1.5 mmol), copper powder (6.4 mg, 0.1 mmol) and $Cs_2CO_3$ (1.0 g, 3 mmol) were added to a stirred suspension of 4-chloropyridine hydrochloride (150 mg, 1 mmol) in anhydrous DMF (2.2 mL). The reaction was placed under an inert atmosphere are heated to 100 °C for 18 hours, then allowed to cool to room temperature and diluted with DCM (20 mL). The solution was washed with 1 M NaOH (40 mL) and water (35 mL), then dried over anhydrous $Na_2SO_4$ and the solvent was removed under reduced pressure. The crude material was purified by flash chromatography (0-5% v/v MeOH in DCM) to yield compound **2d** as a brown solid (6.2 mg, 0.04 mmol, 4%). $^1$H-NMR (400 MHz, $CDCl_3$) $\delta$ 8.46 (m, 2H), 7.43 (app. t, $J$ = 7.9 Hz, 2H), 7.26 (t, $J$ = 7.4 Hz, 1H), 7.10 (d, $J$ = 8.4 Hz, 2H), 6.84 (d, $J$ = 8.4 Hz, 2H) ppm; $^{13}$C-NMR (100 MHz, $CDCl_3$) $\delta$ 165.1, 154.2, 151.5, 130.4, 125.7, 121.0, 112.4 ppm; IR (solid) $\nu_{max}$ 3054, 2920, 2850, 1598, 1572, 1497, 1485, 1264, 1210, 990, 821 cm$^{-1}$; LCMS (+ESI) *m/z* (Calcd. $C_{11}H_{10}NO$ $[M+H]^+$, 172.1) *Obs.*172.2 $[M+H]^+$.

**4-(Phenylsulfonyl)pyridine (2e)**

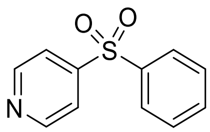

4-Iodopyridine (205 mg, 1 mmol), benzenesulfinic acid sodium salt (197 mg, 1.2 mmol), L-proline sodium salt (27 mg, 0.2 mmol) and CuI (19 mg, 0.1 mmol) were combined in DMSO (3 mL) and the reaction was stirred at 80 °C for 24 hours. An addition equivalent of L-proline sodium salt (27 mg, 0.2 mmol) and CuI (19 mg, 0.1 mmol) were then added, and the reaction was heated for a further 19 hours, before being allowed to cool to room temperature. The reaction was then diluted with water (20 mL), washed with brine (20 mL), the organic fraction was dried over anhydrous $MgSO_4$ and then the solvent was removed under reduced pressure. The crude product was purified by flash chromatography (0-50% v/v EtOAc in DCM) to yield compound **2e** as an off-white solid (5.1 mg, 0.02 mmol, 2%). $^1$H-NMR (400 MHz, $CDCl_3$) $\delta$ 8.81 (d, $J$ = 6.0 Hz, 2H), 7.95 (d, $J$ = 7.5 Hz, 2H), 7.75 (dd, $J$ = 4.5, 1.6 Hz, 2H), 7.63 (dd, $J$ = 7.5, 1.6 Hz, 1H), 7.54 (app. t, $J$ = 7.5 Hz, 2H) ppm; $^{13}$C-NMR (100 MHz, $CDCl_3$) $\delta$ 151.4, 150.0, 140.0, 134.4, 129.9, 128.4, 120.8 ppm; IR (solid) $\nu_{max}$ 3082, 2922, 2850, 1572, 1475, 1450, 1404, 1324, 1158, 1112, 739 cm$^{-1}$; LCMS (+ESI) *m/z* (Calcd. $C_{11}H_{10}NO_2S$ [M+H]$^+$, 220.0) *Obs.* 220.2 [M+H]$^+$.

**(*E*)-4-Styrylpyridine (2g)**

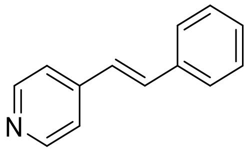

Benzaldehyde (1.12 mL, 11 mmol) was added to a stirred solution of 4-picoline (0.98 mL, 10 mmol) in acetic anhydride (10 mL). The reaction was heated under reflux (140 °C) for 2 hours and the solvent was removed under reduced pressure. Iced water (30 mL) was added to the resulting residue and the mixture was brought to pH 1 using 10% HCl solution. The mixture was washed with DCM (5 -x 25 mL) and then the aqueous fraction was neutralized using 2.5 M NaOH. The product was extracted into DCM (5 x 20 mL) and then the combined organic fractions were passed through a short plug of silica, eluting with DCM (100 mL). The solvent was then removed under reduced pressure to yield compound **2g** as a yellow solid (710 mg, 3.9 mmol, 39%). $^1$H-NMR (400 MHz, $CDCl_3$) $\delta$ 8.58 (d, $J$ = 5.6 Hz, 2H), 7.52 (d, $J$ = 7.5 Hz, 2H), 7.41-7.23 (m, 6H), 7.00 (d, $J$ = 16.4 Hz, 1H) ppm; $^{13}$C-NMR (100 MHz, $CDCl_3$) $\delta$ 150.1, 144.7, 136.1, 133.3, 128.9, 128.8, 127.0, 126.0, 120.9 ppm; IR (solid) $\nu_{max}$ 3025, 1719, 1635, 1589, 1549, 1495, 1455, 1414, 971, 808 cm$^{-1}$; LCMS (+ESI) *m/z* (Calcd. $C_{13}H_{12}N$ [M+H]$^+$, 182.1) *Obs.*183.1 [M+H]$^+$

**4-Phenethylpyridine (2h)**

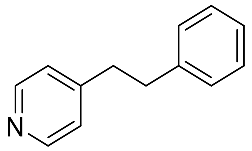

Pd/C (10% wt., 5 mg) was added to a solution of (E)-4-styrylpyridine (X) (50 mg, 0.28 mmol) in EtOH (5 mL) and the reaction as stirred under an atmosphere of $H_2$(g) for 20 hours. When no starting material remained, the reaction was filtered through a short plug of celite and the solvent was removed under reduced pressure to yield compound **2h** as an off-white solid (48 mg, 0.26 mmol, 94%). $^1$H-NMR (400 MHz, $CDCl_3$) $\delta$ 8.46 (d, $J$ = 6.0 Hz, 2H), 7.29-7.11 (m, 5H), 7.06 (d, $J$ = 6.0 Hz, 2H), 2.91 (m, 4H) ppm; $^{13}$C-NMR (100 MHz, $CDCl_3$) $\delta$ 150.7, 149.9, 140.9, 128.7, 128.6, 126.5, 124.2, 37.3, 36.8 ppm; IR (solid) $\nu_{max}$ 3029, 2923, 2858, 1716, 1595, 1558, 1493, 1452, 1410, 1068, 989, 807 cm$^{-1}$; LCMS (+ESI) *m/z* (Calcd. $C_{13}H_{14}N$ [M+H]$^+$, 184.1) *Obs.*184.1 [M+H]$^+$.

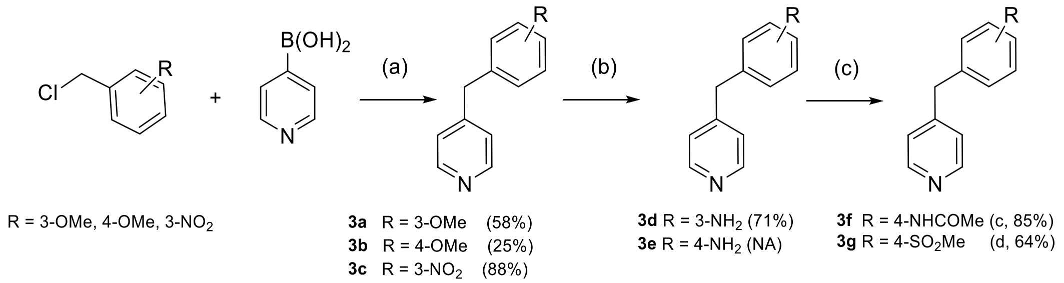


**S.I. Scheme 2.** Synthesis of **1a-i** analogues to explore SAR of "linker". *Reagents and conditions:* (a) $Pd(PPh_3)_4$, $Na_2CO_3$, DME:$H_2O$ (2:1), 100 °C, 4-16 h, or 1 h microwave; (b) **3c**, Pd/C, $N_2H_4$.x$H_2O$, EtOH, 90 °C, 2 h; (c) **3e**, $MeClO_2S$ or $(CH_3CO)_2O$, $Et_3N$, DCM, r.t., o.n. N/A – obtained from commercial sources.

**4-(3-Methoxybenzyl)pyridine (3a)**

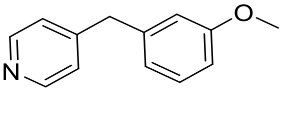

4-Pyridinylboronic acid (73 mg, 0.6 mmol), 3-methoxybenzyl chloride (68 uL, 0.50 mmol), and $Na_2CO_3$ (111 mg, 1.05 mmol) were combined in a microwave vial and flushed with argon. Dry THF (2 mL) and water (1 mL) were added, followed by $Pd(PPh_3)_4$ (11 mg, 0.01 mmol) and the reaction was further flushed with argon. The vial was sealed and heated at 100 °C for 1 hour in a in a microwave reactor. The reaction was cooled to room temperature and diluted with water (5 mL) and DCM (5 mL). The phases were separated and the aqueous phase was extracted with DCM (3 x 5 mL). The organic fractions were combined, dried over anhydrous $Na_2SO_4$ and the solvent was removed under reduced pressure. The crude material was purified by flash chromatography (0-5% v/v MeOH in DCM) and the product containing fractions were combined to yield compound **3a** as a yellow oil (57.9 mg, 0.29 mmol, 58%). $^1$H-NMR (400 MHz, $CDCl_3$) δ 8.48 (dd, *J* = 5.9, 1.6 Hz, 2H), 7.23 (dd, *J* = 8.2, 7.5 Hz, 1H), 7.10 (m, 2H), 6.77 (m, 2H), 6.70 (dd, *J* = 2.1 Hz, 1H), 3.93 (s, 2H), 3.77 (s, 3H) ppm; $^{13}$C-NMR (100 MHz, $CDCl_3$) δ 159.9, 150.0, 149.9, 140.5, 129.8, 124.3, 121.5, 115.0, 111.9, 55.3, 41.3 ppm; LCMS (+ESI) *m/z* 200.1 $[M+H]^+$, 3.47 min, (95%); HRMS (+ESI) *m/z* (Calcd. $C_{13}H_{14}NO$ $[M+H]^+$, 200.1070), Obs. 200.1068 (*δ* 1.2 ppm).

**4-(4-Methoxybenzyl)pyridine (3b)**

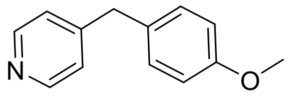

4-Pyridinylboronic acid (73 mg, 0.6 mmol), 4-methoxybenzyl chloride (68 uL, 0.50 mmol), and $Na_2CO_3$ (111 mg, 1.05 mmol) were combined in a microwave vial and flushed with argon. Dry THF (2 mL) and water (1 mL) were added, followed by $Pd(PPh_3)_4$ (11 mg, 0.01 mmol) and the reaction was further flushed with argon. The vial was sealed and heated at 100 °C for overnight (conventional). The reaction was cooled to room temperature and diluted with water (5 mL) and DCM (5 mL). The phases were separated, and the aqueous phase was extracted with DCM (3 x 5 mL). The organic fractions were combined, dried over anhydrous $Na_2SO_4$ and the solvent was removed under reduced pressure. The crude material was purified by flash chromatography (0-5% v/v MeOH in DCM) and the product containing fractions were combined to yield compound **3b** as a colourless oil (25.2 mg, 0.13 mmol, 25%). $^1$H_NMR (400 MHz, CDCl3) δ 8.48 (m, 2H), 7.08 (m, 2H), 6.85 (m, 2H), 3.90 (s, 2H), 3.79 (s, 3H) ppm; $^{13}$C-NMR (100 MHz, CDCl3) δ 158.5, 150.6, 149.9, 131.0, 130.1, 124.2, 114.2, 77.5, 77.2, 76.8, 55.4, 40.4 ppm; LCMS (+ESI) *m/z* 200.1 $[M+H]^+$, 3.43 min, (100%); HRMS (+ESI) *m/z* (Calcd. $C_{13}H_{14}NO$ $[M+H]^+$, 200.1070), Obs. 200.1068 (*δ* 0.8 ppm).

**4-(3-Nitrobenzyl)pyridine (3c)**

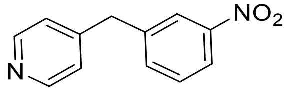

4-Pyridinyl boronic acid (74 mg, 0.60 mmol), 3-nitrobenzyl chloride (86 mg, 0.50 mmol), and $Na_2CO_3$ (111 mg, 0.126 mmol) were combined in a microwave vessel and flushed with $N_2$(g) for 5 minutes. $Pd(PPh_3)_4$ (12 mg, 0.01 mmol) was added, and the reaction vessel was flushed with $N_2$(g) for a further 2-3 minutes before the addition of a mixture of DME and water (2:1 v/v, 3 mL). The reaction was heated at 100 °C for 4 h, then allowed to cool to room temperature and diluted with water (6 mL) and DCM (6 mL). The phases were separated, and the aqueous fraction was extracted with DCM (3 x 3 mL). The organic fractions were combined, dried over anhydrous $Na_2SO_4$ and the solvent was removed under reduced pressure. The crude material was purified twice by flash chromatography (0-5% v/v MeOH in DCM) to yield compound **3c** as an orange crystalline solid (93.8

mg, 0.44 mmol, 88%), which retained a small of $PPh_3$ impurity. $R_f$ 0.47 (10% v/v MeOH in DCM); $^1$H-NMR (500 MHz, $CDCl_3$) $\delta$ 8.55 (d, $J$ = 6.1 Hz, 2H), 8.12 (m, 1H), 8.07 (m, 1H), 7.51 (d, $J$ = 1.1 Hz, 1H), 7.50 (dd, $J$ = 2.1, 1.1 Hz, 1H), 7.11 (d, $J$ = 6.1 Hz, 1H), 4.08 (s, 2H) ppm; $^{13}$C-NMR (125 MHz, $CDCl_3$) $\delta$ 150.2, 148.5, 148.1, 140.9, 135.1, 129.7, 124.0, 123.9, 122.0, 40.8 ppm; IR (solid) $v_{max}$ 3099, 3066, 3024, 1669, 1595, 1561, 1510, 1477, 1439, 1416, 1358, 1346, 1315, 1217, 1098, 1076, 994, 912, 900, 843, 806, 819, 792, 731, 685 (s, C-H), 673, 614 $cm^{-1}$; LCMS (+ESI) *m/z* 215.2 $[M+H]^+$, 1.26 min, (100%); HRMS (+ESI) *m/z* (Calcd. $C_{14}H_{22}N_3O$ $[M+H]^+$, 248.1757), *Obs.* 248.1748 ($\delta$ 3.8 ppm).

**3-(Pyridin-4-ylmethyl)aniline (3d)**

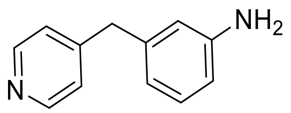

Pd/C (17 mg, 0.16 mmol) was added as a single portion to a warm (55 °C) suspension of the **3c** (172 mg, 0.80 mmol) and hydrazine hydrate (125 µL, 4.02 mmol) in absolute EtOH (12 mL). The reaction was heated at reflux for 2 hours, at which point not starting material remained by LCMS analysis and the solution was colourless. The reaction was cooled to room temperature, filtered through filter paper, then through a short (1-2 cm) plug of silica, using a solution of 10% v/v MeOH in DCM (10-20 mL) to elute the product. The solvent was removed under reduced pressure to yield compound **3d** as a white crystalline solid (105.1 mg, 0.57 mmol, 71%). $R_f$ 0.17 (5% v/v MeOH in DCM); $^1$H-NMR (500 MHz, $CDCl_3$) $\delta$ 8.48 (d, $J$ = 5.9 Hz, 2H), 7.13 (d, $J$ = 5.4 Hz, 2H), 7.10 (app. t, $J$ = 7.8 Hz, 1H), 6.57 (d, $J$ = 7.7 Hz, 2H), 6.47 (app. t, $J$ = 2.0 Hz, 1H), 3.87 (s, 2H) ppm; $^{13}$C-NMR (125 MHz, $CDCl_3$) $\delta$ 150.8, 149.5, 149.4, 146.9, 140.0, 129.8, 124.5, 119.4, 115.7, 113.6, 41.4 ppm; IR (solid) $v_{max}$ 3425, 3317, 3194, 3029, 2093, 1629, 1596, 1585, 1557, 1493, 1459, 1416, 1315, 1266, 1175, 1158, 1119, 997, 858, 809, 773, 752, 721, 693 $cm^{-1}$; LCMS (+ESI) *m/z* 185.2 $[M+H]^+$, 0.34 min, (100%); HRMS (+ESI) *m/z* (Calcd. $C_{12}H_{13}N_2$ $[M+H]^+$, 185.1073), Obs. 185.1071 ($\delta$ 1.2 ppm).

**N-(4-(Pyridin-4-ylmethyl)phenyl)acetamide, (3f)**

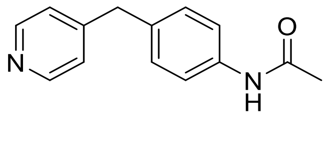

Acetic anhydride (115 uL, 1.1 mmol, 1.1 eq) was added to a stirred solution of 4-(4-aminophenyl)pyridine (184 mg, 1 mmol), and triethylamine (279 uL, 2 mmol) in DCM (5 mL), and the reaction was stirred overnight. Water

(10 mL) was added, and the product was extracted with DCM (3 x 10 mL), dried over anhydrous $Na_2SO_4$ and the solvent removed under reduced pressure. The crude material was purified by flash chromatography (0-5% MeOH in DCM) to yield **3f** as a white solid (193 mg, 0.85 mmol, 85%).

$^1$H-NMR (400 MHz, MeOD) δ 8.39 (m, 2H), 7.49 (m, 2H), 7.25 (dd, $J$ = 8.6, 6.3 Hz, 2H), 7.16 (d, $J$ = 8.7 Hz, 2H), 3.97 (s, 2H), 2.10 (s, 3H) ppm; $^{13}$C-NMR (100 MHz, MeOD) δ 171.6, 153.5, 149.9, 138.5, 136.1, 130.4, 125.8, 121.5, 41.3, 23.8 ppm; LCMS (+ESI) *m/z* 227.1 $[M+H]^+$, 0.70 min; (100%); HRMS (+ESI) *m/z* (Calcd. $C_{14}H_{15}N_2O$ $[M+H]^+$, 227.1179), Obs. 227.1178 ($\delta$ 0.6 ppm).

**N-(4-Pyridin-4-ylmethyl)phenyl)methanesulfonamide (3g)**

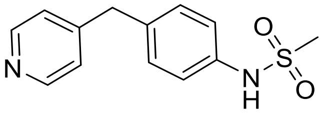


Methane sulfonyl chloride (85 uL, 1.1 mmol, 1.1 eq) was added to a stirred solution of 4-(4-aminophenyl)pyridine (184 mg, 1 mmol), and triethylamine (279 uL, 2 mmol) in DCM (5 mL), and the reaction was stirred overnight. Water (10 mL) was added, and the product was extracted with DCM (3 x 10 mL), dried over anhydrous $Na_2SO_4$ and the solvent removed under reduced pressure. The crude material was purified by flash chromatography (0-5% MeOH in DCM) to yield **3g** as an off-white solid (167 mg, 0.64 mmol, 64%). $^1$H-NMR (400 MHz, $d_6$-DMSO) δ 9.67 (s, 1H), 8.45 (m, 2H), 7.23 (m, 2H), 7.21 (d, $J$ = 8.9 Hz, 2H), 7.16 (d, $J$ = 8.7 Hz, 2H), 3.91 (s, 2H), 2.95 (s, 3H) ppm; $^{13}$C-NMR (100 MHz, $d_6$-DMSO) δ 150.1, 149.7, 136.7, 135.2, 129.8, 129.4, 124.1, 120.3, 39.5, 39.2 ppm; LCMS (+ESI) *m/z* 263.1 $[M+H]^+$, 0.70 min, (98%); HRMS (+ESI) *m/z* (Calcd. $C_{13}H_{15}N_2O_2S$ $[M+H]^+$, 263.0849), Obs. 263.0847 ($\delta$ 0.8 ppm).

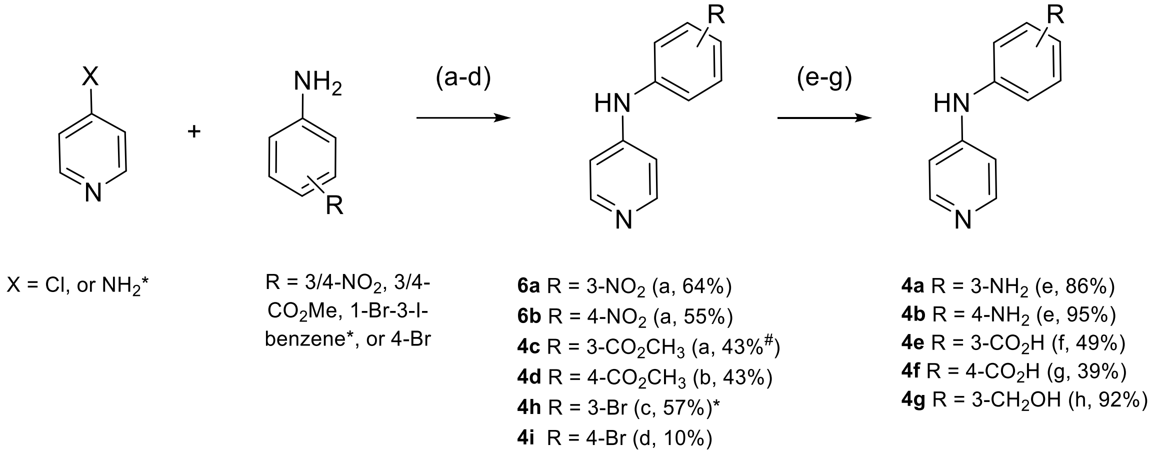


**S.I. Scheme 3.** Synthesis of **2a** analogues to explore SAR of "*linker*"*:* (a) X = Cl.xHCl, HCl (37%), EtOH, 90 °C, 20 h; (b) X = Cl.xHCl, R=4-$CO_2$Me, AcOH, LiOH.$H_2O$, MeOH:$H_2O$:THF, r.t., 4 h; (c) X = $NH_2$, 1-bromo-3-iodobenzene, $Pd_2(dba)_3$, DPPF, Na$^t$OBu, toluene, 115 °C, 24 h; (d) $Pd_2(dba)_3$, IPr.HCl, K$^t$OBu, 1,4-dioxane,

100 °C, 21 h; (e) **6a/b**, $SnCl_2.2H_2O$, HCl (37%), EtOH, 0-80 °C, 1−3 h; (f) **4c**, $LiOH.H_2O$, $MeOH:H_2O:THF$, r.t., 4 h; (g) **4d**, KOH(aq), EtOH, reflux, 2 h; (h) **4c**, $LiAlH_4$, THF, 0 °C-r.t., 20 h. #impure mixture of methyl/ethyl ester

**N-(3-Nitrophenyl)pyridin-4-amine (6a)**

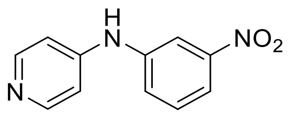


3-Nitroaniline (414 mg, 3.00 mmol) was added to a solution of 4-chloropyridine HCl (450 mg, 3.00 mmol) in EtOH (15 mL), followed by a catalytic amount of conc. HCl (37%, 4 drops). The reaction was heated at 90 °C for 24 hours, at which point a yellow precipitate had formed. The reaction was allowed to cool to room temperature, and then the precipitate was collected by vacuum filtration, washed with ice cold EtOH, and dried under reduced pressure to yield compound **6a** as a yellow solid (411 mg, 1.91 mmol, 64%), which was used without further purification. $R_f$ 0.02 (5% v/v MeOH in DCM); $^1$H-NMR (500 MHz, $d_6$-DMSO) $\delta$ 14.22 (s, 1H), 11.32 (s, 1H), 8.37 (d, $J$ = 7.4 Hz, 2H), 8.16 (app. t, $J$ = 2.1 Hz, 1H), 8.11 (ddd, $J$ = 8.2, 2.2, 1.0 Hz, 1H), 7.84 (ddd, $J$ = 8.0, 2.1, 0.9 Hz, 1H), 7.76 (app. t, $J$ = 8.1 Hz, 1H), 7.32 (d, $J$ = 7.5 Hz, 1H) ppm; $^{13}$C NMR (125 MHz, $d_6$-DMSO) $\delta$ 155.9, 148.6, 140.9, 138.8, 131.3, 129.0, 120.3, 117.2, 109.4 ppm; IR (solid) $v_{max}$ 3178-2700, 3066, 3036, 2872, 2831, 1646, 1610, 1584, 1563, 1523, 1479, 1355, 1342, 1317, 1304, 1243, 1233, 1202, 1103, 1003, 946, 898, 856, 840, 800, 791, 739, 700, 677 $cm^{-1}$; LCMS (+ESI) $m/z$ 216.1 $[M+H]^+$, retention time 1.29 min, (97%); HRMS (+ESI) $m/z$ (Calcd. $C_{11}H_{10}N_3O_2$ $[M+H]^+$, 216.0768), *Obs*. 216.0762 ($\delta$ 2.8 ppm).

**N-(4-Nitrophenyl)pyridin-4-amine (6b)**

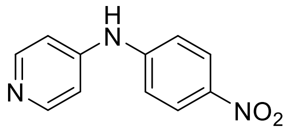


4-Nitroaniline (414 mg, 3.00 mmol) was added to a solution of 4-chloropyridine HCl (450 mg, 3.00 mmol) in EtOH (15 mL), followed by a catalytic amount of conc. HCl (37%, 4 drops). The reaction was heated at 90 °C for 24 hours, at which point a yellow precipitate had formed. The reaction was allowed to cool to room temperature, and then the precipitate was collected by vacuum filtration, washed with ice cold EtOH, and dried under reduced pressure to yield compound **6b** as a yellow solid ((352 mg, 1.64 mmol, 55%), which was used without further purification. $R_f$ 0.18 (10% v/v MeOH in EtOAc); $^1$H-NMR (500 MHz, $d_6$-DMSO) $\delta$ 14.40 (br s, 2H), 11.51 (s, 1H), 8.45 (d, $J$ = 7.4 Hz, 2H), 8.31 (d, $J$ = 9.0 Hz, 2H), 7.62 (d, $J$ = 9.1 Hz, 2H), 7.47 (d, $J$ = 7.4 Hz, 2H) ppm; $^{13}$C-NMR (125 MHz, $d_6$-DMSO) $\delta$ 155.0, 144.2, 143.6, 141.2, 125.5, 121.6, 110.6 ppm; IR (solid) $v_{max}$ 3076-2600, 3029, 2941, 1640, 1621, 1589, 1581, 1501, 1486, 1334, 1208, 1293, 1236, 1207, 1172, 1111,

1008, 853, 836, 810, 750, 736, 698, 631 cm$^{1}$; LCMS (+ESI) *m/z* 216.1 $[M+H]^+$, retention time 1.31 min, (100%); HRMS (+ESI) *m/z* (Calcd. $C_{11}H_{10}N_3O_2$ $[M+H]^+$, 216.0768), *Obs.* 216.0764 ($\delta$ 1.8 ppm).

**N$^1$-(Pyridin-4-yl)benzene-1,3-diamine (4a)**

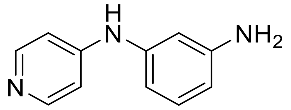

Tin(II) chloride dihydrate (1.08 g, 4.80 mmol) was added to a stirred solution of N-(3-nitrophenyl)pyridin-4-amine (**6a**) (207 mg, 0.96 mmol) in EtOH (6 mL). The reaction was cooled to 0 ˚C and concentrated HCl (37% v/v soln., 100 µL,) was added dropwise. The reaction was then heated under reflux for 3 hours, then allowed to cool to room temperature and quenched with 2 M $Na_2CO_3$ (~ 5 mL). The product was extracted into EtOAc (4 x 10 mL) the organic fractions were combined, washed with brine (5 mL), dried over anhydrous $Na_2SO_4$, and then the solvent was removed under reduced pressure to yield compound **4a** as a yellow solid (152 mg, 0.82 mmol, 86%). $^1$H-NMR (700 MHz, $d_6$-DMSO) $\delta$ 8.51 (s, 1H), 8.13 (d, $J$ = 6.5 Hz, 1H), 6.95 (app. t, $J$ = 7.9 Hz, 1H), 6.85 (m, 2H), 6.43 (app. t, $J$ = 2.1 Hz, 1H), 6.32 (ddd, $J$ = 7.8, 2.0, 0.7 Hz, 1H), 6.25 (ddd, $J$ = 8.0, 2.1, 0.9 Hz, 1H). ppm; $^{13}$C-NMR (175 MHz, $d_6$-DMSO) $\delta$ 150.4, 149.9, 149.7, 141.0, 129.6, 109.2, 108.9, 108.0, 105.5 ppm; IR (solid) $\nu_{max}$ 3455, 3372, 3199-2700, 3055, 2920, 2892, 2843, 2811, 1606, 1579, 1524, 1484, 1445, 1347, 1300, 1242, 1215, 1182, 1160, 1098, 1054, 992, 965, 841, 826, 779, 695, 657 cm$^{-1}$; LCMS (+ESI) *m/z* 186.2 $[M+H]^+$, retention time 0.46 min, (100%); HRMS (+ESI) *m/z* (Calcd. $C_{11}H_{12}N_3$ $[M+H]^+$, 186.1026), *Obs.* 186.1022 ($\delta$ 1.9 ppm).

**N$^1$-(Pyridin-4-yl)benzene-1,4-diamine (4b)**

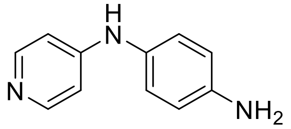

Tin(II) chloride dihydrate (0.56 g, 2.5 mmol) was added to a stirred solution of N-(4-nitrophenyl)pyridin-4-amine (**6b**) (108 mg, 0.50 mmol) in EtOH (3 mL). The reaction was cooled to 0 °C and concentrated HCl (37% v/v soln., 100 µL,) was added dropwise. The reaction was then heated under reflux for 1 hour, at which point all starting material had been consumed and an orange precipitate formed. The reaction was cooled to room temperature, quenched with 2 M $Na_2CO_3$ (~ 5 mL), and the product was extracted into EtOAc (4 x 5 mL). The organic fractions were combined, washed with brine (2 mL), dried over anhydrous $Na_2SO_4$, and the solvent was removed under reduced pressure to yield compound **4b** as a yellow solid (88 mg, 0.48 mmol, 95%). $^1$H-NMR (500 MHz, $d_6$-DMSO) $\delta$ 8.21 (s, 1H), 8.03 (d, $J$ = 6.5 Hz, 2H), 6.86 (d, $J$ = 8.5 Hz, 2H), 6.63–6.53 (m, 4H), 4.97 (br s, 2H) ppm; $^{13}$C-NMR (125 MHz, $d_6$-DMSO) $\delta$ 152.3, 149.7, 145.6, 128.3, 124.4, 114.5, 107.8 ppm; IR (solid) $v_{max}$ 3429, 3379, 3301, 3140, 3029, 1643, 1594, 1570, 1506, 1435, 1411, 1345, 1325, 1282, 1216, 1173, 991, 885, 807, 648, 613 $cm^{-1}$; LCMS (+ESI) *m/z* 186.2 $[M+H]^+$, retention time 0.31 min, (100%); HRMS (+ESI) *m/z* (Calcd. $C_{11}H_{12}N_3$ $[M+H]^+$, 186.1026), *Obs*. 186.1018 ($\delta$ 4.8 ppm).

**Methyl/Ethyl 3-(pyridin-4-ylamino)benzoate (4c)**

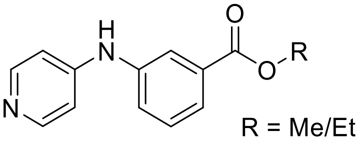


Methyl 3-aminobenzoate (453 mg, 3.00 mmol) was added to a solution of 4-chloropyridine hydrochloride (450 mg, 3.00 mmol) in EtOH (15 mL). A catalytic amount of conc. HCl (37%, 4 drops) was added and the reaction was heated at 90 °C for 24 hours and then the solvent was removed under reduced pressure. The resulting residue was redissolved in DCM (60 mL), washed with saturated $NaHCO_3$ (15 mL), and brine (10 mL), dried over anhydrous $Na_2SO_4$ and the solvent was removed under reduced pressure. The crude material was purified by flash chromatography (0−10% v/v MeOH in DCM) to yield **4c**, which was a mixture of the methyl and ethyl 3-(pyridin-4-ylamino)benzoates in 1.7:1.3 ratio by $^1$H NMR integration, respectively, as a pink amorphous solid (298 mg, 1.3 mmol, 24/18%). $R_f$ 0.14 (10% v/v MeOH in DCM); $^1$H-NMR (500 MHz, $CDCl_3$) $\delta$ 8.32 (d, $J$ = 5.0 Hz, 2H), 7.86 (d, $J$ = 1.4 Hz, 1H), 7.78 (m, 1H), 7.43 (app. t, $J$ = 7.7 Hz, 1H), 7.39 (d, $J$ = 7.7 Hz, 1H), 6.84 (d, $J$ = 4.9 Hz, 2H), 6.39 (d, $J$ = 5.3 Hz, 1H), 4.38 (q, $J$ = 7.1 Hz, 1H), 3.92 (s, 2H), 1.39 (t, $J$ = 7.1 Hz, 1H) ppm; $^{13}$C-NMR (125 MHz, $CDCl_3$) $\delta$ 166.7, 166.2, 150.7, 150.2, 150.1, 140.2, 140.1, 132.2, 131.8, 129.8, 129.7, 125.6, 125.5, 125.1, 122.3, 122.2, 109.9, 109.8, 61.4, 52.5, 14.5 ppm; IR (solid) $v_{max}$ 3164-2800, 3058, 2916, 2819, 1717, 1625, 1587, 1569, 1527, 1500, 1479, 1435, 1418, 1345, 1294, 1266, 1214, 1160, 1102, 1077, 994, 813, 749, 690, 670 $cm^{-1}$; LCMS (+ESI) *m/z* 229.0, 243.0 $[M+H]^+$, retention time 1.38, 1.53 min, (55%, 45%).

**Methyl 4-(pyridin-4-ylamino)benzoate (4d)**

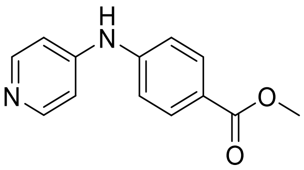


Methyl 4-aminobenzoate (252 mg, 1.67 mmol) was added to a stirred solution of 4-chloropyridine hydrochloride (250 mg, 1.67 mmol) in glacial AcOH (2.5 mL). The reaction was heated at 100 °C for 20 hours and then concentrated under reduced pressure. The resulting residue was resuspended in water (2.5 mL) and brought to pH 10 using 6 M NaOH. The product was extracted into $CHCl_3$ (4 x 4 mL), concentrated and then purified by flash chromatography (0−5% v/v MeOH in DCM) to yield compound **4d** as a white solid (164 mg, 0.72 mmol, 43%). $^1$H-NMR (400 MHz, MeOD) $\delta$ 8.22 (dd, $J$ = 7.2, 1.3 Hz, 2H), 7.99 (dd, $J$ = 6.8, 2.0 Hz, 2H), 7.29 (dd, $J$

= 6.8, 2.0 Hz, 2H), 7.10 (dd, *J* = 5.0, 1.6 Hz, 2H), 3.88 (s, 3H) ppm; $^{13}$C NMR (100 MHz, MeOD) $\delta$ 168.4, 152.2, 150.2, 146.6, 132.4, 125.2, 119.8, 112.0, 52.6 ppm; IR (solid) $\nu_{max}$ 2920, 1709, 1638, 1583, 1522, 1502, 1431, 1345, 1267, 1104, 998, 818 cm$^{-1}$; LCMS (+ESI) *m/z* 229.0 [M+H]$^+$; HRMS (+ESI) *m/z* (Calcd. $C_{13}H_{13}N_2O_2$ [M+H]$^+$, 229.0972), *Obs*. 229.0961 ($\delta$ 4.8 ppm).

**3-(Pyridin-4-ylamino)benzoic acid (4e)**

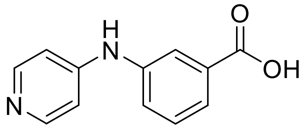


Lithium hydroxide monohydrate (319 mg, 7.6 mmol) was added to a solution of ester **4c** (288 mg, approximately 55:45 methyl/ethyl ester, ~1.27 mmol) in THF:MeOH:$H_2O$ (2:1:1) (6 mL) and the reaction was stirred at room temperature for 4 hours. The reaction was then concentrated under reduced pressure and the residue was redissolved in water (75 mL) and brought to pH ~1-2 using 3 M HCl. The product was extracted into a mixture of $CHCl_3$ and *i*-PrOH (2:1, 20 x 30 mL). The organic fractions were combined and concentrated under reduced pressure to yield **4e** as a pink amorphous solid (133 mg, 0.62 mmol, 49%). $^1$H-NMR (400 MHz, MeOD) $\delta$ 8.21 (d, *J* = 7.5 Hz, 2H), 8.01–7.97 (m, 2H), 7.65–7.58 (m, 2H), 7.16 (d, *J* = 7.5 Hz, 2H) ppm; $^{13}$C-NMR (100 MHz, MeOD) $\delta$ 168.6, 158.7, 141.7, 138.7, 134.1, 131.4, 129.2, 129.2, 125.8, 125.6 ppm; IR (solid) $\nu_{max}$ 3500-2800, 3404, 3228, 3086, 2981, 1720, 1644, 1586, 1571, 1524, 1493, 1476, 1430, 1384, 1306, 1270, 1255, 1212, 1201, 1110, 1003, 934, 900, 822, 791, 747, 688, 663 cm$^{-1}$; LCMS (+ESI) m/z 215.0, 215.0 [M+H]+, retention time 0.31, 1.19 min, (17%, 83%); HRMS (-ESI) m/z (Calcd. $C_{12}H_9N_2O_2$ [M-H]$^-$, 213.0664), *Obs*. 213.0674 ($\delta$ 2.3 ppm).

**4-(Pyridin-4-ylamino)benzoic acid (4f)**

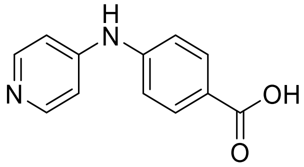


A solution of potassium hydroxide (15 mg, 0.26 mmol) in water (1.4 mL) was added to a stirred solution of ester **4d** (30 mg, 0.13 mmol) in EtOH (0.7 mL) and the reaction was heated under reflux for 2 hours. The reaction was then concentrated under reduced pressure brought to pH 2 using 2 M HCl. The mixture was washed with $CHCl_3$ (8 x 25 mL) and then the aqueous fraction was concentrated under reduced pressure. The residue was redissolved in water (15 mL), brought to pH 4 and the product was extracted using EtOAc (15 x 20 mL) and then $CHCl_3$/***i***-PrOH (2:1 v/v, 5 x 15 mL). The organic fractions were combined and the solvent was removed under reduced pressure to yield compound **4f** as a white solid (11 mg, 0.05 mmol, 39%). $^1$H-NMR (400 MHz, MeOD) $\delta$ 8.22

(d, *J* = 6.3 Hz, 2H), 8.09 (d, *J* = 8.6 Hz, 2H), 7.39 (*J* = 8.6 Hz, 2H), 7.21 (d, *J* = 6.3 Hz, 2H) ppm; $^{13}$C-NMR (100 MHz, MeOD) $\delta$ 171.2, 156.9, 142.0, 141.2, 131.1, 121.4, 109.5, 93.9 ppm; IR (solid) $\nu_{max}$ 2920, 2820, 1708, 1647, 1603, 1531, 1493, 1476, 1430, 1384, 1306, 1270, 1255, 1212 cm$^{-1}$; LCMS (+ESI) m/z 215.2 [M+H]$^+$; HRMS (-ESI) m/z (Calcd. $C_{12}H_9N_2O_2$ [M+H]$^+$, 215.0815), *Obs*. 215.0818 ($\delta$ 1.4 ppm).

**(3-Pyridin-4-ylamino)phenyl)methanol (4g)**

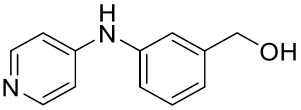


$LiAlH_4$ (10 mg, 0.26 mmol) was added slowly, under an inert atmosphere to a stirred solution of ester **4c** (30 mg, 0.13 mmol) in anhydrous THF (0.7 mL) at 0 °C. When the addition was complete, the reaction was allowed to warm slowly to room temperature and stirred for 16 hours. As starting material remained, additional $LiAlH_4$ (25 mg, 0.66 mmol) in THF (1.5 mL) was added and the reaction was stirred for a further 4 hours at room temperature before being quenched with water (0.5 mL), followed by 4 M NaOH (0.5 mL), and additional water (1.5 mL). The reaction was then filtered through celite, eluting the product with DCM (1 mL). The organic solvent was collected and concentrated under reduced pressure to yield compound **4g** as a yellow solid (24 mg, 0.12 mmol, 92%). $^1$H-NMR (400 MHz, MeOD) $\delta$ 8.10 (d, *J* = 5.4 Hz, 2H), 7.34 (app.t, *J* = 7.8 Hz, 2H), 7.25 (s, 1H), 7.09 (d, *J* = 7.8 Hz, 1H), 7.12 (d, *J* = 7.8 Hz, 1H), 6.93 (d, *J* = 5.4 Hz, 2H), 4.61 (s, 2H) ppm; $^{13}$C-NMR (100 MHz, MeOD) $\delta$ 153.7, 150.0, 144.5, 141.6, 130.6, 123.5, 121.4, 121.0, 110.3, 65.1 ppm; IR (solid) $\nu_{max}$ 3264, 3059, 1615, 1591, 1520, 1474, 1345, 1030, 1000, 817 cm$^{-1}$; LCMS (+ESI) *m/z* 201.2 [M+H]$^+$; HRMS (+ESI) *m/z* (Calcd. $C_{12}H_{13}N_2O$ [M+H]$^+$, 201.1028), *Obs*. 201.1025 ($\delta$ 1.5 ppm).

**N-(3-Bromophenyl)pyridin-4-amine (4h)**

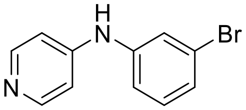


4-Aminopyridine (500 mg, 5.31 mmol), 1-bromo-3-iodobenzene (0.75 mg, 5.8 mmol), NaO$^t$Bu (608 mg, 16.32 mmol), $Pd_2(dba)_3$ (73 mg, 0.19 mmol), and DPPF (107 mg, 0.19 mmol) were combined in anhydrous toluene (16 mL) and heated under reflux (115 °C) for 24 hours. The reaction was then allowed to cool to room temperature, diluted with $Et_2O$ (50 mL) and filtered through a short plug of celite. The filtrate was concentrated under reduced pressure and the crude material was purified by flash chromatography (0-5% v/v MeOH in DCM), to yield compound **4h** as a brown solid (755 mg, 3.0 mmol, 57%). $R_f$ 0.07 (10% v/v MeOH in DCM); $^1$H-NMR (400 MHz, MeOD) $\delta$ 8.16 (d, *J* = 5.1 Hz, 2H), 7.37 (dd, *J* = 1.9, 1.9 Hz, 1H), 7.29-7.19 (m, 3H), 6.95 (d, *J* = 5.1 Hz, 2H), 6.07 (br, s, 1H) ppm; $^{13}$C-NMR (100 MHz, MeOD) $\delta$ 152.8, 150.5, 143.4, 132.1, 127.4, 124.0, 120.5

(2C), 111.0 ppm; IR (solid) $v_{max}$ 2913, 1606, 1580, 1520, 1469, 1346, 1215, 993, 808 cm$^{-1}$; HRMS (+ESI) *m/z* (Calcd. $C_{11}H_{10}BrN_2$ [M+H]$^+$, 249.0022), *Obs.* 249.0017 ($\delta$ 2.0 ppm).

**N-(4-Bromophenyl)pyridin-4-amine (4i)**

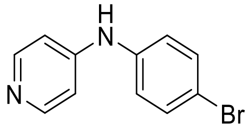


4-Chloropyrine hydrochloride (500 mg, 3.33 mmol) was desalted using 6 M NaOH and extracted into $Et_2O$ (4 x 15 mL). The solution was dried over anhydrous $Na_2SO_4$ and then the solvent was removed under reduced pressure to yield a solid product. Desalted 4-chloropyridine (113 mg, 1 mmol), KO$^t$Bu (168 mg, 1.5 mmol), $Pd_2(dba)_3$ (18 mg, 0.02 mmol), 1,3-bis(2,6-diisopropylphenyl)imidazol-2-ylidene hydrochloride (9 mg, 0.02 mmol), and 4-bromoaniline (189 mg, 1.1 mmol) were combined in 1,4-dioxane (3.1 mL) heated under an inert atmosphere. The reaction was heated at 100 °C for 21 hours, then cooled to room temperature and diluted with 1 M HCl (7 mL). The solution was washed with $E_2O$ (2 x 4 mL) and then the aqueous phase was brought to pH 14 using 6 M NaOH. The product was extracted into $CHCl_3$ (3 x 4 mL), the organic fractions were combined, washed with brine (4 mL), dried over $Na_2SO_4$ and the solvent was removed under reduced pressure. The crude product was purified by flash chromatography (2:3 v/v DCM:EtOAC) to yield compound **4i** as a brown solid (25 mg, 0.10 mmol, 10%). $^1$H-NMR (500 MHz, $CDCl_3$) $\delta$ 8.29 (dd, *J* = 4.8, 1.6 Hz, 2H), 7.45 (dd, *J* = 6.6, 2.2 Hz, 2H), 7.05 (dd, *J* = 6.6, 2.2 Hz, 2H), 6.77 (dd, *J* = 4.8, 1.6 Hz, 2H), 5.93 (s, 1H) ppm; $^{13}$C-NMR (125 MHz, $CDCl_3$) $\delta$ 150.9, 150.1, 138.9, 132.8, 123.3, 116.9, 109.9 ppm; IR (solid) $v_{max}$ 2902, 1739, 1636, 1603, 1576, 1524, 1484, 1430, 1346, 1217, 1071, 996 cm$^{-1}$; LCMS (+ESI) *m/z* 248.9 [M+H]$^+$; HRMS (+ESI) *m/z* (Calcd. $C_{11}H_{10}BrN_2$ [M+H]$^+$, 249.0022), *Obs.* 249.0018 ($\delta$ 1.6 ppm).

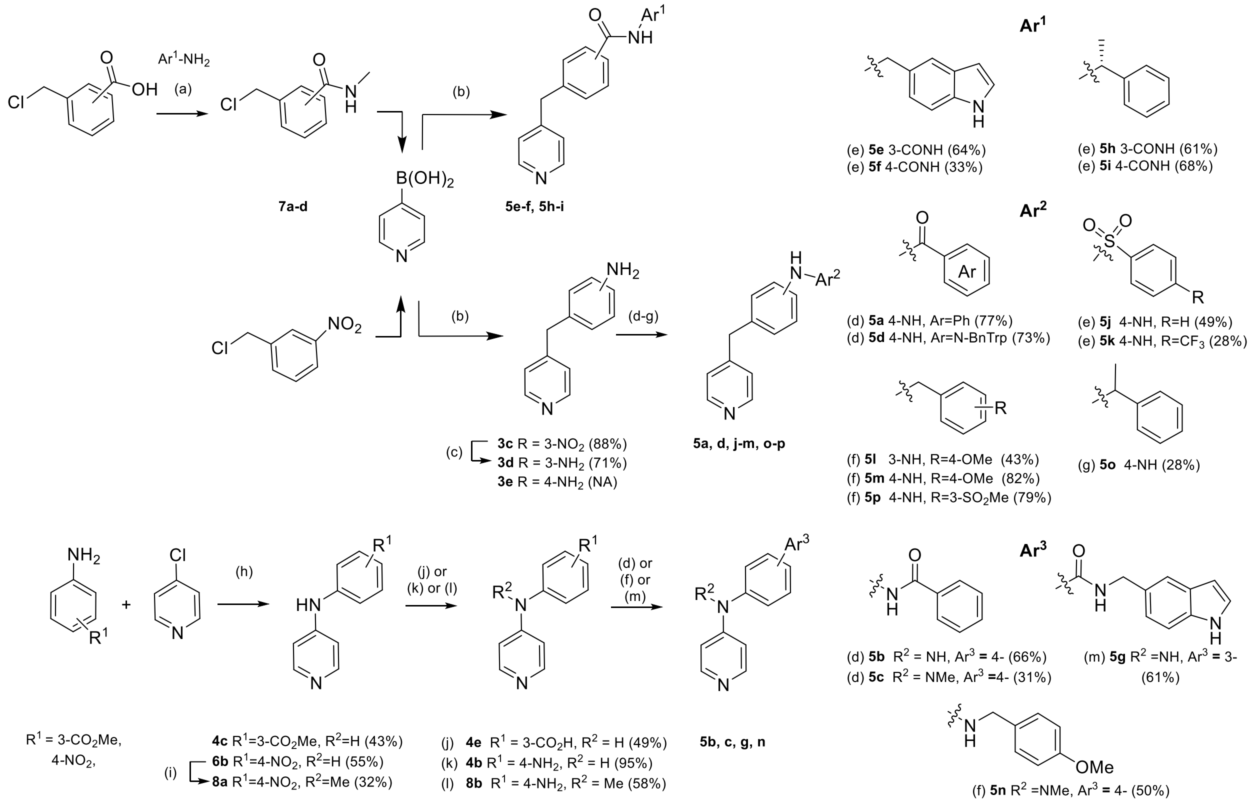


**Scheme 1.** (a) HBTU, $Et_3N$, DMC:DMF (7:1), r.t., 24 h; (b) $Pd(PPh_3)_4$, $Na_2CO_3$, DME:$H_2O$ (2:1), 100 °C, 4 h; (c) Pd/C, $N_2H_4$.x$H_2O$, EtOH, 90 °C, 2 h; (d) $ArCO_2H$, HATU, DIPEA, DCM, 0 °C−r.t., 24 h (**5a**, **5b**, **5c**); or PyBOP, NMM, DCM, DMF, r.t., 5 h (**5d**); (e) $ArSO_2Cl$, pyridine, r.t., 20 h (**5j**); or $ArSO_2Cl$, $Et_3N$, DCM, r.t., 20 h (**5k**); (f) RCOH, AcOH, $NaCNBH_3$, MeOH, r.t., 20 h (**5l**, **5m**, **5p**, **5n**); (g) RCOMe, $TiCl_4$, DCM, 0 °C, 3 h; then $Na(CN)BH_3$, MeOH, r.t., 24 h (**5o**); (h) HCl (37%), EtOH, 90 °C, 20 h; (i) NaH, DMF, MeI, 0 °C- r.t., 8 h; (j) **4c**, LiOH.$H_2O$, MeOH:$H_2O$:THF, r.t., 4 h; (k) **6b**, $SnCl_2$.2$H_2O$, HCl (37%), EtOH, 0-80 °C, 1 h; (l) **8a**, Zn(s), $NH_4Cl$, DMF, r.t., 24 h; (m) EDC.HCl, HOAt, DIPEA, DMF:DCM (1:10).

**N-((1H-Indol-5-yl)methyl)-3-(chloromethyl)benzamide (7a)**

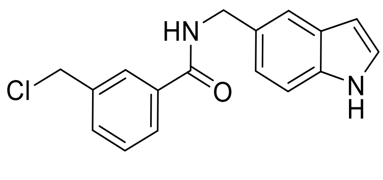

HBTU (455 mg, 1.20 mmol) was added to a solution of 3-(chloromethyl)benzoic acid (225 mg, 1.32 mmol) in anhydrous DCM (1 mL) and anhydrous DMF (250 µL) and stirred for 10 minutes at room temperature. Then, a solution of 5-(aminomethyl)indole (153 mg, 1.2 mmol) and $Et_3N$ (250 µL, 1.8 mmol) in anhydrous DCM (1 mL) was added slowly, and the reaction was stirred at room temperature for 24 hours. The reaction was then diluted with EtOAc (50 mL), washed with saturated citric acid (6 mL), and saturated $NaHCO_3$ (10 mL) until basic, followed by water (3 x 3 mL), brine (3 mL), dried over anhydrous $Na_2SO_4$, and the solvent was removed under reduced pressure. The crude product was purified twice by flash chromatography (0-50% v/v EtOAc in PE), to yield compound **7a** as a white solid (202 mg, 0.68 mmol, 56%). $R_f$ 0.37 (50% v/v EtOAc in PE);$^1$H-NMR (500 MHz, $CDCl_3$) $\delta$ 8.27 (br s, 1H), 7.81 (td, $J$ = 1.9, 0.5 Hz, 1H), 7.72 (ddd, $J$ = 7.7, 1.8, 1.2 Hz, 1H), 7.64 (dt, $J$ = 1.6, 0.8 Hz, 1H), 7.52 (dtd, $J$ = 7.7, 1.3, 0.6 Hz, 1H), 7.43 (dd, $J$ = 7.7, 0.5 Hz, 1H), 7.40 (dt, $J$ = 8.3, 0.8 Hz, 1H), 7.24 (dd, $J$ = 3.2, 2.4 Hz, 1H), 7.21 (dd, $J$ = 8.4, 1.7 Hz, 1H), 6.55 (ddd, $J$ = 3.1, 2.0, 0.9 Hz, 1H), 6.37 (br s, 1H), 4.74 (d, $J$ = 5.3 Hz, 2H), 4.60 (s, 2H) ppm; $^{13}$C-NMR (125 MHz, $CDCl_3$) $\delta$ 166.7, 138.2, 135.5, 135.3, 131.6, 129.4, 129.2, 128.3, 127.4, 127.0, 125.1, 122.6, 120.6, 111.6, 102.8, 45.7, 45.1 ppm; IR (solid) $v_{max}$ 3271, 2922, 2547, 1633, 1605, 1581, 1532, 1482, 1427, 1355, 1298, 1263, 1216, 1093, 1052, 1000, 876, 805, 753, 704 cm$^{-1}$; LCMS (+ESI) *m/z* 299.2 [M+H]$^+$, retention time 1.96 min, (100%); HRMS (+ESI) *m/z* (Calcd. $C_{17}H_{15}N_2OClNa$ [M+Na]$^+$, 321.0765), *Obs*. 321.0763 ($\delta$ 0.6 ppm).

**N-(($^1$H-Indol-5-yl)methyl)-4-(chloromethyl)benzamide (7b))**

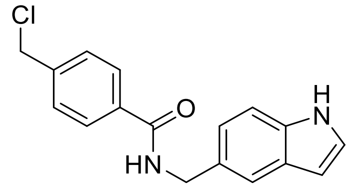


HBTU (228 mg, 0.60 mmol) was added to a stirred solution of 4-(chloromethyl)benzoic acid (113 mg, 0.66 mmol) in anhydrous DMF (700 µL). After 10 minutes, a solution of 5-(aminomethyl)indole (88 mg, 0.60 mmol) and $Et_3N$ (125 µL, 0.9 mmol) in anhydrous DMF (300 µL) was added to the stirred reaction over period of 1-2 minutes. The reaction was stirred at room temperature for 24 hours then diluted with EtOAc (25 mL), washed with saturated citric acid (3 mL), saturated $NaHCO_3$ (10 mL), dried over anhydrous $Na_2SO_4$ and the solvent was removed under reduced pressure. The crude product was purified twice by flash chromatography (0-50% v/v EtOAc in PE), then redissolved in EtOAc (50 mL) and washed with water (4 x 5 mL), and brine (5 mL) to remove residual DMF. The solvent was removed under reduced pressure to yield compound **7b** as an off-white solid (81 mg, 0.27 mmol, 45%). $R_f$ 0.29 (1:2 v/v EtOAc:PE); $^1$H-NMR (500 MHz, $CDCl_3$) $\delta$ 8.25 (br s, 1H), 7.78 (d, $J$ = 8.3 Hz, 2H), 7.63 (s, 1H), 7.44 (d, $J$ = 8.3 Hz, 2H), 7.39 (d, $J$ = 8.3 Hz, 1H), 7.24 (app. t, $J$ = 2.8 Hz, 1H), 7.20 (dd, $J$ = 8.3, 1.6 Hz, 1H), 6.54 (ddd, $J$ = 3.1, 2.0, 1.0 Hz, 1H), 6.35 (br s, 1H), 4.73 (d, $J$ = 5.4 Hz, 2H), 4.60 (s, 2H) ppm; $^{13}$C-NMR (125 MHz, $CDCl_3$) $\delta$ 166.7, 140.9, 135.5, 134.7, 129.4, 128.8, 128.3, 127.5, 125.1, 122.6, 120.5, 111.6, 102.8, 45.5, 45.0 ppm; IR (solid) $v_{max}$ 3332, 2925, 1620, 1570, 1543, 1505, 1465, 1423, 1359, 1329, 1303, 1269, 1185, 1095, 1033 987, 867, 852, 752, 723, 677 cm$^{-1}$; LCMS (+ESI) *m/z* 299.2 [M+H]$^+$, 1.92 min, (98%); HRMS (+ESI) *m/z* (Calcd. $C_{17}H_{16}N_2OCl$ [M+H]$^+$, 299.0946), *Obs*. 299.0941 ($\delta$ 1.4 ppm).

**(R)-3-(Chloromethyl)-N-(1-phenylethyl)benzamide (7c)**

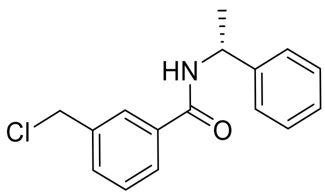

HBTU (455 mg, 1.20 mmol) was added to a solution of 3-(chloromethyl)benzoic acid (225 mg, 1.32 mmol) in anhydrous DCM (1 mL) and anhydrous DMF (250 µL) and stirred for 10 minutes at room temperature. Then, a solution of *(R)*-(+)-alpha-methylbenzylamine (153 µL, 1.2 mmol) and $Et_3N$ (250 µL, 1.8 mmol) in anhydrous DCM (1 mL) was added slowly, and the reaction was stirred at room temperature for 24 hours. The reaction was then diluted with EtOAc (50 mL), washed with saturated citric acid (6 mL), and saturated $NaHCO_3$ (10 mL) until basic, followed by water (3 x 3 mL), brine (3 mL), dried over anhydrous $Na_2SO_4$, and the solvent was removed under reduced pressure. The crude product was purified by flash chromatography (0-40% v/v EtOAc in PE), to yield compound **7c** as a white solid (194 mg, 0.71 mmol, 59%). $R_f$ 0.45 (50% v/v EtOAc in PE); $^1$H-NMR (500 MHz, $CDCl_3$) $\delta$ 7.79 (app. t, $J$ = 1.8 Hz, 1H), 7.71 (dt, $J$ = 7.8, 1.5 Hz, 1H), 7.53 (dt, $J$ = 7.7, 1.5 Hz, 1H), 7.43 (d, $J$ = 7.7 Hz, 1H), 7.41–7.39 (m, 2H), 7.38–7.35 (m, 2H), 7.29 (tt, $J$ = 7.2, 1.7 Hz, 1H), 6.34 (d, $J$ = 7.7 Hz, 1H), 5.34 (m, 1H), 1.62 (d, $J$ = 6.9 Hz, 3H) ppm; $^{13}$C-NMR (125 MHz, $CDCl_3$) $\delta$ 166.1, 143.1, 138.2, 135.3, 131.7, 129.2, 128.9, 127.7, 127.3, 126.9, 126.4, 49.5, 45.7, 21.8 ppm; IR (solid) $\nu_{max}$ 3309, 3060, 2978, 2931, 2874, 1635, 1603, 1589, 1536, 1494, 1446, 1322, 1281, 1260, 1221, 1140, 1094, 1082, 1015, 897, 824, 761, 701, 666 cm$^{-1}$; LCMS (+ESI) *m/z* 296.0 [M+Na]$^+$, 2.08 min, (100%); HRMS (+ESI) *m/z* (Calcd. $C_{16}H_{17}NOCl$ [M+H]$^+$, 274.0993), *Obs.* 274.0982 ($\delta$ 4.2 ppm).

**(R)-4-(Chloromethyl)-N-(1-phenylethyl)benzamide (7d)**

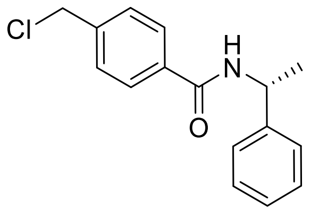

HBTU (455 mg, 1.20 mmol) was added to a solution of 4-(chloromethyl)benzoic acid (225 mg, 1.32 mmol) in anhydrous DCM (1 mL) and anhydrous DMF (250 µL) and stirred for 10 minutes at room temperature. Then, a solution of *(R)*-(+)-alpha-methylbenzylamine (153 µL, 1.2 mmol) and $Et_3N$ (250 µL, 1.8 mmol) in anhydrous DCM (1 mL) was added slowly, and the reaction was stirred at room temperature for 24 hours. The reaction was then diluted with EtOAc (50 mL), washed with saturated citric acid (6 mL), and saturated $NaHCO_3$ (10 mL) until basic, followed by water (3 x 3 mL), brine (3 mL), dried over anhydrous $Na_2SO_4$, and the solvent was removed under reduced pressure. The crude product was purified by flash chromatography (0-40% v/v EtOAc in PE), to yield compound **7d** as a white solid (195 mg, 0.71 mmol, 59%). $R_f$ 0.55 (50% v/v EtOAc in PE); $^1$H NMR (500 MHz, $CDCl_3$) $\delta$ 7.76 (d, $J$ = 8.3 Hz, 2H), 7.44 (d, $J$ = 8.4 Hz, 2H), 7.38–7.35 (m, 4H), 7.29 (tt, $J$ =

7.0, 6.7, 1.6 Hz, 1H), 6.31 (d, $J$ = 7.8 Hz, 1H), 5.34 (m, 1H), 4.60 (s, 2H), 1.61 (d, $J$ = 6.9 Hz, 3H) ppm; $^{13}$C-NMR (125 MHz, $CDCl_3$) $\delta$ 166.1, 143.1, 141.0, 134.7, 128.9, 128.8, 127.7, 127.5, 126.4, 49.4, 45.5, 21.8 ppm; IR (solid) $v_{max}$ 3334, 3035, 2980, 1627, 1571, 1528, 1502, 1495, 1450, 1322, 1279, 1210, 1149, 1123, 1087, 1013, 910, 876, 832, 815, 764, 702 cm$^{-1}$; LCMS (+ESI) *m/z* 274.2 $[M+H]^+$, retention time 2.10 min, (100%); HRMS (+ESI) *m/z* (Calcd. $C_{16}H_{16}NOClNa$ $[M+Na]^+$, 296.0813), *Obs*. 296.0811 ($\delta$ 0.7 ppm).

**N-Methyl-N-(4-nitrophenyl)pyridin-4-amine (8a)**

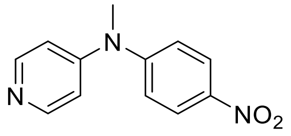

Sodium hydride (48 mg, 1.2 mmol, 60% dispersion in mineral oil) was added as a single portion to a solution of *N*-(4-nitrophenyl)pyridin-4-amine 6b (107 mg, 0.50 mmol) in anhydrous DMF (2.5 mL) at 0 °C. The reaction was stirred at room temperature for 45 minutes, then cooled to 0 °C and methyl iodide (74 µL, 1.2 mmol) was added dropwise over 2 minutes. The reaction was then stirred for 8 hours at room temperature and then quenched with water (5 mL). The product was extracted into EtOAc (3 x 10 mL) and organic fractions were combined, washed with brine (10 mL), dried over anhydrous $Na_2SO_4$ and then the solvent was removed under reduced pressure. The crude product was purified by flash chromatography (0-10% v/v MeOH in DCM) to yield compound 8a as a brown solid (36 mg, 0.16 mmol, 32%). $R_f$ 0.37 (10% v/v MeOH in DCM); $^1$H-NMR (500 MHz, MeOD) $\delta$ 8.34–8.17 (m, 4H), 7.44 (d, $J$ = 9.2 Hz, 2H), 7.01 (d, $J$ = 6.6 Hz, 2H), 3.48 (s, 3H) ppm; $^{13}$C-NMR (125 MHz, MeOD) $\delta$ 155.2, 153.2, 150.6, 145.1, 126.4, 124.8, 113.3, 39.8 ppm; IR (solid) $v_{max}$ 3019, 1574, 1491, 1418, 1361, 1333, 1318, 1305, 1225, 1195, 1148, 1112, 1092, 1062, 996, 878, 851, 826, 746, 732, 695 cm$^{-1}$; LCMS (+ESI) *m/z* 230.1, retention time 1.71 min, (97%); HRMS (+ESI) *m/z* (Calcd. $C_{12}H_{12}N_3O_2$ $[M+H]^+$, 230.0924), *Obs.* 230.0916 ($\delta$ 3.6 ppm).

**N$^1$-Methyl-N$^1$-(pyridin-4-yl)benzene-1,4-diamine (8b)**

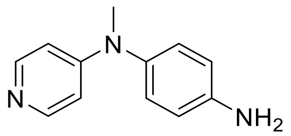

Ammonium chloride (361 mg, 6.75 mmol) and Zn(s) (441 mg, 6.75 mmol) were added to a solution of **8a** (62 mg, 0.27 mmol) in anhydrous DMF (4 mL) and the reaction was stirred at room temperature for 24 hours. The reaction was then diluted with EtOAc (25 mL) and filtered through a plug of celite, eluting with EtOAc (100 mL). The filtrate was washed with brine (3 x 25 mL), dried over $Na_2SO_4$, and the solvent was removed under reduced pressure. The residue was redissolved in EtOAc (75 mL), washed with water (3 x 20 mL), brine (15 mL), dried over $Na_2SO_4$, and the solvent was removed under reduced pressure to yield **8b** as a brown solid (31 mg, 0.16 mmol, 58%). $R_f$ 0.08 (20% v/v MeOH in EtOAc); $^1$H-NMR (400 MHz, $d_6$-DMSO) $\delta$ 8.04 (d, $J$ = 4.5 Hz, 2H), 6.87 (d, $J$ = 8.6 Hz, 2H), 6.62 (d, $J$ = 8.6 Hz, 2H), 6.45 (d, $J$ = 6.1 Hz, 2H), 5.20 (s, 2H), 3.16 (s, 3H) ppm; $^{13}$C-NMR (100 MHz, $d_6$-DMSO) $\delta$ 154.2, 149.1, 147.5, 134.0, 127.5, 114.8, 107.5, 39.4 ppm; IR (solid) $\nu_{max}$ 3324, 3182, 1635, 1595, 1536, 1506, 1469, 1371, 1297, 1287, 1244, 1223, 1168, 1130, 1071, 987, 877, 831, 802, 727, 699, 668 cm$^{-1}$; LCMS (+ESI) *m/z* 200.0, retention time 1.22 min, (97%); HRMS (+ESI) *m/z* (Calcd. $C_{12}H_{14}N_3$ [M+H]$^+$, 200.1188), *Obs.* 200.1190 ($\delta$ 1.0 ppm).

**N-((1H-Indol-5-yl)methyl)-3-(pyridin-4-ylmethyl)benzamide, (5e)**

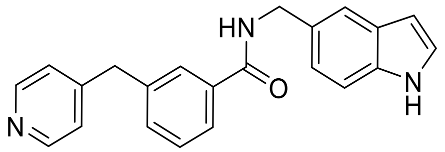


4-Pyridinylboronic acid (49 mg, 0.40 mmol), benzyl chloride **7a** (100 mg, 0.33 mmol), and $Na_2CO_3$ (74 mg, 0.70 mmol) were combined in a microwave vessel and flushed with $N_2$(g) for 5 minutes before the addition of $Pd(PPh_3)_4$ (8 mg, 0.01 mmol). The reaction vessel was flushed with $N_2$(g) for a further 2-3 minutes, and then a mixture of DME and water (2:1 v/v, 2 mL) was added, and the reaction was heated at 100 °C for 4 h. The reaction was then allowed to cool to room temperature, and diluted with water (5 mL) and DCM (5 mL). The phases were separated, and the aqueous fraction was extracted with DCM (3 x 3 mL). The organic fractions were combined, dried over anhydrous $Na_2SO_4$ and the solvent was removed under reduced pressure. The crude product was purified by flash chromatography (0-5% v/v MeOH in EtOAc) to yield compound **5e** as a pale purple solid (72 mg, 0.21 mmol, 64%). $R_f$ 0.28 (5% v/v MeOH in EtOAc); $^1$H-NMR (400 MHz, MeOD) $\delta$ 8.39 (d, $J$ = 6.1 Hz, 2H), 7.74 (s, 1H) 7.72 (dt, $J$ = 6.2, 2.1 Hz, 1H), 7.53 (s, 1H), 7.39 (m, 2H), 7.34 (d, $J$ = 8.4 Hz, 1H), 7.27 (d, $J$ = 5.7 Hz, 2H), 7.20 (d, $J$ = 3.1 Hz, 1H), 7.12 (dd, $J$ = 8.4, 1.6 Hz, 1H), 6.40 (dd, $J$ = 3.1, 0.7 Hz, 1H), 4.63 (s, 2H), 4.06 (s, 2H) ppm; $^{13}$C-NMR (100 MHz, MeOD) $\delta$ 169.8, 152.8, 150.0, 141.0, 137.0, 136.4, 133.3, 130.4, 130.0, 129.5, 129.1, 126.7, 126.0, 125.9, 122.4, 120.4, 112.2, 102.3, 45.3, 41.7 ppm; IR (solid) $\nu_{max}$ 3416-3250, 3031, 2920, 1639, 1600, 1581, 1526, 1480, 1416, 1322, 1285, 1216, 1099, 1067, 999, 891, 888, 809, 723, 693 cm$^{-1}$; LCMS (+ESI) *m/z* 342.3 [M+H]$^+$, retention time 1.40 min, (100%); HRMS (+ESI) *m/z* (Calcd. $C_{22}H_{20}N_3O$ [M+H]$^+$, 342.1606), *Obs.* 342.1602 ($\delta$ 1.2 ppm).

**N-((1H-Indol-5-yl)methyl)-4-(pyridin-4-ylmethyl)benzamide, (5f)**

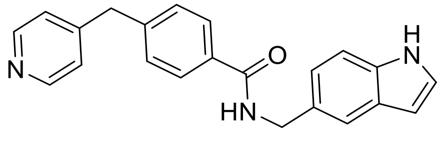


4-Pyridinylboronic acid (34 mg, 0.27 mmol), benzyl chloride **7b** (68 mg, 0.23 mmol), and $Na_2CO_3$ (51 mg, 0.48 mmol) were combined in a microwave vessel and flushed with $N_2$(g) for 5 minutes. $Pd(PPh_3)_4$ (5 mg, 0.005

mmol) was added, and the reaction vessel was flushed with $N_2$(g) for an additional 2-3 minutes before the addition of a mixture of DME and water (2:1 v/v, 1.5 mL). The reaction was heated at 100 °C for 4 h, then allowed to cool to room temperature and diluted with water (5 mL) and DCM (15 mL). The phases were separated, and the aqueous fraction was extracted with DCM (3 x 3 mL). The organic fractions were combined, dried over anhydrous $Na_2SO_4$ and the solvent was removed under reduced pressure. The crude material was purified twice by flash chromatography (0-5% v/v MeOH in DCM) to yield compound **5f** as a white solid (26 mg, 0.08 mmol, 33%). $R_f$ 0.08 (5% v/v MeOH in DCM);$^1$H-NMR (500 MHz, MeOD) $\delta$ 8.41 (d, $J$ = 6.2 Hz, 2H), 7.81 (d, $J$ = 8.4 Hz, 2H), 7.53 (d, $J$ = 0.8 Hz, 1H), 7.33 (m, $J$ = 8.6 Hz, 3H), 7.28 (d, $J$ = 6.1 Hz, 2H), 7.20 (d, $J$ = 3.1 Hz, 1H), 7.12 (dd, $J$ = 8.4, 1.6 Hz, 1H), 6.40 (dd, $J$ = 3.1, 0.9 Hz, 1H), 4.64 (s, 2H), 4.08 (s, 2H) ppm; $^{13}$C-NMR (125 MHz, MeOD) $\delta$ 169.7, 152.7, 150.1, 144.3, 137.0, 134.3, 130.4, 130.3, 129.5, 128.9, 126.0, 125.9, 122.4, 120.4, 112.2, 102.3, 45.2, 41.7 ppm; IR (solid) $v_{max}$ 3319, 3126, 3029, 2926, 2845, 1651, 1630, 1600, 1546, 1504, 1480, 1417, 1332, 1298, 1240, 1218, 1194, 1098, 1001, 986, 893, 877, 779, 757, 723, 663 cm$^{-1}$; LCMS (+ESI) $m/z$ 342.3 $[M+H]^+$, retention time 1.55 min, (100%); HRMS (+ESI) $m/z$ (Calcd. $C_{22}H_{20}N_3O$ $[M+H]^+$, 342.1601), Obs. 342.1598 ($\delta$ 0.8 ppm).

**(R)-N-(1-Phenylethyl)-3-(pyridin-4-ylmethyl)benzamide, (5h)**

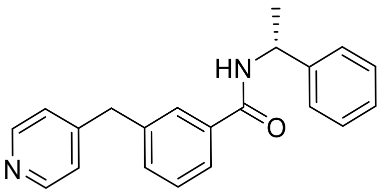


4-Pyridinylboronic acid (56 mg, 0.46 mmol), benzyl chloride **7c** (105 mg, 0.38 mmol), and $Na_2CO_3$ (85 mg, 0.80 mmol) were combined in a microwave vessel and flushed with $N_2$(g) for 5 minutes before the addition of $Pd(PPh_3)_4$ (9 mg, 0.01 mmol). The reaction vessel was flushed with $N_2$(g) for a further 2-3 minutes, and then a mixture of DME and water (2:1 v/v, 2 mL) was added, and the reaction was heated at 100 °C for 4 h. The reaction was then allowed to cool to room temperature, and diluted with water (5 mL) and DCM (5 mL). The phases were separated, and the aqueous fraction was extracted with DCM (3 x 3 mL). The organic fractions were combined, dried over anhydrous $Na_2SO_4$ and the solvent was removed under reduced pressure. The crude material was purified twice by flash chromatography (0-5% v/v MeOH in EtOAc, then 30-100% v/v EtOAc in DCM)). The product containing fractions were combined after each successive purification to yield compound **5h** as a colourless amorphous solid (73 mg, 0.23 mmol, 61%). $R_f$ 0.34 (5% v/v MeOH in EtOAc); $^1$H-NMR (500 MHz, MeOD) $\delta$ 8.41 (d, $J$ = 5.7 Hz, 2H), 7.73–7.71 (m, 2H), 7.43–7.38 (m, 4H), 7.32 (app. t, $J$ = 7.7 Hz, 2H), 7.29 (d, $J$ = 5.1 Hz, 2H), 7.22 (t, $J$ = 7.3 Hz, 1H), 5.23 (q, $J$ = 7.1 Hz, 1H), 4.08 (s, 2H), 1.55 (d, $J$ = 7.1 Hz, 3H) ppm; $^{13}$C-NMR (125 MHz, MeOD) $\delta$ 169.4, 152.8, 150.1, 145.3, 140.9, 136.4, 133.3, 130.0, 129.5, 129.1, 128.0, 127.2, 126.8, 125.9, 50.7, 41.7, 22.2 ppm; IR (solid) $v_{max}$ 3219, 3056, 3023, 2968, 2924, 2365, 1622, 1598, 1583, 1538, 1493, 1430, 1417, 1326, 1272, 1217, 1205, 1128, 1088, 1020, 999, 912, 791, 760, 698 cm$^1$; LCMS (+ESI)

*m/z* 317.1 $[M+H]^+$, 1.57 min, (100%); HRMS (+ESI) *m/z* (Calcd. $C_{21}H_{21}N_2O$ $[M+H]^+$, 317.1654), *Obs.* 317.1641 ($\delta$ 4.1 ppm).

**(R)-N-(1-Phenylethyl)-4-(pyridin-4-ylmethyl)benzamide, (5i)**

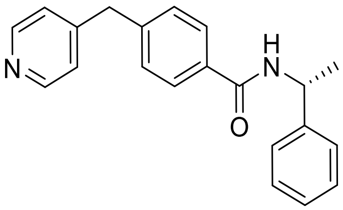

4-Pyridinylboronic acid (53 mg, 0.43 mmol), benzyl chloride **7d** (99 mg, 0.36 mmol), and $Na_2CO_3$ (80 mg, 0.76 mmol) were combined in a microwave vessel and flushed with $N_2$(g) for 5 minutes before the addition of $Pd(PPh_3)_4$ (8 mg, 0.01 mmol). The reaction vessel was flushed with $N_2$(g) for a further 2-3 minutes, and then a mixture of DME and water (2:1 v/v, 2 mL) was added, and the reaction was heated at 100 °C for 4 h. The reaction was then allowed to cool to room temperature, and diluted with water (5 mL) and DCM (5 mL). The phases were separated, and the aqueous fraction was extracted with DCM (3 x 3 mL). The organic fractions were combined, dried over anhydrous $Na_2SO_4$ and the solvent was removed under reduced pressure. The crude material was purified twice by flash chromatography (0-5% v/v MeOH in EtOAc, and 30-100% v/v EtOAc in DCM). The product containing fractions were combined after each successive purification to yield compound **5i** as a colourless amorphous solid (78 mg, 0.25 mmol, 68%). $R_f$ 0.29 (5% v/v MeOH in EtOAc); $^1$H-NMR (500 MHz, MeOD) $\delta$ 8.41 (d, $J$ = 5.8 Hz, 2H), 7.80 (d, $J$ = 8.2 Hz, 2H), 7.38 (d, $J$ = 7.1 Hz, 2H), 7.34–7.27 (m, 6H), 7.22 (t, $J$ = 7.3 Hz, 1H), 5.23 (q, $J$ = 7.1 Hz, 1H), 4.08 (s, 2H), 1.55 (d, $J$ = 7.1 Hz, 3H) ppm; $^{13}$C-NMR (125 MHz, MeOD) $\delta$ 169.3, 152.7, 150.1, 145.4, 144.3, 134.3, 130.2, 129.5, 128.9, 128.0, 127.2, 125.9, 50.7, 41.7, 22.3 ppm; IR (solid) $v_{max}$ 3411, 3180, 3034, 2974, 2931, 2164, 1631, 1601, 1542, 1505, 1493, 1446, 1416, 1348, 1310, 1297, 1277, 1206, 1189, 1119, 1065, 1015, 1006, 876, 831, 754, 743, 695 cm$^{-1}$; LCMS (+ESI) *m/z* 316.9 $[M+H]^+$, retention time 1.55 min, (100%); HRMS (+ESI) *m/z* (Calcd. $C_{21}H_{21}N_2O$ $[M+H]^+$, 317.1648), *Obs.* 317.1649 ($\delta$ 0.1 ppm).

**N-((1H-Indol-5-yl)methyl)-3-(pyridin-4-ylamino)benzamide, (5g)**

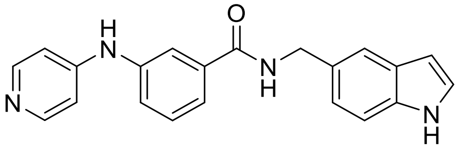

EDC.HCl (131 mg, 0.69 mmol) and HOAt (106 mg, 0.78 mmol) were added to a stirred solution of **4e** (122 mg, 0.57 mmol) in dry DCM (10 mL). 5-(Aminomethyl)indole (83 mg, 0.57 mmol) and DIPEA (199 µL, 1.14 mmol) were added to the solution, followed by anhydrous DMF (1 mL) to aid solubility. The reaction was allowed to stir for 48-72 hours at room temperature, then diluted with EtOAc (100 mL) and washed with water (20 mL). The aqueous fraction was extracted with EtOAc (10 mL) and the combined organic fractions were washed with saturated $NaHCO_3$ (10 mL) and brine (5 mL), before being dried over anhydrous $Na_2SO_4$ and concentrated under reduced pressure. The crude product was purified by flash chromatography (0-10% v/v MeOH in EtOAc) to yield compound **5g** as a pink amorphous solid (120 mg, 0.35 mmol, 61%). $R_f$ 0.03 (10% v/v MeOH in EtOAc); $^1$H-NMR (500 MHz, $d_6$-DMSO) $\delta$ 11.01 (s, 1H), 8.99 (app. t, $J$ = 6.0 Hz, 1H), 8.92 (s, 1H), 8.21 (d, $J$ = 6.4 Hz, 1H), 7.72 (app. t, $J$ = 1.9 Hz, 1H), 7.55 (dt, $J$ = 7.7, 1.3 Hz, 1H), 7.48 (s, 1H), 7.42 (app. t, $J$ = 7.8 Hz, 1H), 7.34–7.33 (m, 2H), 7.31 (app. t, $J$ = 2.7 Hz, 1H), 7.08 (dd, $J$ = 8.4, 1.6 Hz, 1H), 6.92 (d, $J$ = 6.4 Hz, 1H), 6.38 (tt, $J$ = 2.0, 0.9 Hz, 1H), 4.54 (d, $J$ = 5.9 Hz, 2H) ppm; $^{13}$C-NMR (125 MHz, $d_6$-DMSO) $\delta$ 165.7, 150.2, 149.8, 140.7, 135.9, 134.9, 129.9, 129.3, 127.5, 125.5, 122.3, 121.0, 121.0, 118.7, 118.7, 111.2, 109.4, 100.9, 43.2 ppm; IR (solid) $v_{max}$ 3300-2900, 3265, 3022, 2912, 1724, 1635, 1575, 1516, 1482, 1432, 1341, 1216, 1147, 1094, 1044, 995, 984, 814, 754, 727, 697 cm$^{-1}$; LCMS (+ESI) *m/z* 343.1 $[M+H]^+$, retention time 1.54 min, (100%); HRMS (+ESI) *m/z* (Calcd. $C_{21}H_{19}N_4O$ $[M+H]^+$, 343.1559), *Obs.* 343.1565 ($\delta$ 2.3 ppm).

**N-(4-(Pyridin-4-ylamino)phenyl)benzamide, (5b)**

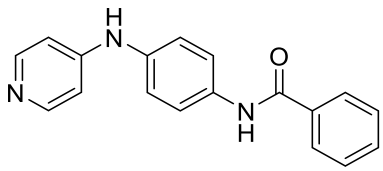

Aniline **4b** (49 mg, 0.26 mmol) and DIPEA (45 uL, 0.26 mmol) were added to a stirred solution of benzoic acid (32 mg, 0.26 mmol) and HATU (99 mg, 0.26 mmol) in anhydrous DCM (3 mL) at 0 $^oC$. The reaction was allowed to come slowly to room temperature and stirred for 24 hours. When complete, the reaction was diluted with EtOAc (20 mL) and water (10 mL), made slightly basic with sat. $NaHCO_3$. The phases were separated and the aqueous phase was extracted with EtOAc (3 x 10 mL). The organic fractions were combined, washed with brine (5 mL) and the solvent was removed under reduced pressure. The crude product was purified by flash chromatography (10-100% v/v EtOAc in DCM, then 5-15% v/v MeOH in EtOAc) to yield compound **5b** as an off-white solid (50 mg, 0.17 mmol, 66%). $R_f$ 0.33 (10% v/v MeOH in DCM); $^1$H-NMR (500 MHz, $d_6$-DMSO) $\delta$ 10.23 (s, 1H), 8.72 (d, $J$ = 1.8 Hz, 1H), 8.16 (d, $J$ = 5.6 Hz, 2H), 7.95 (d, $J$ = 7.1 Hz, 2H), 7.76 (d, $J$ = 8.5 Hz, 2H), 7.59 (t, $J$ = 7.3 Hz, 1H), 7.53 (app. t, $J$ = 7.5 Hz, 2H), 7.19 (d, $J$ = 8.8 Hz, 2H), 6.85 (d, $J$ = 6.1 Hz, 2H) ppm; $^{13}$C-NMR (125 MHz, $d_6$-DMSO) $\delta$ 165.3, 150.5, 150.0, 136.0, 135.0, 134.3, 131.5, 128.4, 127.6, 121.5, 120.9, 108.8 ppm; IR (solid) $v_{max}$ 3381, 3248, 3033, 2938, 1646, 1619, 1592, 1537, 1511, 1430, 1405, 1349, 1315, 1258, 1215, 1105, 995, 893, 814, 794, 714, 702, 688, 649 cm$^{-1}$; LCMS (+ESI) *m/z* 290.2 $[M+H]^+$, retention time 1.55 min, (100%); HRMS (+ESI) *m/z* (Calcd. $C_{18}H_{16}N_3O$ $[M+H]^+$, 290.1288), *Obs.* 290.1275 ($\delta$ 4.3 ppm).

**N-(4-(Methyl(pyridin-4-yl)amino)phenyl)benzamide, (5c)**

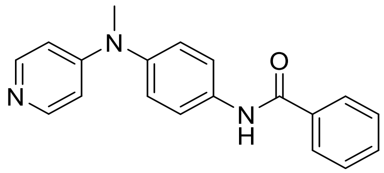

Aniline **8b** (60 mg, 0.30 mmol) and DIPEA (52 uL, 0.3 mmol) were added to a stirred solution of benzoic acid (37 mg, 0.3 mmol) and HATU (114 mg, 0.3 mmol) in anhydrous DCM (2.5 mL) at 0 °C. The reaction was allowed to come slowly to room temperature and stirred for 24 hours. When complete, the reaction was diluted with EtOAc (20 mL) and water (10 mL), made slightly basic with sat. $NaHCO_3$. The phases were separated, and the aqueous phase was extracted with EtOAc (2 x 5 mL). The organic fractions were combined, washed with brine (5 mL), and the solvent was removed under reduced pressure. The crude product was purified by flash chromatography (10-50% v/v EtOAc in PE, then 10% v/v MeOH in EtOAc) to yield compound **5c** as an amorphous brown solid (28 mg, 0.09 mmol, 31%). $R_f$ 0.21 (10% v/v MeOH in DCM); $^1$H-NMR (400 MHz, $CDCl_3$) $\delta$ 8.19 (d, $J$ = 6.1 Hz, 2H), 8.04 (s, br, 1H), 7.90 (d, $J$ = 7.0 Hz, 2H), 7.73 (d, $J$ = 8.8 Hz, 2H), 7.58 (tt $J$ = 7.3, 1.3 Hz, 1H), 7.51 (t, $J$ = 7.3 Hz, 2H), 7.23 (d, $J$ = 8.8 Hz, 2H), 6.56 (d, $J$ = 6.6 Hz, 2H), 3.33 (s, 3H); $^{13}$C-NMR (100 MHz, $CDCl_3$) $\delta$ 166.0, 154.3, 149.1, 142.2, 136.5, 134.8, 132.2, 129.0, 127.6, 127.2, 121.9, 108.3, 39.7 ppm; IR (solid) $\nu_{max}$ 3219, 3054, 1645, 1592, 1529, 1502, 1406, 1363, 1313, 1243, 1223, 1137, 1102, 1072, 991, 880, 841, 807, 704, 669 cm$^{-1}$; LCMS (+ESI) *m/z* 304.0, retention time 1.49 min, (100%); HRMS (+ESI) *m/z* (Calcd. $C_{19}H_{18}N_3O$ [M+H]$^+$, 304.1444), *Obs.* 304.1439 ($\delta$ 2.0 ppm).

**N-(4-(Pyridin-4-ylmethyl)phenyl)benzamide, (5a)**

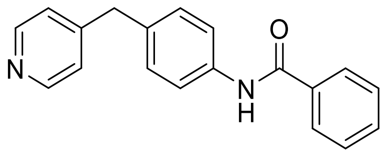

4-(4-Aminophenyl)pyridine **3d** (110 mg, 0.60 mmol) and DIPEA (105 uL, 1.2 mmol) were added to a stirred solution of benzoic acid (73 mg, 0.60 mmol) and HATU (228 mg, 0.60 mmol) in anhydrous DCM (3 mL) at 0 °C. The reaction was allowed to come slowly to room temperature and stirred for 24 hours. When complete, the reaction was diluted with EtOAc (20 mL), water (10 mL), and made slightly basic with sat. $NaHCO_3$. The phases were separated, and the aqueous phase was extracted with EtOAc (3 x 10 mL), the organic fractions were combined, washed with brine (5 mL), and the solvent was removed under reduced pressure. The crude product was purified by flash chromatography (10-100% v/v EtOAc in PE, then 5% v/v MeOH in EtOAc) to yield compound **5a** as a white solid (133 mg, 0.46 mmol, 77%). $R_f$ 0.09 (2:1 v/v EtOAc:PE); $^1$H-NMR (400 MHz, $d_6$-DMSO) $\delta$ 10.22 (s, 1H), 8.46 (d, $J$ = 6.1 Hz, 2H), 7.94 (m, 2H), 7.71 (d, $J$ = 8.5 Hz, 2H), 7.58 (tt, $J$ = 7.4, 1.4 Hz, 1H), 7.52 (dd, $J$ = 7.7, 7.0 Hz, 2H), 7.25-7.22 (m, 4H), 3.94 (s, 2H) ppm; $^{13}$C-NMR (100 MHz, $d_6$-DMSO) $\delta$ 165.5, 150.3, 149.6, 137.6, 134.8, 131.5, 129.1, 128.4, 127.6, 124.0, 120.6, 38.3 ppm; IR (solid) $\nu_{max}$ 3362,

3044, 2916, 1655, 1597, 1579, 1525, 1491, 1411, 1323, 1311, 1259, 1227, 1184, 1104, 1072, 1026, 1006, 858, 829, 815, 793, 775, 741, 712, 690, 672 cm$^{-1}$; LCMS (+ESI) *m/z* 289.2 [M+H]$^+$, 1.90 min, (100%); HRMS (+ESI) *m/z* (Calcd. $C_{19}H_{17}N_2O$ [M+H]$^+$, 289.1335), *Obs.* 289.1322 ($\delta$ 4.7 ppm).

**(S)-2-(Benzylamino)-3-(1H-indol-3-yl)-N-(4-(pyridin-4-ylmethyl)phenyl)propanamide, (5d)**

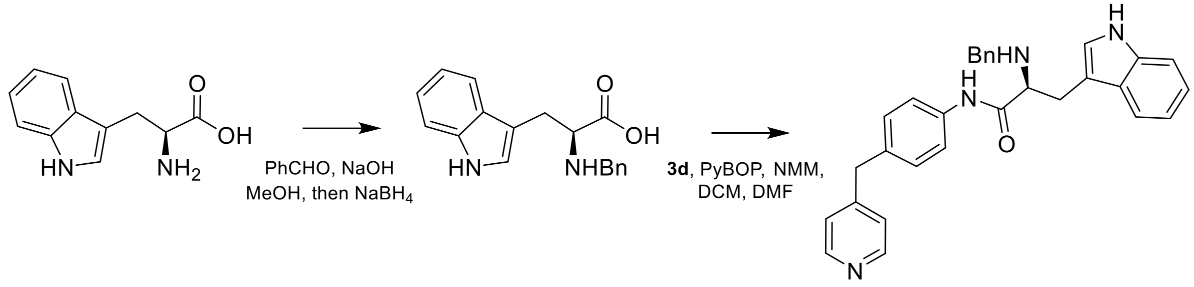


**Step 1:** Benzaldehyde (203 µL, 2.0 mmol) was added to a stirred suspension of L-tryptophan (408 mg, 2.0 mmol) and NaOH (84 mg, 2.1 mmol) in dry MeOH (5 mL), and allowed to stir at room temperature for 1 hour. The reaction was then cooled to 0 °C and sodium borohydride (99 mg, 2.6 mmol) was added as a single portion. The reaction was allowed to come slowly to room temperature and stirred for 2 hours and then concentrated under reduced pressure. The residue was diluted with water (5 mL) and brought to pH ~ 5 using 1.5 M HCl solution. The resulting precipitate was collected under reduced pressure, washed with iced water (10 mL), ice cold MeOH (5 mL), and then dried under reduced pressure to yield benzyl-L-tryptophan as a white solid (486 mg, 1.65 mmol, 83%). $^1$H-NMR (500 MHz, $d_6$-DMSO) $\delta$ 10.9 (s, 1H), 7.49 (d, *J* = 7.9 Hz, 1H), 7.33 (d, *J* = 8.1 Hz, 1H), 7.29–7.24 (m, 5H), 7.15 (d, *J* = 2.3 Hz, 1H), 7.05 (ddd, *J* = 8.1, 6.9, 1.2 Hz, 1H), 6.95 (ddd, *J* = 7.9, 7.4, 0.9 Hz, 1H), 3.81 (d, *J* = 13.4 Hz, 2H), 3.69 (d, *J* = 13.4 Hz, 2H), 3.41 (t, *J* = 6.5 Hz, 1H), 3.12 (dd, *J* = 14.6, 6.2 Hz, 1H), 3.02 (dd, *J* = 14.6, 6.7 Hz, 1H) ppm; $^{13}$C-NMR (125 MHz, $d_6$-DMSO) $\delta$ 173.6, 138.0, 136.1, 128.4, 128.2, 127.4, 127.2, 123.7, 120.8, 118.4, 118.21, 111.3, 110.1, 61.2, 50.5, 27.9 ppm; IR (solid) $\nu_{max}$ 3049, 2968, 2880, 2702-2452, 1596, 1551, 1529, 1517, 1497, 1431, 1417, 1348, 1232, 1222, 1147, 1090, 1065, 1022, 933, 805, 751, 691, 654 cm$^{-1}$; LCMS (+ESI) *m/z* 295.2 [M+H]$^+$, 1.46 min, (100%); HRMS (+ESI) *m/z* (Calcd. $C_{18}H_{18}N_2O_2Na$ [M+Na]$^+$, 317.1260), *Obs.* 317.1267

**Step 2:** Benzyl-L-tryptophan (147 mg, 0.50 mmol), *n*-methyl morpholine (121 µL, 1.10 mmol) and then PyBOP (260 mg, 0.50 mmol) were added in quick succession to a stirred solution of 4-(pyridin-4-ylmethyl)aniline (92 mg, 0.5 mmol) in anhydrous DCM (2 mL). The reaction was allowed to stir at room temperature for 1 hour and then anhydrous DMF (300 µL) was added to aid solubility. The reaction was stirred at room temperature for a further 4 hours and then concentrated under reduced pressure. The residue was diluted with EtOAc (30 mL), washed with water (5 mL) and saturated $NaHCO_3$ (5 mL), dried over anhydrous $Na_2SO_4$ and then the solvent was removed under reduced pressure. The crude product was purified by flash chromatography (0-5% v/v MeOH in DCM) and the product containing fractions were concentrated under reduced pressure. The resulting oil was redissolved in EtOAc (50 mL), washed with water (4 x 10 mL), and brine (10 mL), dried over anhydrous $Na_2SO_4$ and the solvent was removed under reduced pressure to yield compound **5d** as a yellow amorphous solid (168 mg, 0.37 mmol, 73%). $R_f$ 0.08 (2:1 v/v EtOAc:PE); $^1$H-NMR (500 MHz, $CDCl_3$) $\delta$ 9.41 (s, 1H, NH), 8.50 (d, *J* = 5.0 Hz, 2H), 8.47 (d, *J* = 4.6 Hz, 1H, NH), 8.11 (br s, 1H, NH), 7.66 (d, *J* = 7.8 Hz, 1H), 7.53 (d, *J* = 8.5 Hz, 1H), 7.38 (d, *J* = 8.2 Hz, 1H), 7.26-7.19 (m, 3H), 7.14 (d, *J* = 8.4 Hz, 2H), 7.12–7.09 (m, 3H), 7.06 (dd, *J* = 7.4, 2.0 Hz, 1H), 7.01 (d, *J* = 2.4 Hz, 1H), 6.96 (d, *J* = 8.4 Hz, 1H), 6.65 (d, *J* = 8.4 Hz, 1H), 3.95 (s, 2H), 3.76 (d, *J* = 13.4 Hz, 1H), 3.63 (d, *J* = 13.8 Hz, 1H), 3.61 (m, 1H), 3.44 (ddd, *J* = 14.7, 4.1, 1.0 Hz, 1H), 3.04 (dd, *J* = 14.8, 9.4 Hz, 2H) ppm; $^{13}$C-NMR (125 MHz, $CDCl_3$) $\delta$ 172.4, 150.3, 149.9, 145.1, 139.1, 136.6, 136.5, 134.6, 130.1, 129.7, 128.7, 128.0, 127.6, 127.4, 124.3, 123.0, 122.6, 120.0, 119.9, 119.0, 115.5, 111.4, 63.1, 53.0, 40.8, 29.1 ppm; IR (solid) $v_{max}$ 3431, 3321, 3193, 3028, 2921, 2853, 1665, 1632, 1601, 1515, 1496, 1454, 1412, 1342, 1295, 1234, 1180, 1108, 1067, 999, 919, 846, 811, 738, 697 cm$^{-1}$; LCMS (+ESI) *m/z* 461.4 [M+H]$^+$, 1.45 min, (100%); HRMS (+ESI) *m/z* (Calcd. $C_{30}H_{29}N_4O$ [M+H]$^+$, 461.2336), *Obs*. 461.2269 ($\delta$ 1.3 ppm).

**N-(4-(Pyridin-4-ylmethyl)phenyl)benzenesulfonamide, (5j)**

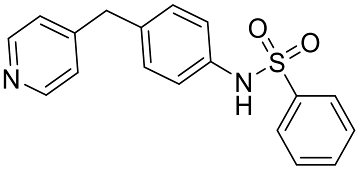


Benzene sulfonyl chloride (77 µL, 0.6 mmol) was added to a solution of 4-(4-aminobenzyl)pyridine **3d** (110 mg, 0.6 mmol) in anhydrous pyridine (2 mL) and the reaction was stirred overnight at room temperature. When complete, the reaction was diluted with DCM (25 mL) and water (10 mL), the phases were separated, and the aqueous phase was extracted with DCM (3 mL). The organic fractions were combined, dried over anhydrous $Na_2SO_4$ and the solvent was removed under reduced pressure. The crude product was purified by flash chromatography (0-100% v/v EtOAC in PE, then 0-10% v/v MeOH in EtOAc) to yield compound **5j** as a white solid (95 mg, 0.29 mmol, 49%). $R_f$ 0.09 (2:1 v/v EtOAc:PE); $^1$H-NMR (400 MHz, $d_6$-DMSO) $\delta$ 10.22 (s, 1H), 8.42 (d, *J* = 6.0 Hz, 2H), 7.73 (m, 2H), 7.59 (tt, *J* = 7.5, 1.2 Hz, 1H), 7.53 (app. t, *J* = 7.5 Hz, 2H), 7.15 (d, *J* = 5.9 Hz, 2H), 7.09 (d, *J* = 8.4 Hz, 2H), 7.01 (d, *J* = 8.5 Hz, 2H), 3.84 (s, 2H) ppm; $^{13}$C-NMR (100 MHz, $d_6$-DMSO) $\delta$ 149.9, 149.6, 139.5, 136.0, 135.3, 132.9, 129.6, 129.2, 126.6, 124.0, 120.5, 39.4 ppm; IR (solid) $v_{max}$ 3063, 3020, 2829, 2654, 1677, 1604, 1558, 1508, 1445, 1425, 1326, 1304, 1290, 1230, 1219, 1161, 1092, 1068, 1007, 964, 922, 849, 804, 787, 755, 723, 714, 700, 689 cm$^{-1}$; LCMS (+ESI) *m/z* 325.2 [M+H]$^+$, retention time 1.40 min, (96%); HRMS (+ESI) *m/z* (Calcd. $C_{18}H_{17}N_2O_2S$ [M+H]$^+$, 325.1005), *Obs*. 325.0994 ($\delta$ 3.4 ppm).

**N-(4-(Pyridin-4-ylmethyl)phenyl)-4-(trifluoromethoxy)benzenesulfonamide, (5k)**

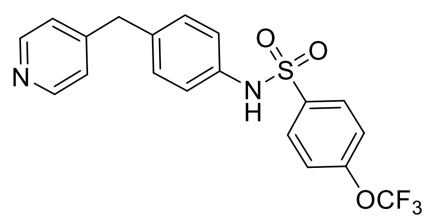

4-(Trifluoromethoxy)benzene sulfonyl chloride (51 µL, 0.44 mmol) was added to a stirred solution of 4-(pyridin-4-ylmethyl)aniline (74 mg, 0.40 mmol) and $Et_3N$ (112 µL, 0.80 mmol) in dry DCM (3 mL). The reaction was stirred at room temperature for 20 hours and then diluted with DCM (20 mL) and washed with water (5 mL). The aqueous phase was extracted with DCM (2 mL) and the combined organic fractions were washed with brine (2 mL). The solvent was removed under reduced pressure and the crude product was purified by flash chromatography (20-80% v/v EtOAc in PE) to yield compound **5k** as a pale pink solid (46 mg, 0.11 mmol, 28%). $R_f$ 0.10 (50% v/v EtOAc in PE); $^1$H-NMR (500 MHz, $CDCl_3$) $\delta$ 8.49 (d, $J$ = 6.0 Hz, 2H), 7.80 (d, $J$ = 9.0 Hz, 2H), 7.25 (d, $J$ = 7.5 Hz, 2H), 7.14 (s, 1H), 7.08–7.02 (m, 6H), 3.91 (s, 2H) ppm; $^{13}$C-NMR (125 MHz, $CDCl_3$) $\delta$ 152.5, 150.0, 149.7, 137.6, 136.7, 134.8, 130.2, 129.5, 124.3, 122.6, 120.9, 119.3, 40.7 ppm; IR (solid) $v_{max}$ 3009, 2924, 2645, 1606, 1563, 1510, 1489, 1421, 1332, 1258, 1212, 1154, 1095, 1008, 826, 808, 765, 708, 686, 625 cm$^{-1}$; LCMS (+ESI) *m/z* 409.2 [M+H]$^+$, 1.80 min, (100%); HRMS (+ESI) *m/z* (Calcd. $C_{19}H_{16}N_2F_3S$ [M+H]$^+$, 409.0828), *Obs.* 409.0813 ($\delta$ 3.8 ppm).

**N-(4-Methoxybenzyl)-3-(pyridin-4-ylmethyl)aniline, (5l)**

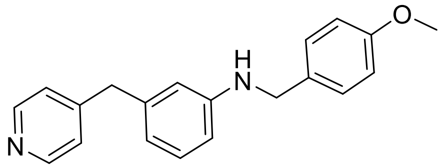

Glacial AcOH (0.7 mL) was added to a solution of 3-(pyridin-4-ylmethyl)aniline **3c** (73 mg, 0.39 mmol) and *p*-anisaldehyde (88 uL, 0.75 mmol) in dry MeOH (5 mL). The reaction was stirred at room temperature for 30 min, and then $NaCNBH_3$ (25 mg, 0.39 mmol) was added as a single portion. The reaction was stirred at room temperature for 24 h and then concentrated under reduced pressure. EtOAc (40 mL) and water (10 mL) were added, and the mixture was brought to pH 8 using saturated $NaHCO_3$ solution. The phases were separated, and the aqueous fraction was extracted with EtOAc (2 x 10 mL). The combined organic fractions were washed with brine (5 mL), dried over anhydrous $Na_2SO_4$ and the solvent was removed under reduced pressure. The crude product was purified by flash chromatography (0-5% v/v MeOH in EtOAc) to yield compound **5l** as a yellow solid (51 mg, 0.17 mmol, 43%). $R_f$ 0.22 (50% v/v EtOAc:PE); $^1$H-NMR (500 MHz, $CDCl_3$) $\delta$ 8.47 (d, $J$ = 5.1 Hz, 2H), 7.26 (m, 2H), 7.11 (m, 3H), 6.87 (dd, $J$ = 8.2, 1.5Hz, 2H), 6.52 (dd, $J$ = 8.3, 3.1 Hz, 2H), 6.41 (m, 1H), 4.22 (s, 2H), 3.86 (s, 2H) 3.78 (s, 3H) ppm; $^{13}$C-NMR (125 MHz, $CDCl_3$) $\delta$ 158.9, 150.5, 149.4, 148.5, 139.9, 131.1, 129.6, 128.8, 124.3, 118.2, 114.0, 113.4, 111.2, 55.3, 47.7, 41.4 ppm; IR (solid) $v_{max}$ 3272, 3033, 2995,

2837, 1599, 1584, 1558, 1530, 1510, 1488, 1466, 1416, 1331, 1298, 1249, 1172, 1159, 1103, 1032, 997, 927, 854, 830, 813, 762, 748, 726, 690, 625, 605 $cm^{-1}$; LCMS (+ESI) *m/z* 305.3, retention time 2.48 min, (100%); HRMS (+ESI) *m/z* (Calcd. $C_{20}H_{21}N_2$ $O_1$ $[M+H]^+$, 305.1648), *Obs.* 305.1642 ($\delta$ 2.2 ppm).

**N-(4-Methoxybenzyl)-4-(pyridin-4-ylmethyl)aniline, (5m)**

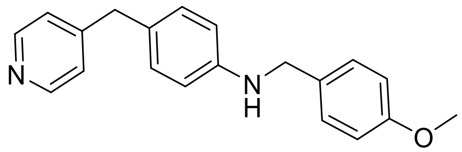

Glacial AcOH (1 mL) was added to a solution of 4-(pyridin-4-ylmethyl)aniline **3d** (138 mg, 0.75 mmol) and *p*-anisaldehyde (91 uL, 0.75 mmol) in dry MeOH (7.5 mL). The reaction was stirred at room temperature for 30 min, and then $NaCNBH_3$ (47 mg, 0.75 mmol) was added as a single portion. The reaction was stirred at room temperature for 17 h and then concentrated under reduced pressure. DCM (10 mL) and water (10 mL) were added, and the mixture was brought to pH 8 using saturated $NaHCO_3$ solution. The phases were separated, and the aqueous fraction was extracted with DCM (2 x 5 mL). The combined organic fractions were washed with brine (3 mL), dried over anhydrous $Na_2SO_4$ and the solvent was removed under reduced pressure. The crude product was purified by flash chromatography (25-100% EtOAc in DCM) to yield compound **5m** as a white solid (186 mg, 0.61 mmol, 82%). $R_f$ 0.20 (2:1 v/v EtOAc:PE); $^1$H-NMR (500 MHz, $CDCl_3$) $\delta$ 8.47 (d, $J$ = 6.1 Hz, 2H), 7.28 (d, $J$ = 8.7 Hz, 2H), 7.09 (d, $J$ = 6.1 Hz, 2H), 6.97 (d, $J$ = 8.6 Hz, 2H), 6.88 (d, $J$ = 8.6 Hz, 2H), 6.59 (d, $J$ = 8.5 Hz, 2H), 4.24 (s, 2H), 3.85 (s, 2H), 3.80 (s, 3H) ppm; $^{13}$C-NMR (125 MHz, $CDCl_3$) $\delta$ 159.0, 151.1, 149.8, 147.1, 131.4, 130.0, 128.9, 127.8, 124.2, 114.2, 113.2, 55.4, 48.0, 40.5 ppm; IR (solid) $v_{max}$ 3250, 3072, 3018, 2960, 2913, 2837, 1609, 1599, 1512, 1472, 1457, 1441, 1415, 1312, 1299, 1260, 1243, 1219, 1181, 1171, 1106, 1089, 1027, 997, 857, 817, 807, 777, 717 $cm^{-1}$; LCMS (+ESI) *m/z* 305.3 $[M+H]^+$, 1.72 min, (100%); HRMS (+ESI) *m/z* (Calcd. $C_{20}H_{21}N_2O$ $[M+H]^+$, 305.1648), *Obs.* 305.1644 ($\delta$ 1.4 ppm).

**N-(1-(4-Methoxyphenyl)ethyl)-4-(pyridin-4-ylmethyl)aniline, (5o)**

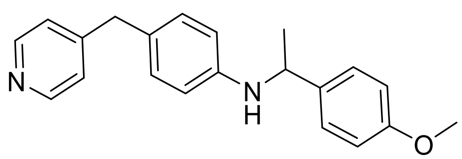

A 1 M solution of $TiCl_4$ in DCM (1.3 mL, 1.3 mmol) was added to a solution of 4'-methoxyacetophenone (151 g, 1.0 mmol) in dry DCM (6 mL). The mixture was cooled to 0 °C and 4-(pyridin-4-ylmethyl)aniline (372 mg, 2 mmol) was added. The reaction was then allowed to warm to room temperature and stir for 3 hours before a methanolic solution of $Na(CN)BH_3$ (185 µL of a 6.5 M solution, 1.2 mmol) was added. The reaction was stirred at room temperature for 24 hours and then quenched with 2 M NaOH (until pH 10). The mixture was filtered, and the filtrate was partitioned between EtOAc (50 mL) and water (25 mL). The organic layer separated and washed with water (2 x 20 mL) and brine (20 mL), dried over anhydrous $Na_2SO_4$ and the solvent was removed under reduced pressure. The crude product was purified by flash chromatography (20-80% v/v EtOAc in DCM, followed by 0-5% v/v MeOH in EtOAC) to obtain compound **5o** as a yellow amorphous solid (88 mg, 0.28 mmol, 28%). $R_f$ 0.26 (50% v/v EtOAc in Pet. ether); $^1$H-NMR (400 MHz, $CDCl_3$) $\delta$ 8.44 (d, $J$ = 5.9 Hz, 2H), 7.27 (d, $J$ = 8.8 Hz, 2H), 7.06 (d, $J$ = 5.9 Hz, 2H), 6.89 (d, $J$ = 8.4 Hz, 2H), 6.85 (d, $J$ = 8.7 Hz, 2H), 6.46 (d, $J$ = 8.5 Hz, 2H), 4.41 (q, $J$ = 6.7 Hz, 1H), 3.98 (br s, 1H), 3.80 (s, 2H), 3.78 (s, 3H), 1.48 (d, $J$ = 6.8 Hz, 3H) ppm; $^{13}$C-NMR (100 MHz, $CDCl_3$) $\delta$ 158.5, 151.0, 149.7, 146.1, 137.2, 129.7, 127.3, 126.9, 124.1, 114.0, 113.5, 55.3, 53.0, 40.4, 25.1 ppm; IR (solid) $v_{max}$ 3307, 3035, 2960, 2833, 1610, 1586, 1559, 1509, 1460, 1437, 1415, 1364, 1321, 1286, 123, 1222, 1182, 1167, 1097, 1028, 1012, 1006, 995, 941, 911, 879, 851, 826, 808, 767, 741, 666 cm$^{-1}$; LCMS (+ESI]) *m/z* 319.3, retention time 1.55 min, (100%); HRMS (+ESI) *m/z* (Calcd. $C_{21}H_{23}ON_2$ $[M+H]^+$, 319.1810), *Obs.* 319.1815 ($\delta$ 1.6 ppm).

**N-(3-(Methylsulfonyl)benzyl)-4-(pyridin-4-ylmethyl)aniline, (5p)**

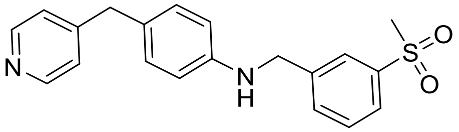


Glacial AcOH (1 mL) was added to a solution of 4-(pyridin-4-ylmethyl)aniline **3d** (138 mg, 0.75 mmol) and 3-methylsulfonylbenzaldehyde (131 mg, 0.75 mmol) in dry MeOH (7.5 mL). The reaction was stirred at room temperature for 30 min, and then $NaCNBH_3$ (47 mg, 0.75 mmol) was added as a single portion. The reaction was stirred at room temperature for 17 h and then concentrated under reduced pressure. DCM (10 mL) and water (10 mL) were added, and the mixture was brought to pH 8 using saturated $NaHCO_3$ solution. The phases were separated, and the aqueous fraction was extracted with DCM (2 x 5 mL). The combined organic fractions were washed with brine (3 mL), dried over anhydrous $Na_2SO_4$ and the solvent was removed under reduced pressure. The crude product was purified by flash chromatography (25-100% EtOAc in DCM) to yield compound **5p** as a yellow solid (209 mg, 0.59 mmol, 79%). $R_f$ 0.05 (2:1 v/v EtOAc:PE); $^1$H-NMR (500 MHz, $CDCl_3$) $\delta$ 8.46 (d, $J$ = 6.0 Hz, 1H), 7.94 (s, 1H), 7.84 (ddd, $J$ = 7.7, 2.0, 1.1 Hz, 1H), 7.66 (ddd, $J$ = 7.6, 1.9, 1.0 Hz, 1H), 7.53 (app. t, $J$ = 7.7 Hz, 1H), 7.08 (d, $J$ = 6.0 Hz, 1H), 6.97 (d, $J$ = 8.5 Hz, 2H), 6.55 (d, $J$ = 8.5 Hz, 2H), 4.42 (d, $J$ = 4.8 Hz, 2H), 4.21 (t, $J$ = 5.7 Hz, 1H), 3.84 (s, 2H), 3.03 (s, 3H) ppm; $^{13}$C-NMR (125 MHz, $CDCl_3$) $\delta$ 150.9, 149.8, 146.3, 141.8, 141.1, 132.6, 130.1, 129.8, 128.5, 126.2, 126.0, 124.2, 113.3, 47.9, 44.5, 40.5 ppm; (solid) $v_{max}$ 3251, 3027, 2916, 2869, 2845, 1611, 1603, 1560, 1519, 1477, 1416, 1316, 1291, 1261, 1220, 1181, 1142, 1094, 996, 960, 924, 865, 810, 778, 760, 686 cm$^{-1}$; LCMS (+ESI) *m/z* 353.2 $[M+H]^+$, 1.43 min, (100%); HRMS (+ESI) *m/z* (Calcd. $C_{20}H_{21}N_2O_2S$ $[M+H]^+$, 353.1318), *Obs.* 353.1313 ($\delta$ 1.4 ppm).

**N$^1$-(4-Methoxybenzyl)-N$^4$-methyl-N$^4$-(pyridin-4-yl)benzene-1,4-diamine, (5n)**

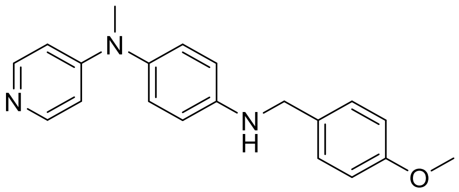

Glacial AcOH (0.7 mL) was added to a solution of aniline **8b** (62 mg, 0.31 mmol) and *p*-anisaldehyde (73 uL, 0.62 mmol) in dry MeOH (5 mL). The reaction was stirred at room temperature for 30 min, and then $NaCNBH_3$ (20 mg, 0.31 mmol) was added as a single portion. The reaction was stirred at room temperature for 24 h and then concentrated under reduced pressure. EtOAc (40 mL) and water (10 mL) were added, and the mixture was brought to pH 8 using saturated $NaHCO_3$ solution. The phases were separated, and the aqueous fraction was extracted with EtOAc (2 x 10 mL). The combined organic fractions were washed with brine (5 mL), dried over anhydrous $Na_2SO_4$ and the solvent was removed under reduced pressure. The crude product was purified by flash chromatography (0-10% v/v MeOH in DCM) to yield compound **5n** as an off-white solid (49 mg, 0.15 mmol, 50%). $R_f$ 0.32 (10% v/v MeOH in DCM); $^1$H-NMR (500 MHz, $CDCl_3$) $\delta$ 8.15 (d, $J$ = 6.0 Hz, 2H), 7.31 (d, $J$ = 8.7 Hz, 2H), 6.99 (d, $J$ = 8.7 Hz, 2H), 6.91 (d, $J$ = 8.7 Hz, 2H), 6.67 (d, $J$ = 8.7 Hz, 2H), 6.47 (d, $J$ = 6.6 Hz, 2H), 4.27 (s, 2H), 3.82 (s, 3H), 3.25 (s, 3H) ppm; $^{13}$C-NMR (125 MHz, $CDCl_3$) $\delta$ 159.1, 154.9, 148.9, 147.1, 135.9, 131.1, 128.9, 128.2, 114.2, 113.9, 107.8, 55.5, 48.0, 39.8 ppm; IR (solid) $\nu_{max}$ 3291, 2826, 1642, 1608, 1593, 1535, 1508, 1471, 1442, 1372, 1322, 1303, 1243, 1222, 1181, 1114, 1035, 983, 872, 833, 807, 798, 759 cm$^{-1}$; LCMS (+ESI) *m/z* 320.1, retention time 1.80 min (100%); HRMS (+ESI) *m/z* (Calcd. $C_{20}H_{22}N_3O$ [M+H]$^+$, 320.1757), Obs. 320.1751 ($\delta$ 2.0 ppm).

**NMR Spectra of lead compounds tested in anti-tubercular assays**

N-(4-(Pyridin-4-ylmethyl)phenyl)benzamide, (5a), 1H-NMR, 400 MHz, d6-DMSO

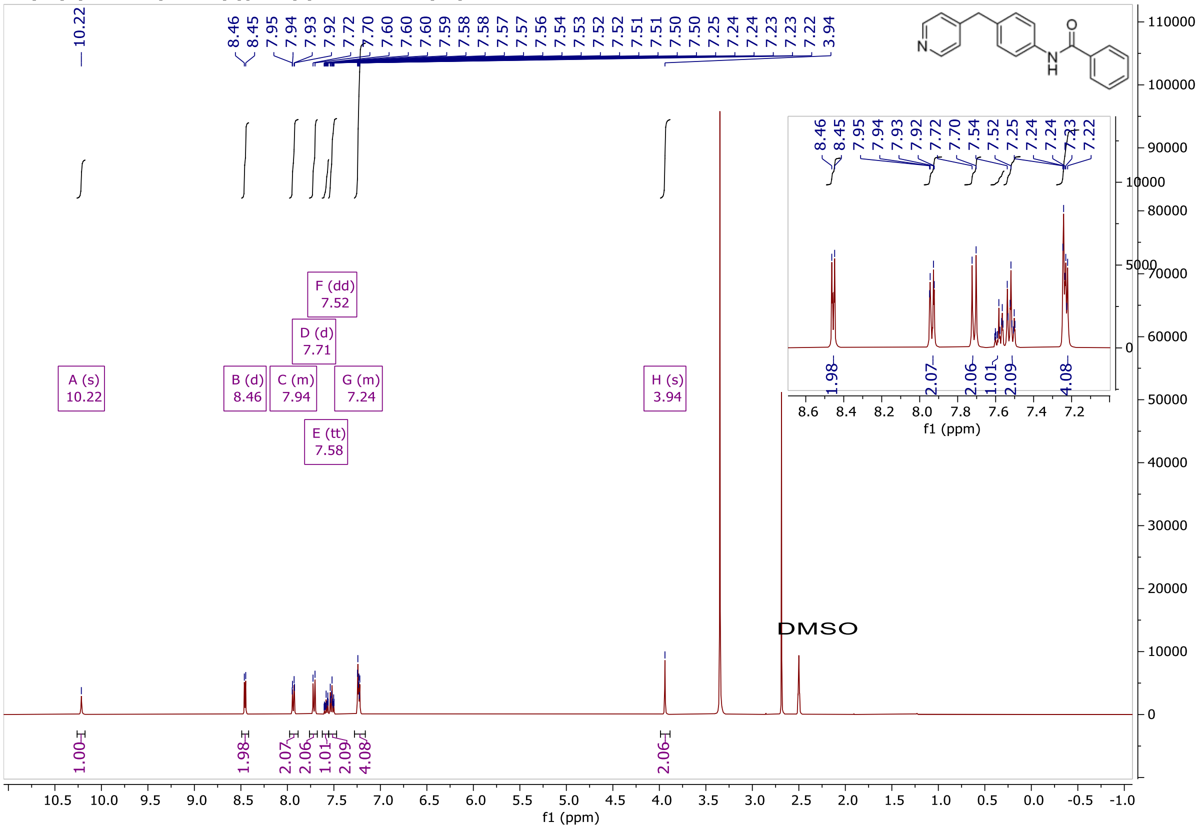


N-(4-(Pyridin-4-ylmethyl)phenyl)benzamide, (5a), 13C-NMR, 100 MHz, d6-DMSO

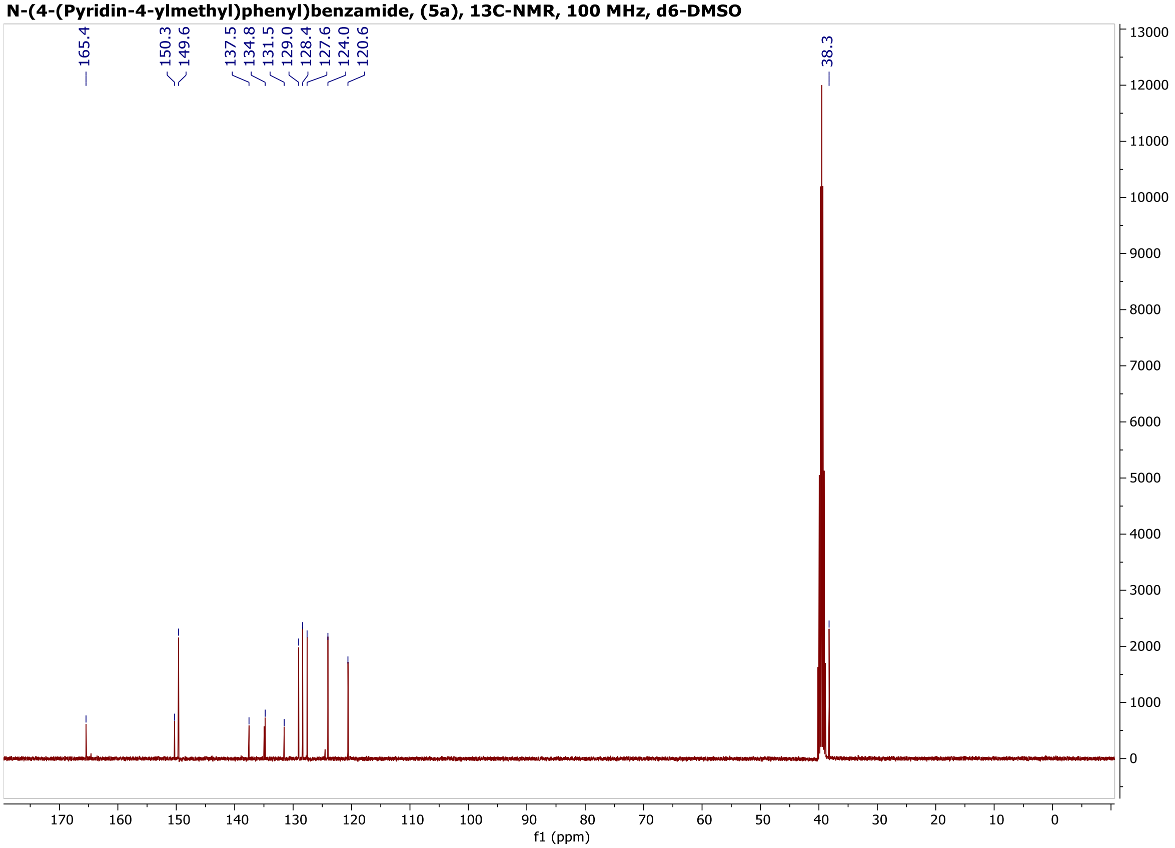

**N-(4-(Pyridin-4-ylamino)phenyl)benzamide, (5b), 1H-NMR, 500 MHz, d6-DMSO**

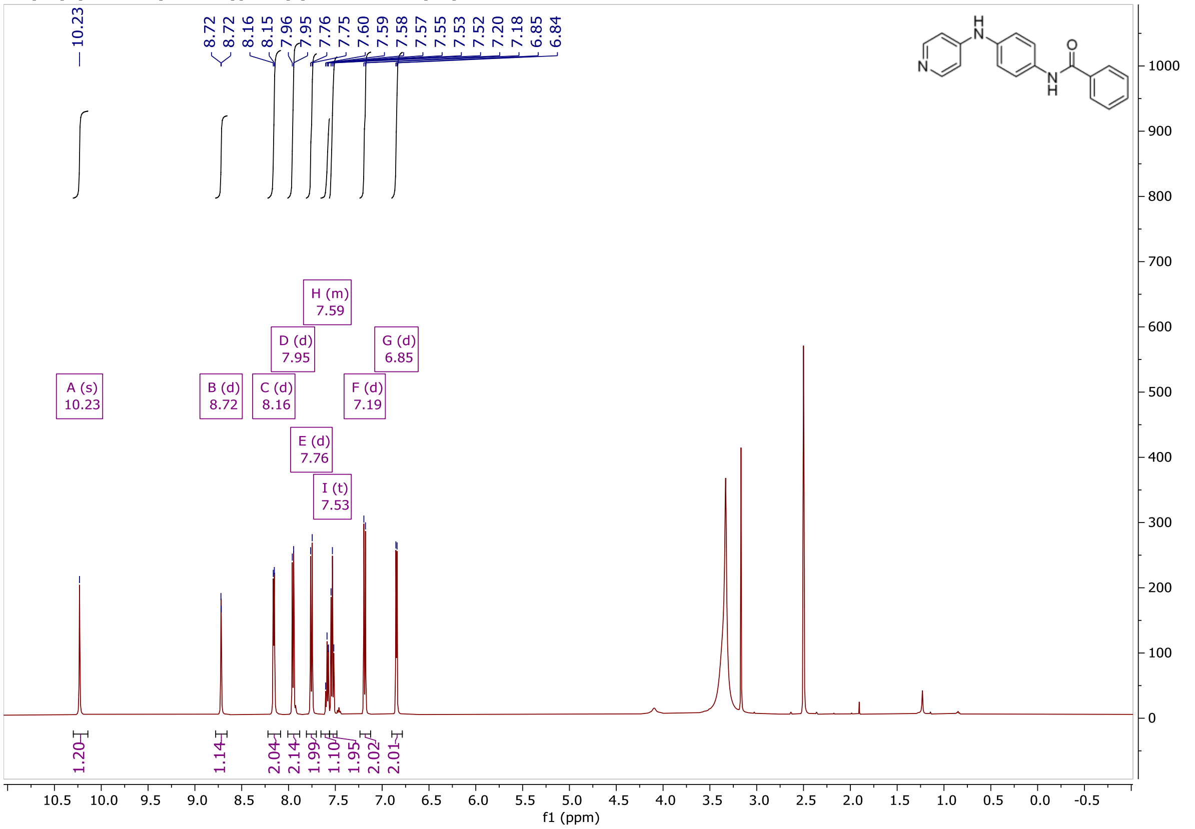


**N-(4-(Pyridin-4-ylamino)phenyl)benzamide, (5b), 13C-NMR, 125 MHz, d6-DMSO)**

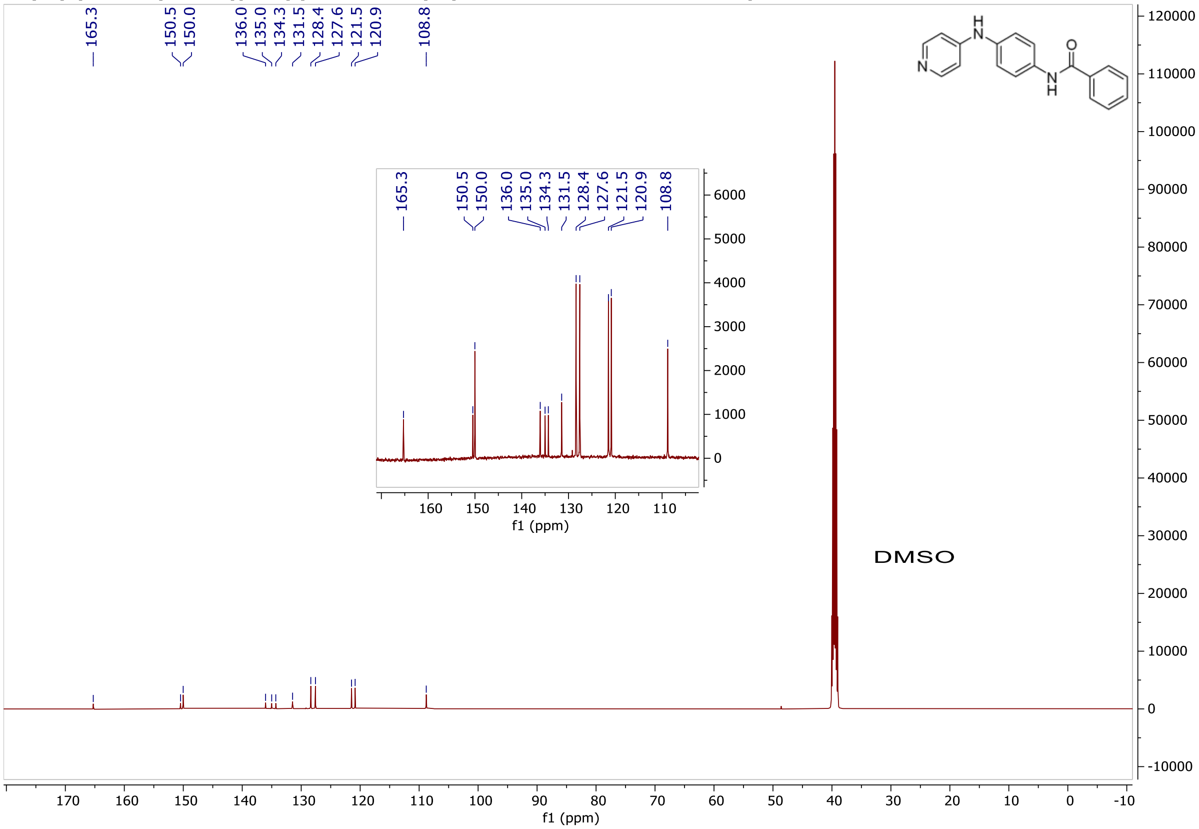

**N-(4-(Methyl(pyridin-4-yl)amino)phenyl)benzamide, (5c), 1H-NMR, 400 MHz, CDCl3**

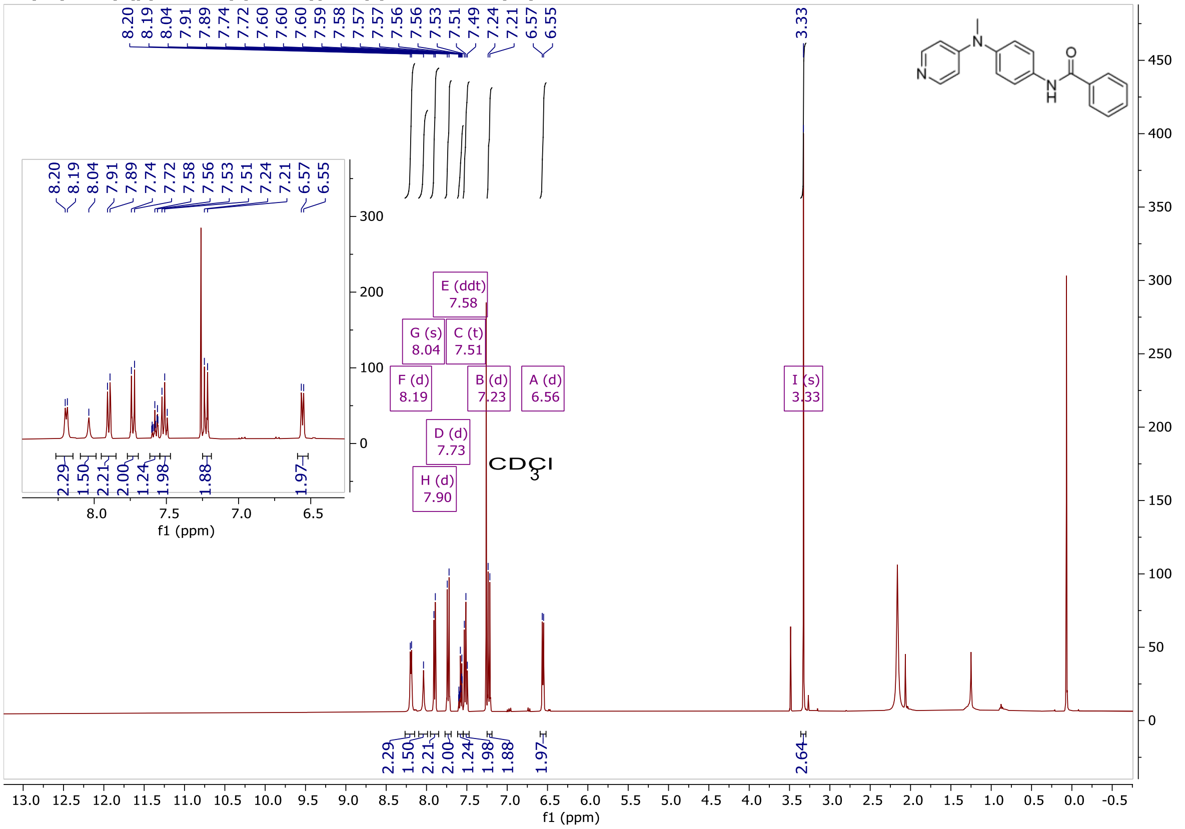

**N-(4-(Methyl(pyridin-4-yl)amino)phenyl)benzamide, (5c), 13C-NMR, 100 MHz, CDCl3**

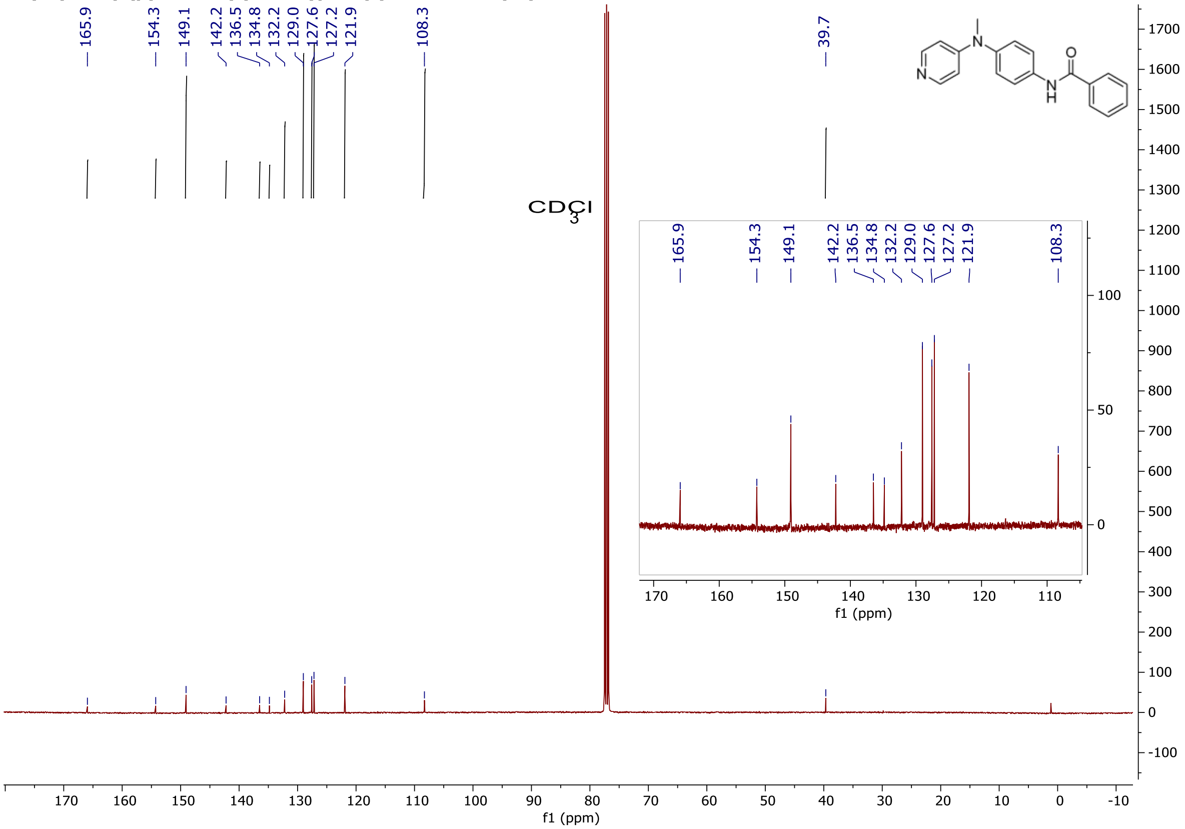

**(S)-2-(Benzylamino)-3-(1H-indol-3-yl)-N-(4-(pyridin-4-ylmethyl)phenyl)propanamide, (5d), 1H-NMR, 500 MHz, CDCl3**

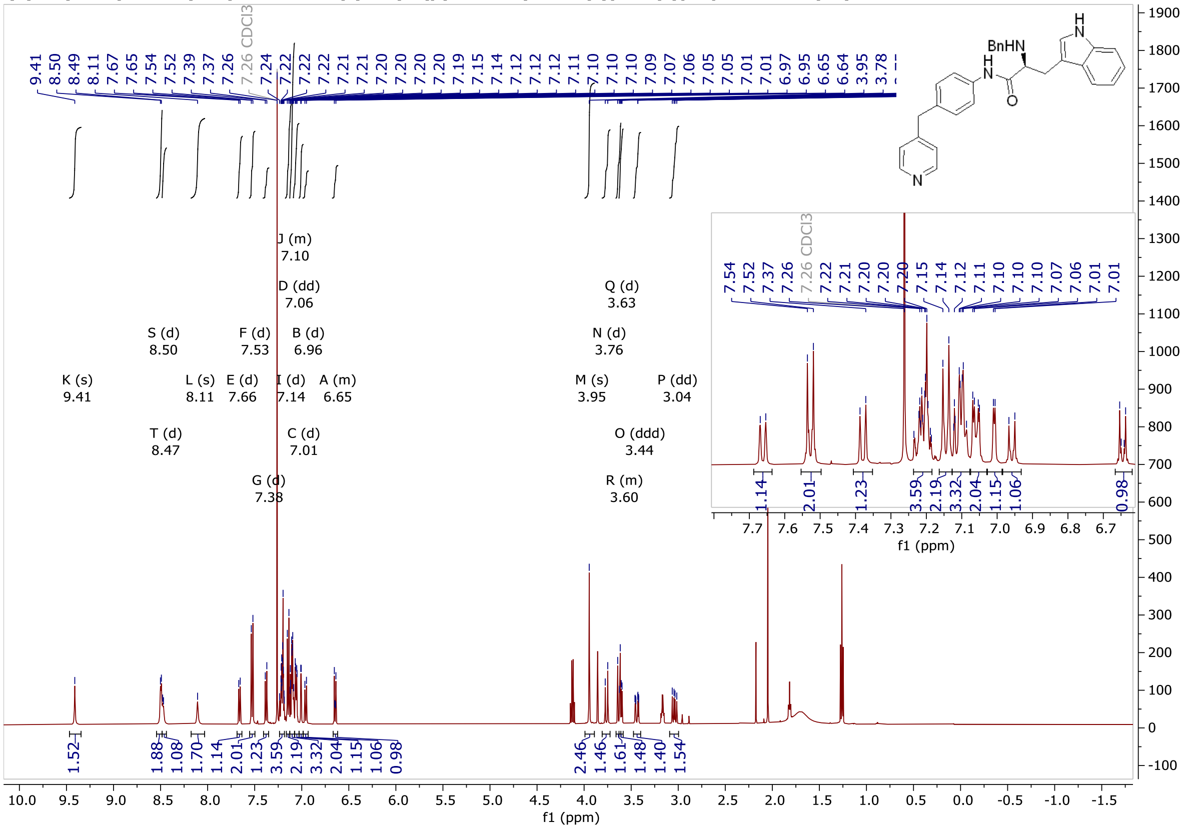

**(S)-2-(Benzylamino)-3-(1H-indol-3-yl)-N-(4-(pyridin-4-ylmethyl)phenyl)propanamide, (5d), 13C-NMR, 125 MHz, CDCl3**

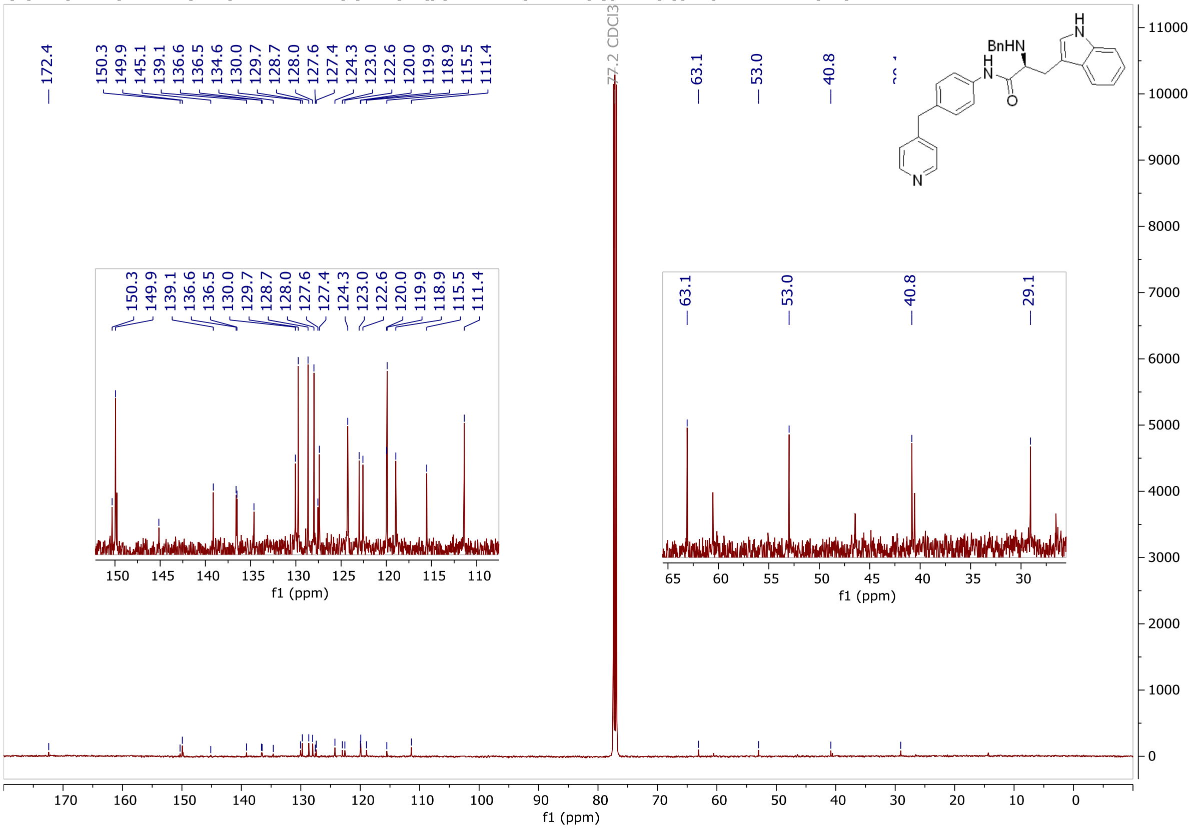

**N-((1H-Indol-5-yl)methyl)-3-(pyridin-4-ylmethyl)benzamide, (5e), 1H-NMR, 500 MHz, CDCl3**

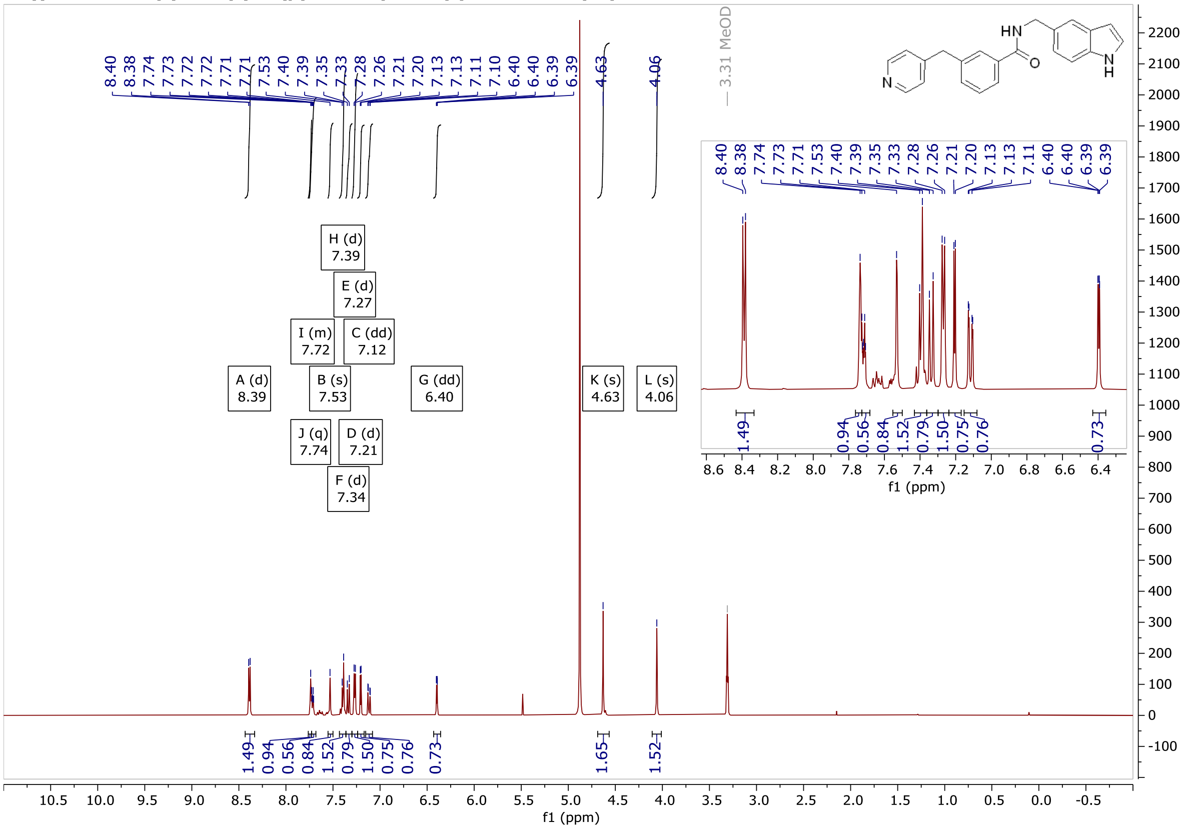


**N-((1H-Indol-5-yl)methyl)-3-(pyridin-4-ylmethyl)benzamide, (5e), 13C-NMR, 125 MHz, CDCl3**

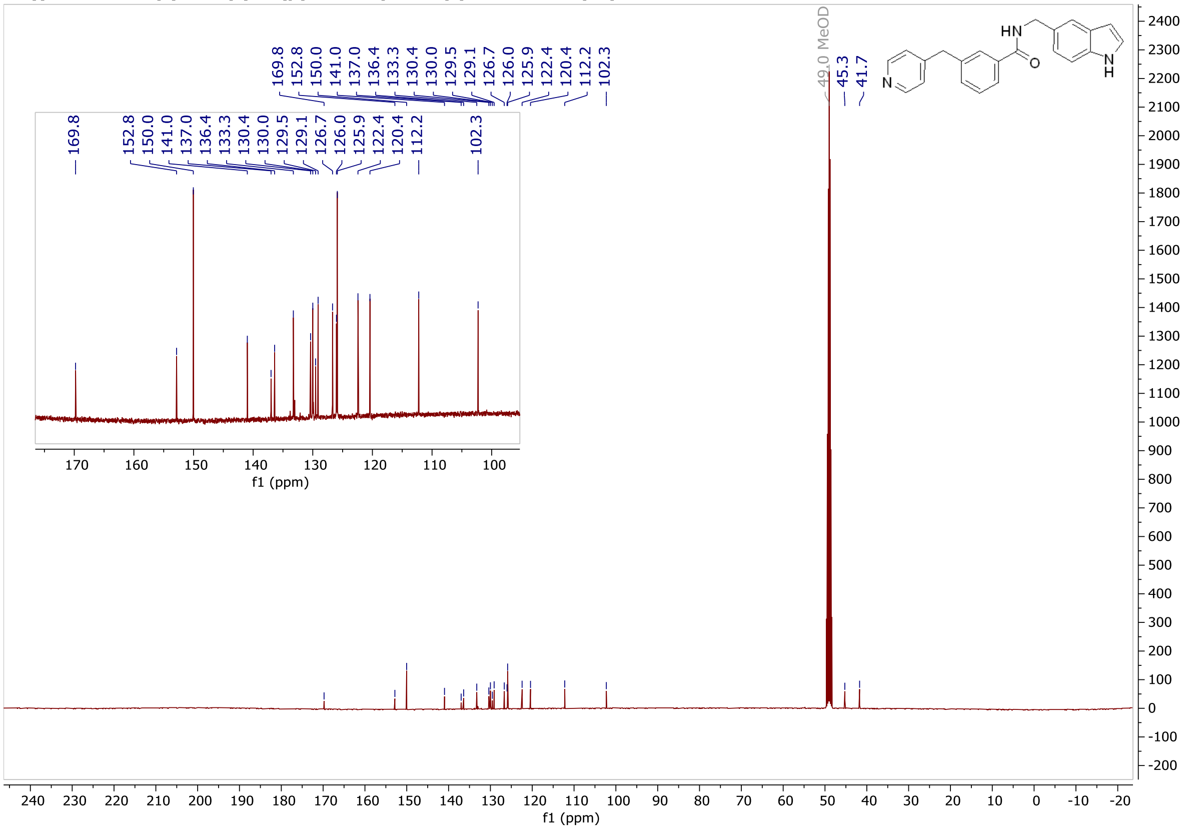

**N-((1H-Indol-5-yl)methyl)-4-(pyridin-4-ylmethyl)benzamide, (5f), 1H-NMR, 500 MHz, MeOD**

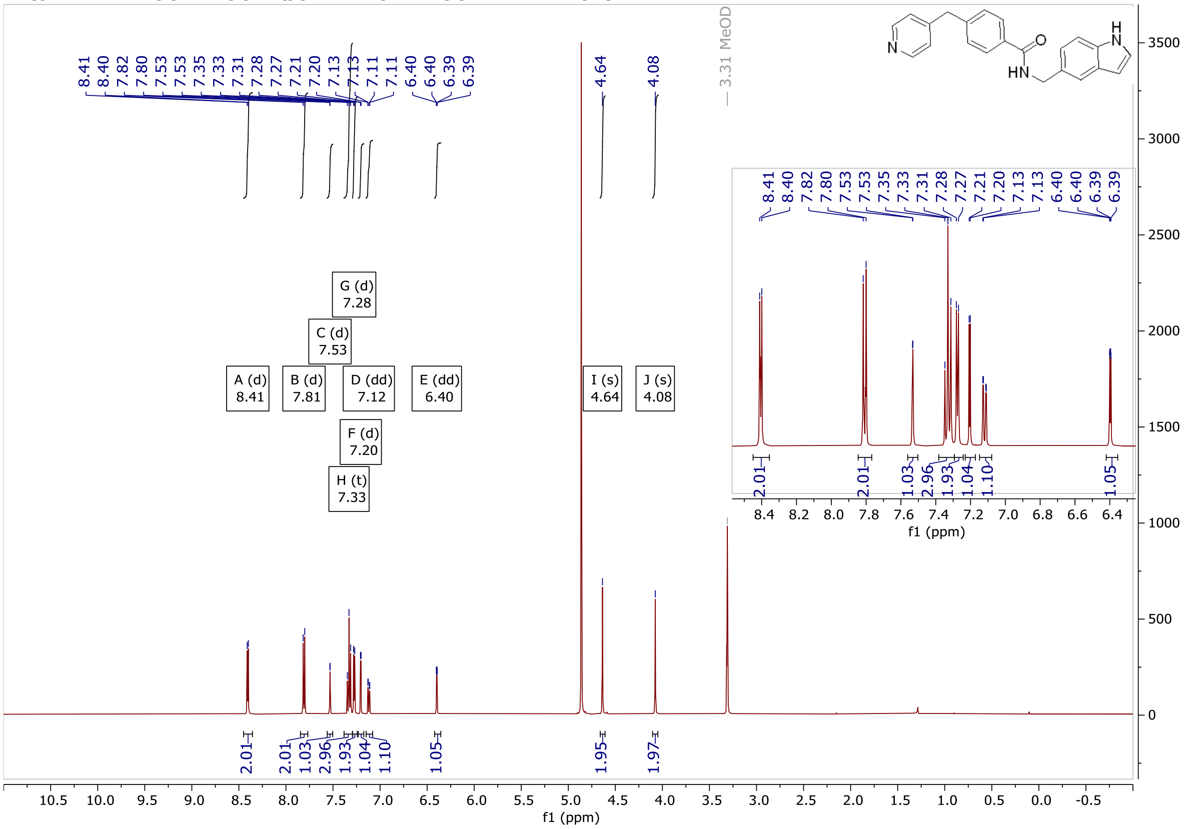

**N-((1H-Indol-5-yl)methyl)-4-(pyridin-4-ylmethyl)benzamide, (5f), 13C-NMR, 125 MHz, MeOD**

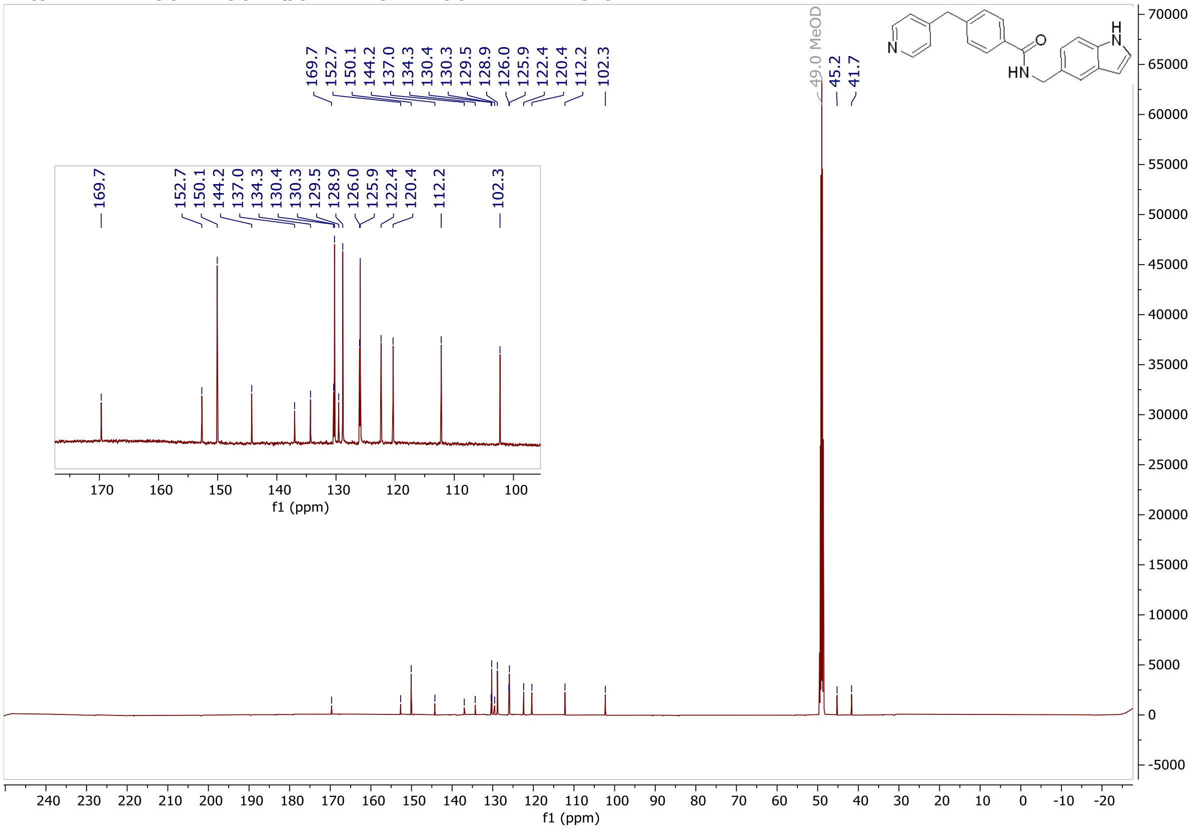

**(R)-N-(1-Phenylethyl)-3-(pyridin-4-ylmethyl)benzamide, (5h), 500 MHz, MeOD**

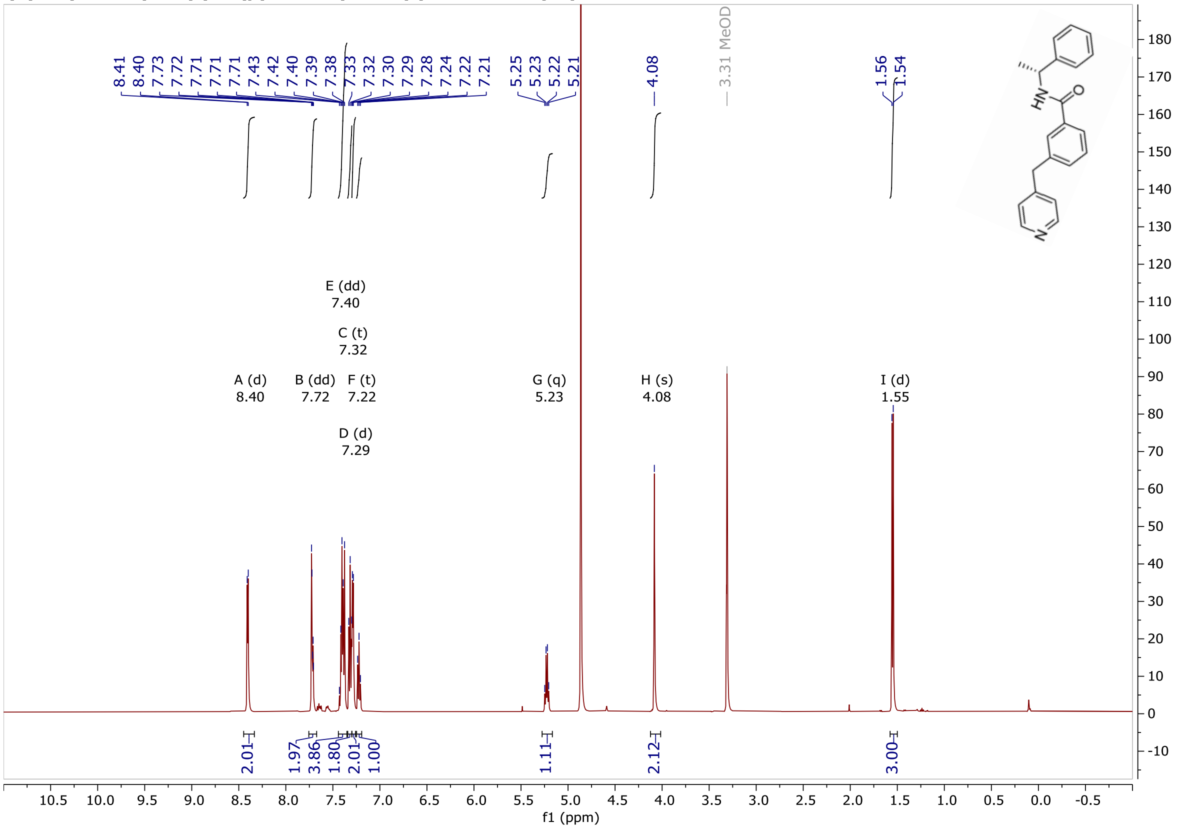


**(R)-N-(1-Phenylethyl)-3-(pyridin-4-ylmethyl)benzamide, (5h), 125 MHz, MeOD**

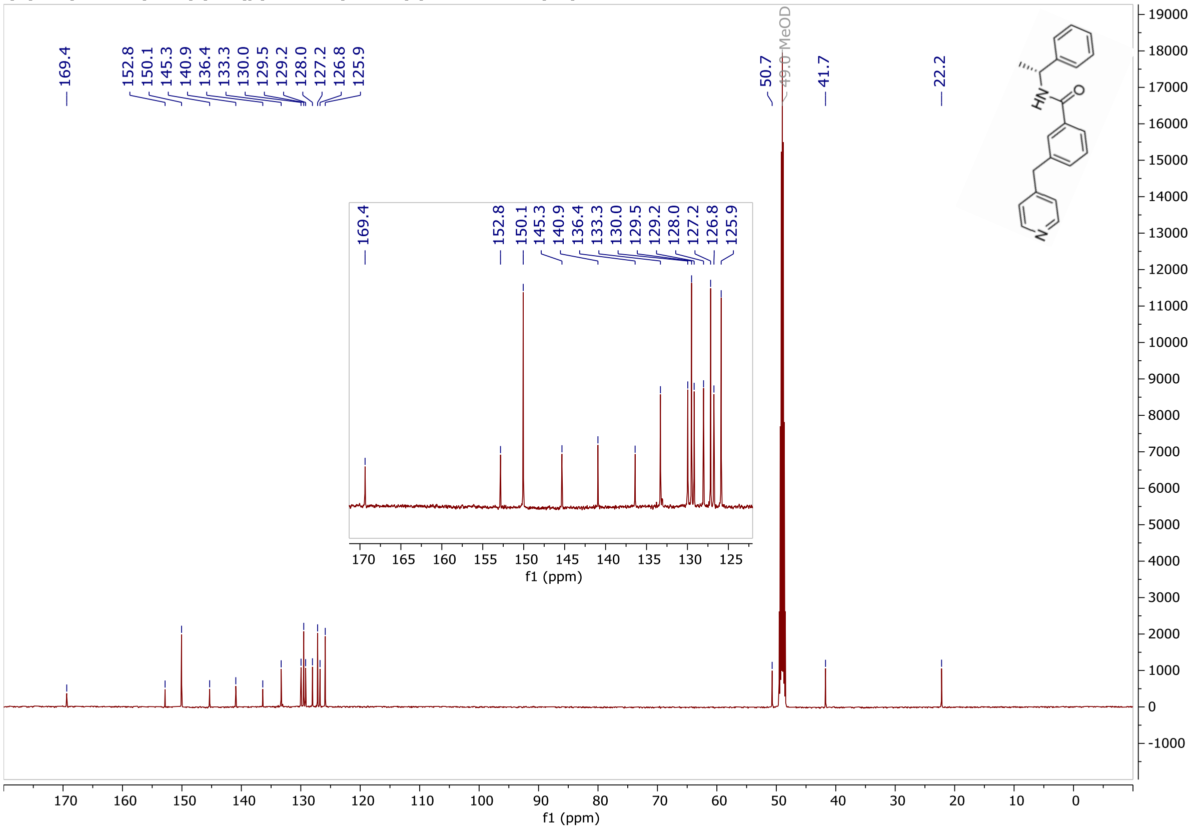

**(R)-N-(1-Phenylethyl)-4-(pyridin-4-ylmethyl)benzamide, (5i), 500 MHz, MeOD**

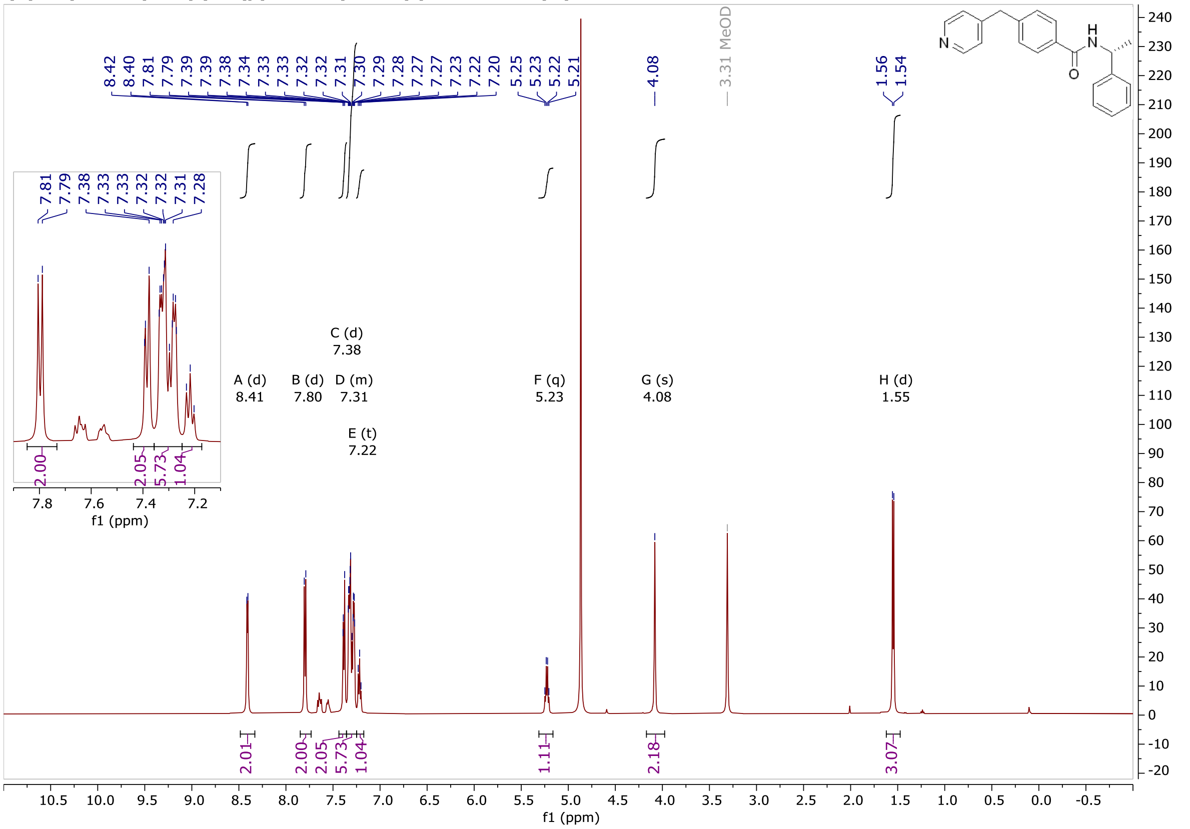


**(R)-N-(1-Phenylethyl)-4-(pyridin-4-ylmethyl)benzamide, (5i), 500 MHz, MeOD**

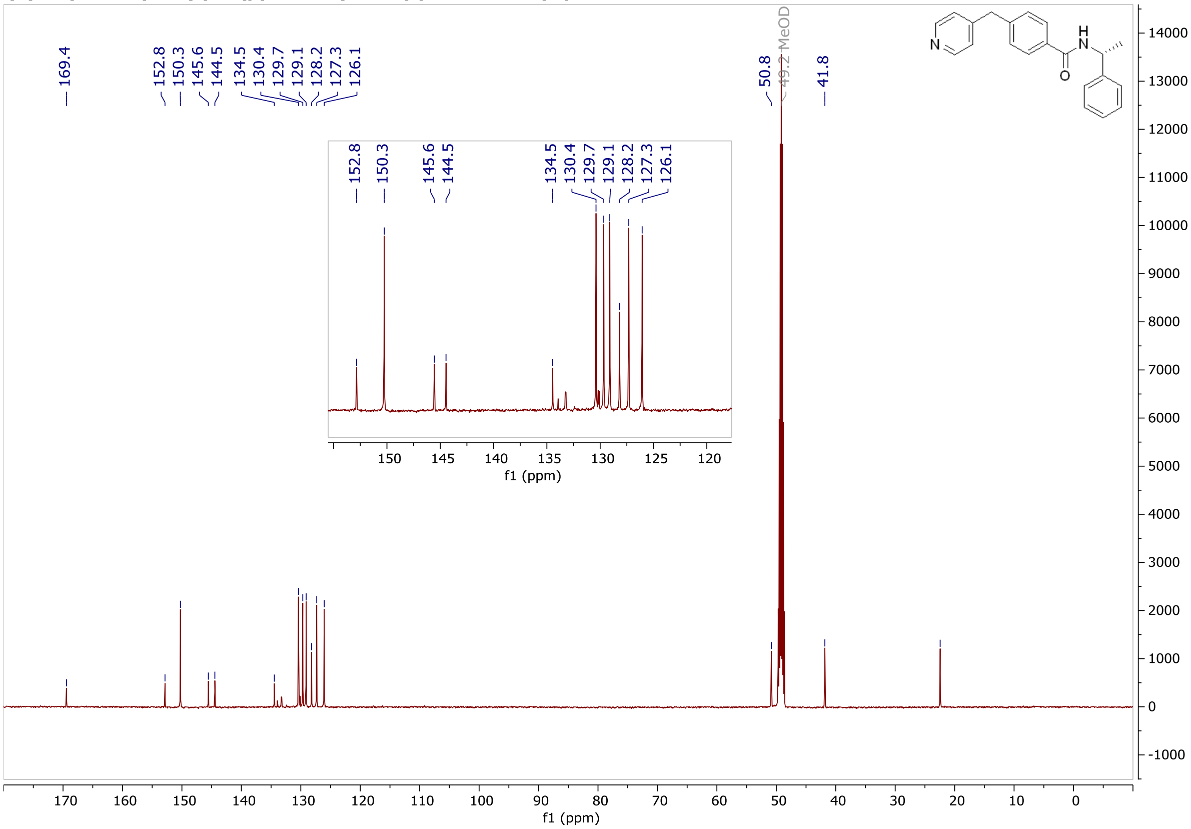

**N-(4-(Pyridin-4-ylmethyl)phenyl)benzenesulfonamide, (5j), 1H-NMR, 400 MHz, d6-DMSO**

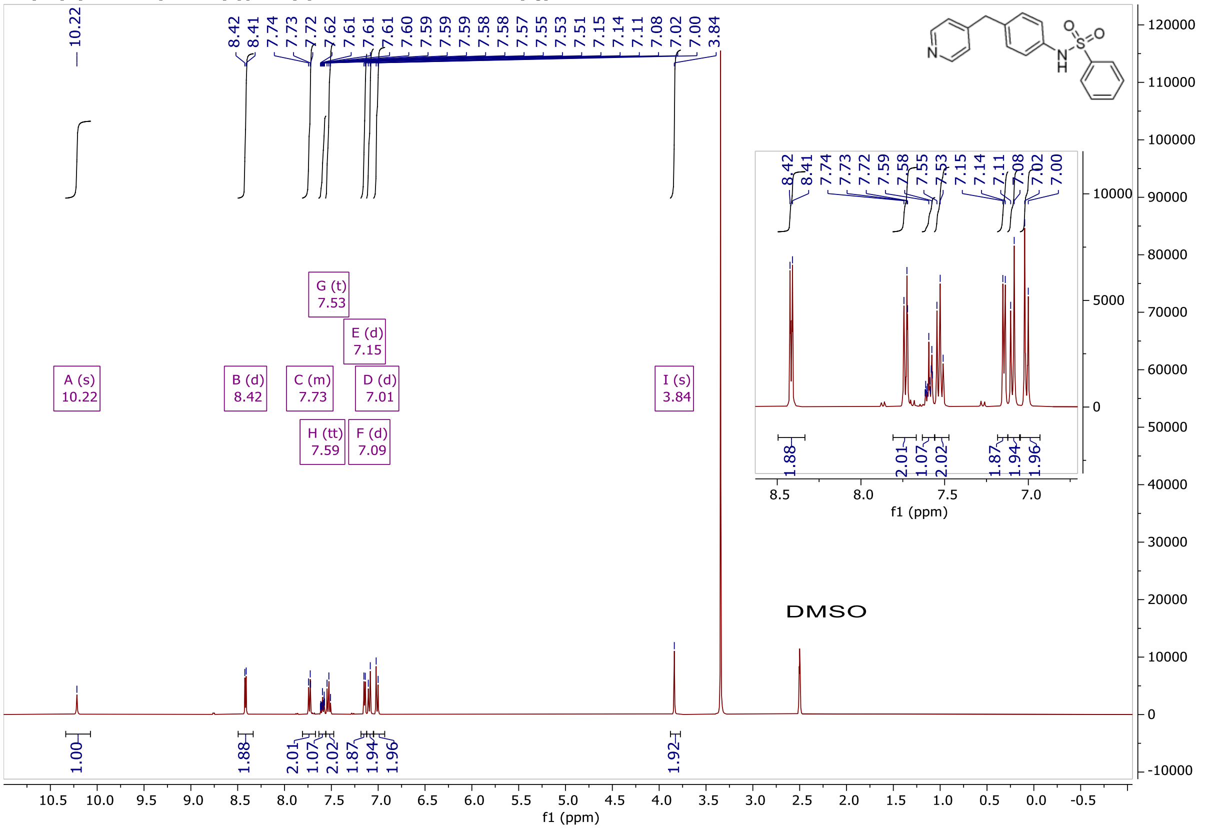


**N-(4-(Pyridin-4-ylmethyl)phenyl)benzenesulfonamide, (5j), 13C-NMR, 100 MHz, d6-DMSO**

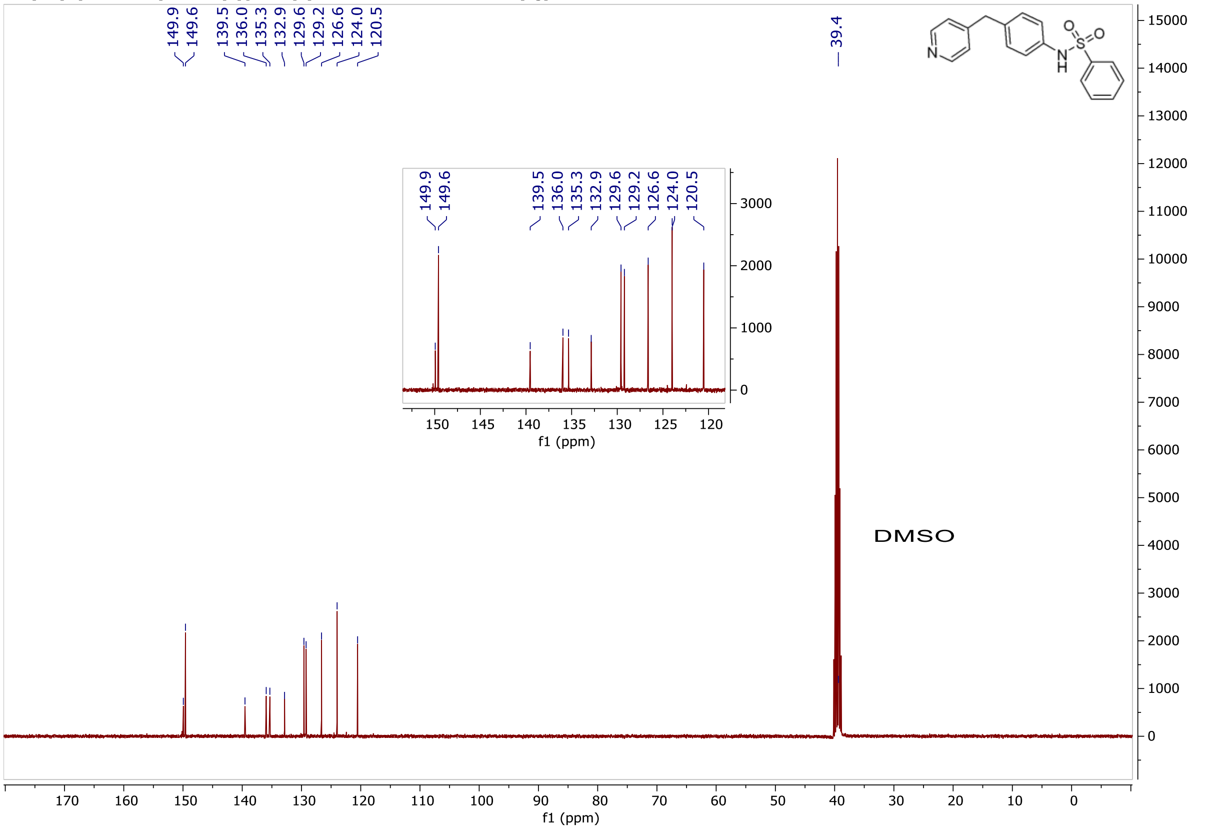

**N-(4-(Pyridin-4-ylmethyl)phenyl)-4-(trifluoromethoxy)benzenesulfonamide, (5k), 1H-NMR, CDCl3**

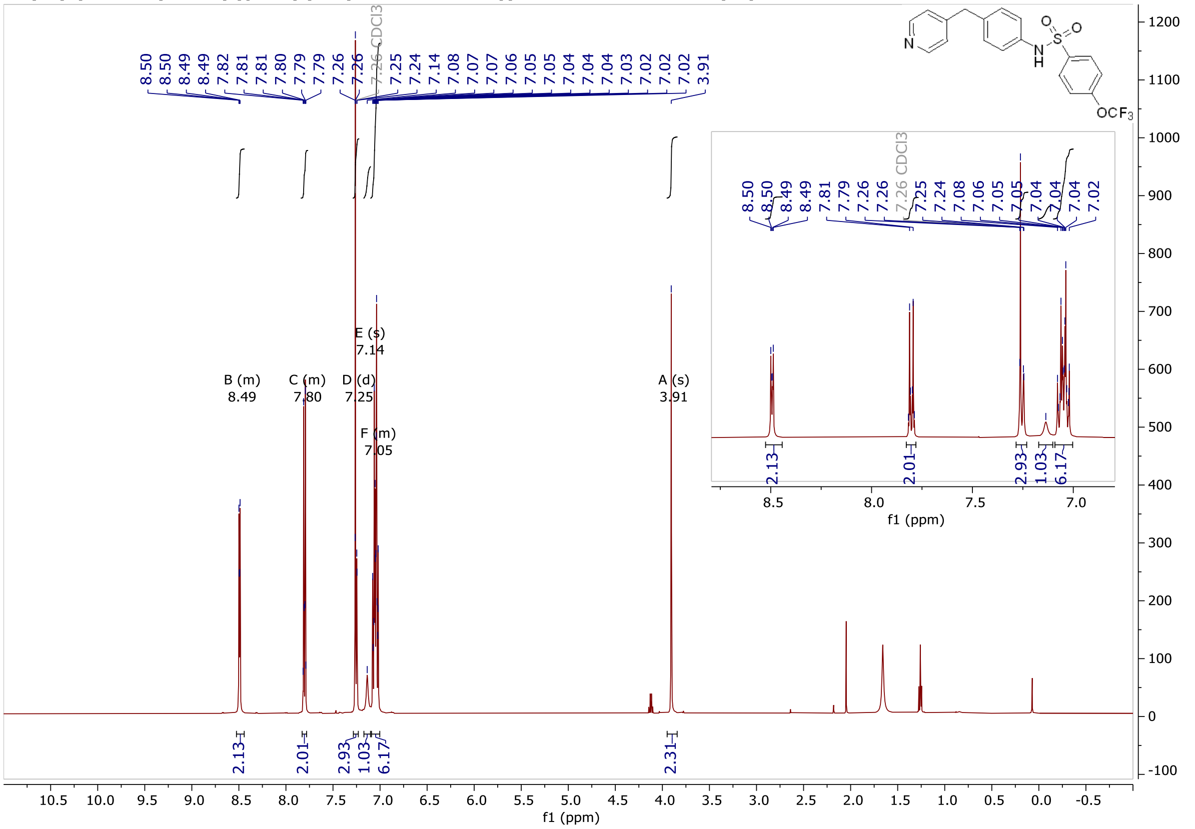

**N-(4-(Pyridin-4-ylmethyl)phenyl)-4-(trifluoromethoxy)benzenesulfonamide, (5k), 13C-NMR, CDCl3**

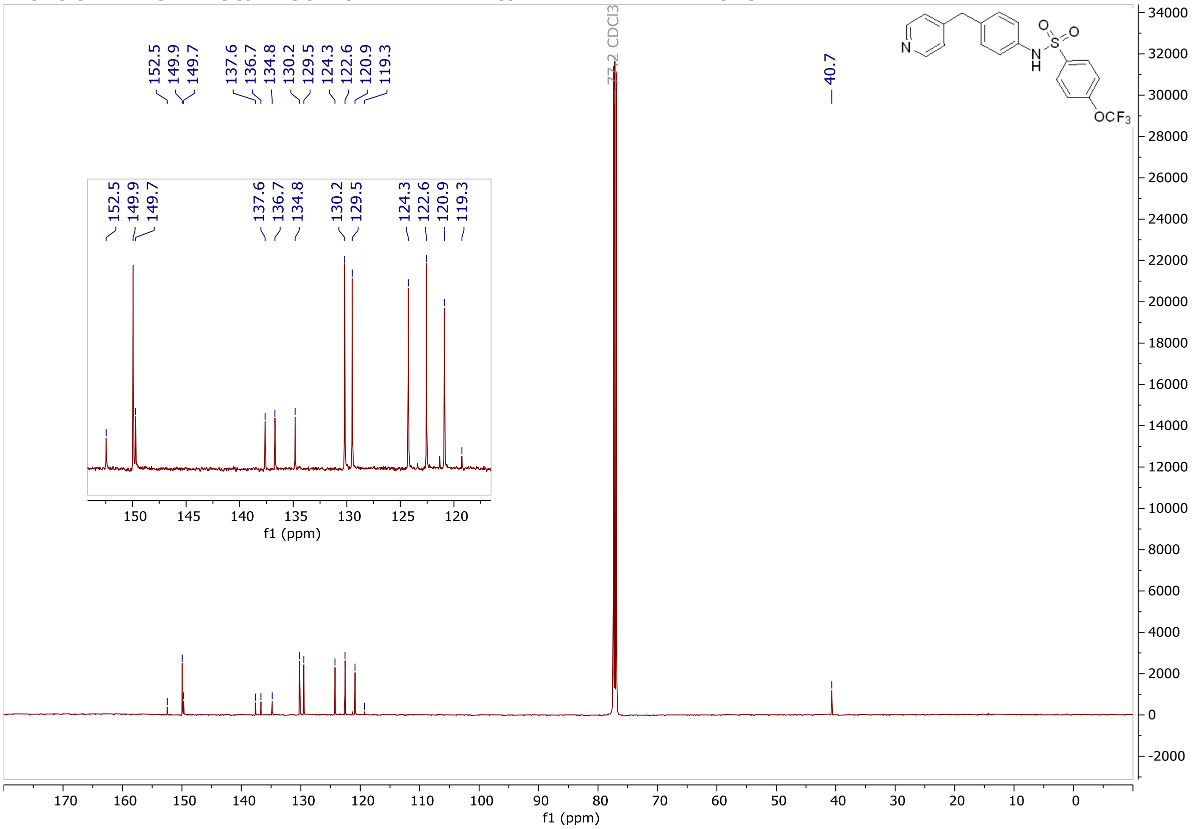

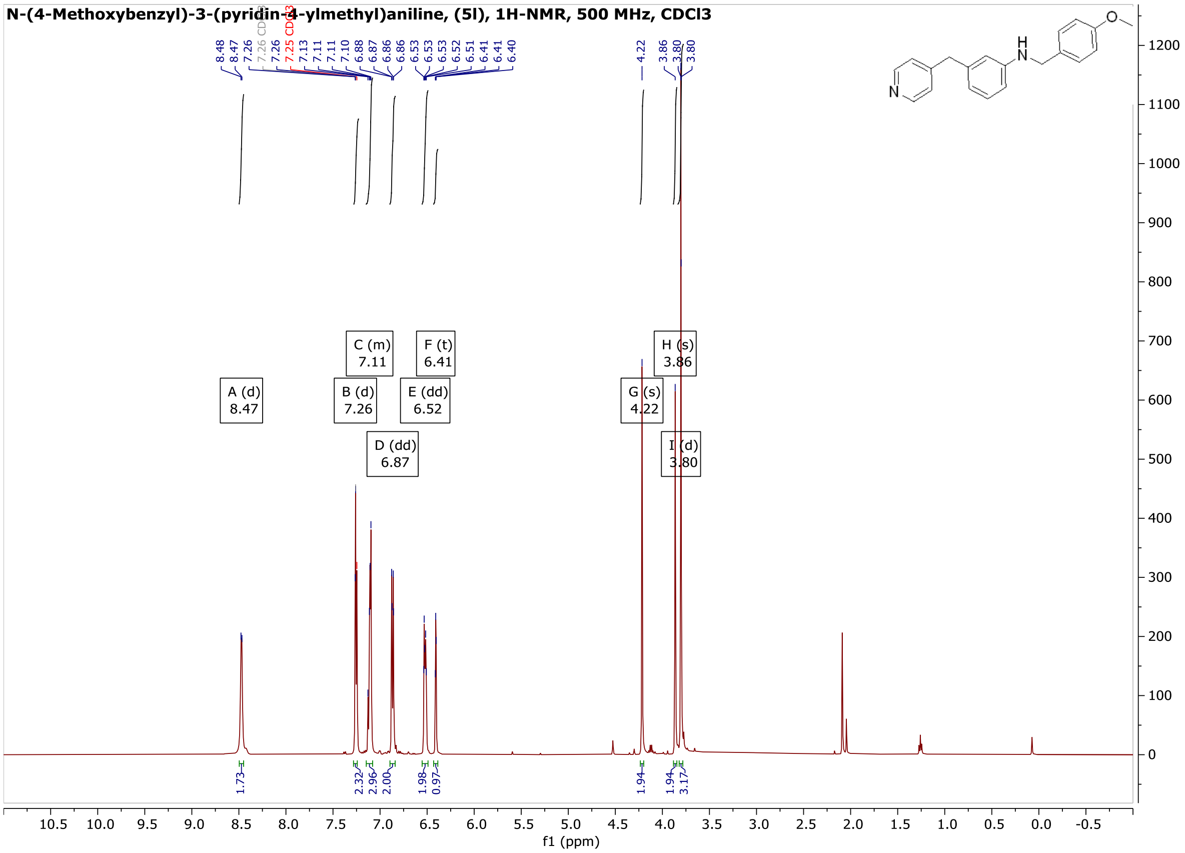
N-(4-Methoxybenzyl)-3-(pyridin-4-ylmethyl)aniline, (5l), 1H-NMR, 500 MHz, CDCl3
A (d) 8.47
B (d) 7.26
C (m) 7.11
D (dd) 6.87
E (dd) 6.52
F (t) 6.41
G (s) 4.22
H (s) 3.86
I (d) 3.80
f1 (ppm)

N-(4-Methoxybenzyl)-3-(pyridin-4-ylmethyl)aniline, (5l), 1H-NMR, 500 MHz, CDCl3

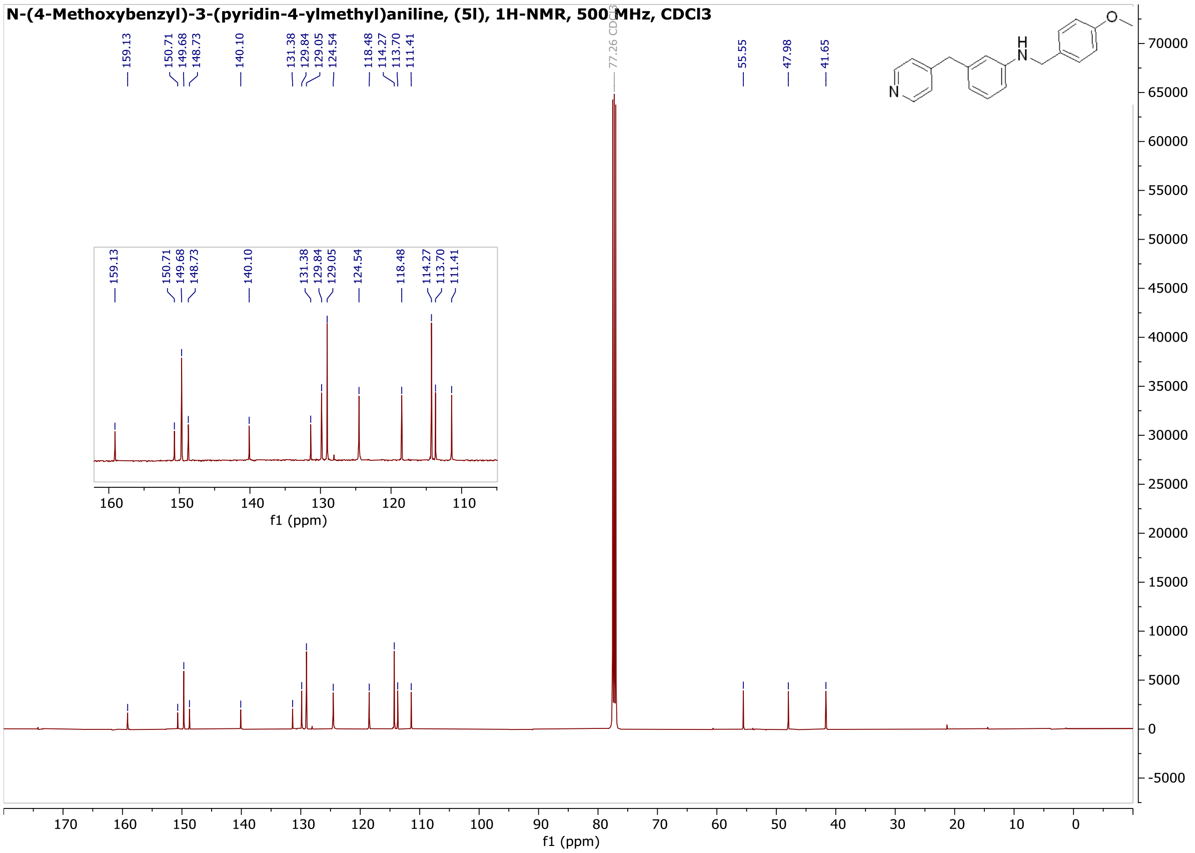


N-(4-Methoxybenzyl)-4-(pyridin-4-ylmethyl)aniline, (5m), 1H-NMR, 500 MHz, CDCl3

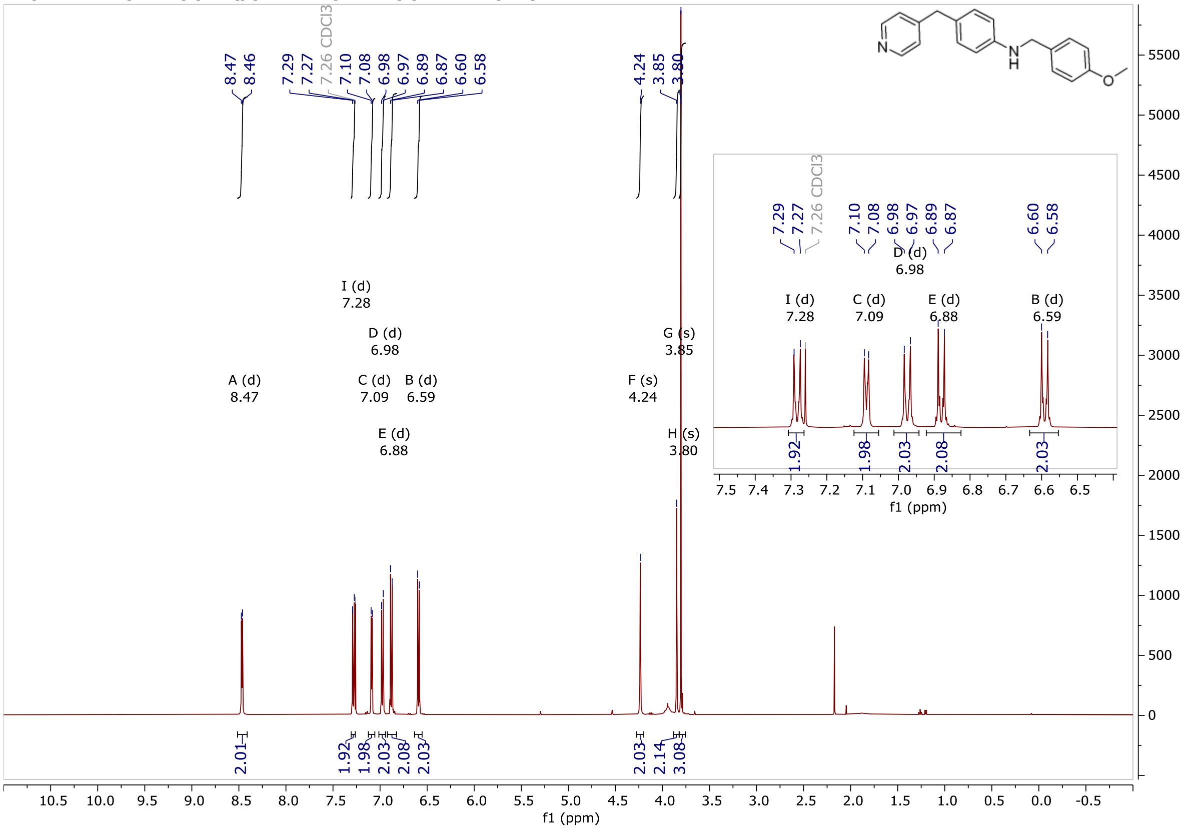

**N-(4-Methoxybenzyl)-4-(pyridin-4-ylmethyl)aniline, (5m), 13C-NMR, 125 MHz, CDCl3**

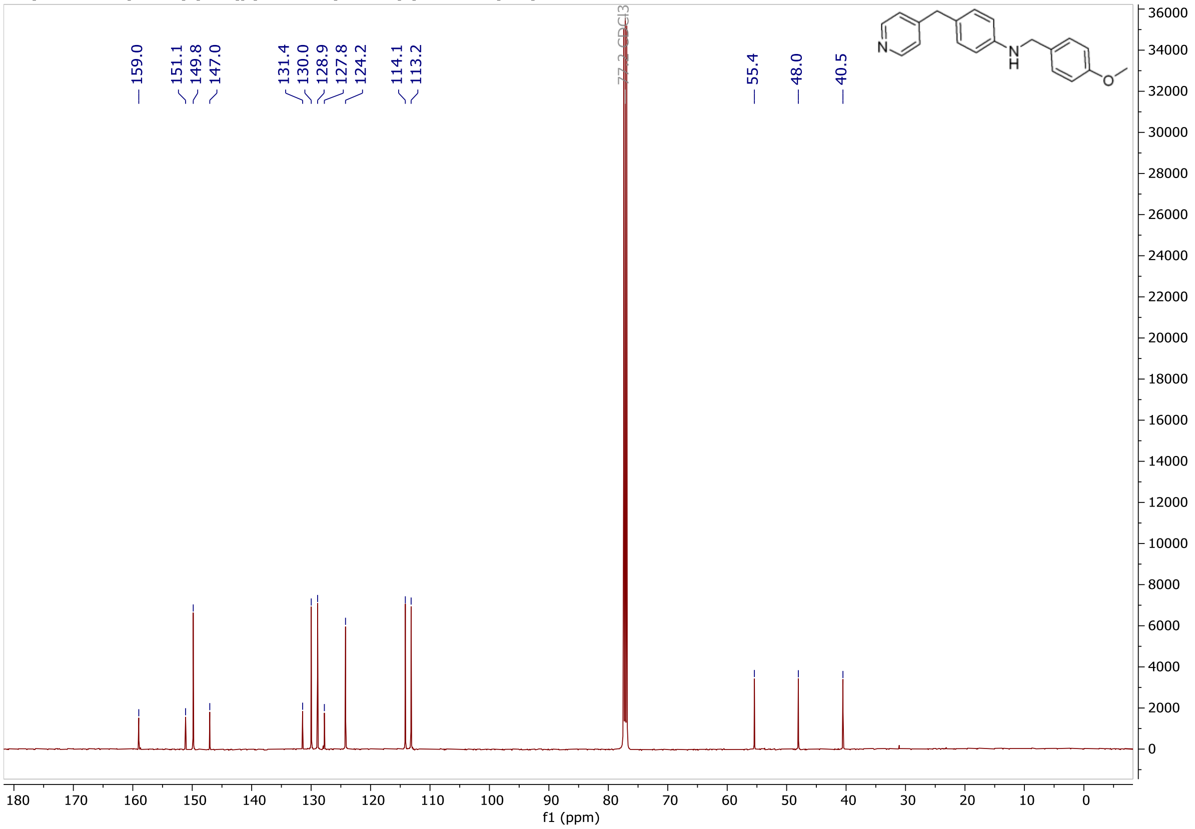


**N1-(4-Methoxybenzyl)-N4-methyl-N4-(pyridin-4-yl)benzene-1,4-diamine, (5n), 1H-NMR, 500 MHz, CDCl3**

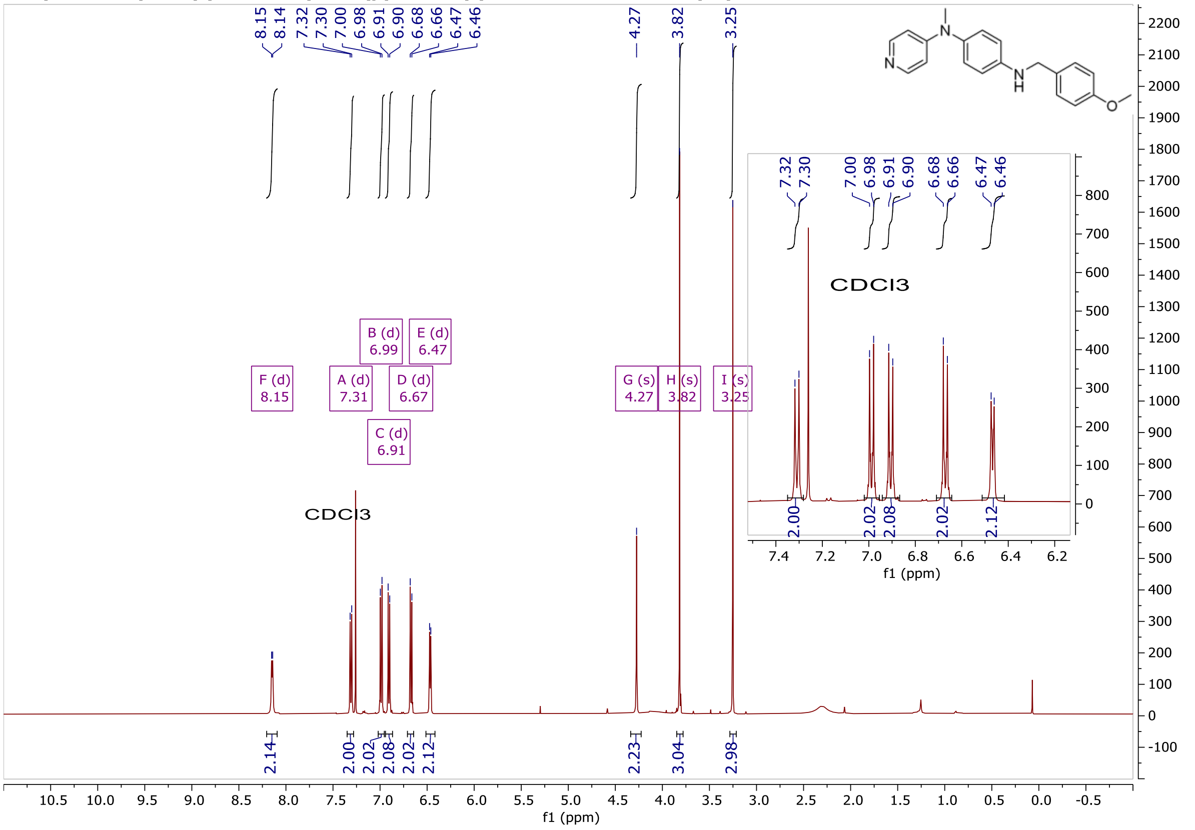

**N1-(4-Methoxybenzyl)-N4-methyl-N4-(pyridin-4-yl)benzene-1,4-diamine, (5n), 13C-NMR, 125 MHz, CDCl3**

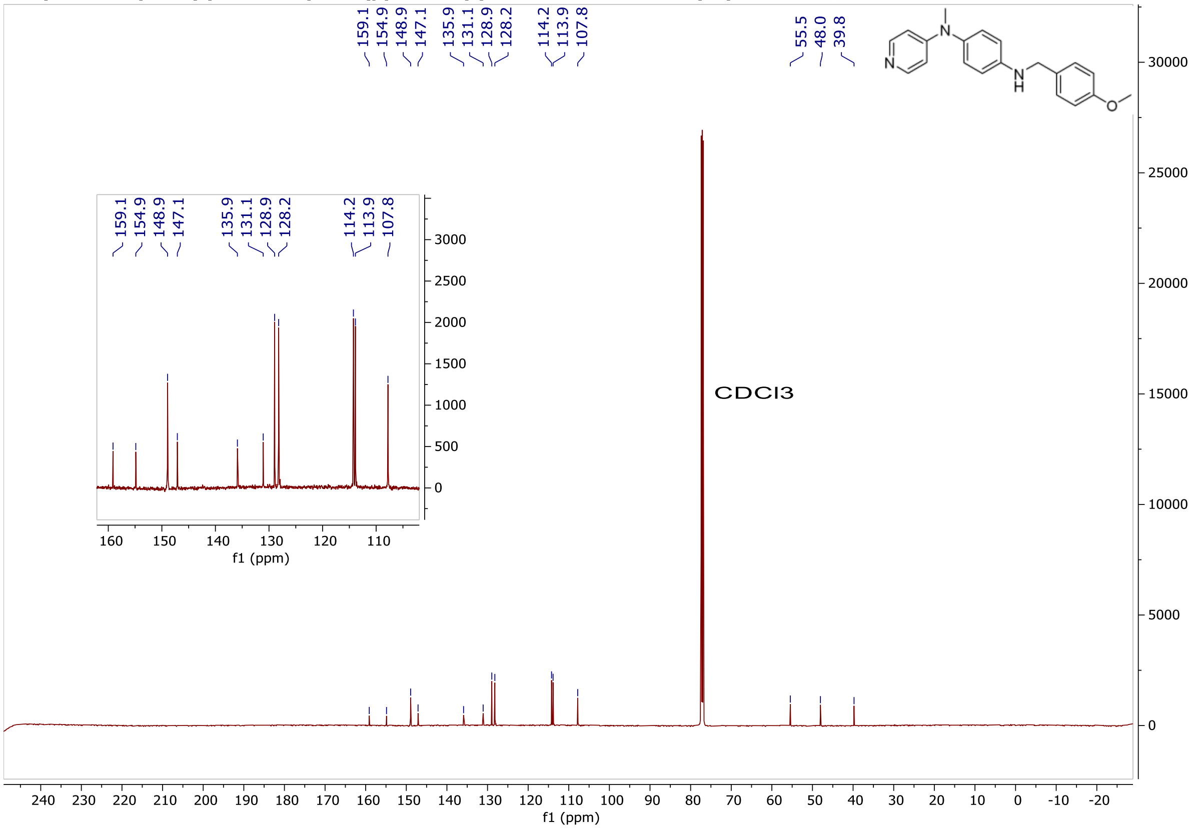

N-(1-(4-Methoxyphenyl)ethyl)-4-(pyridin-4-ylmethyl)aniline, (5o), 1H-NMR, 400 MHz, CDCl3

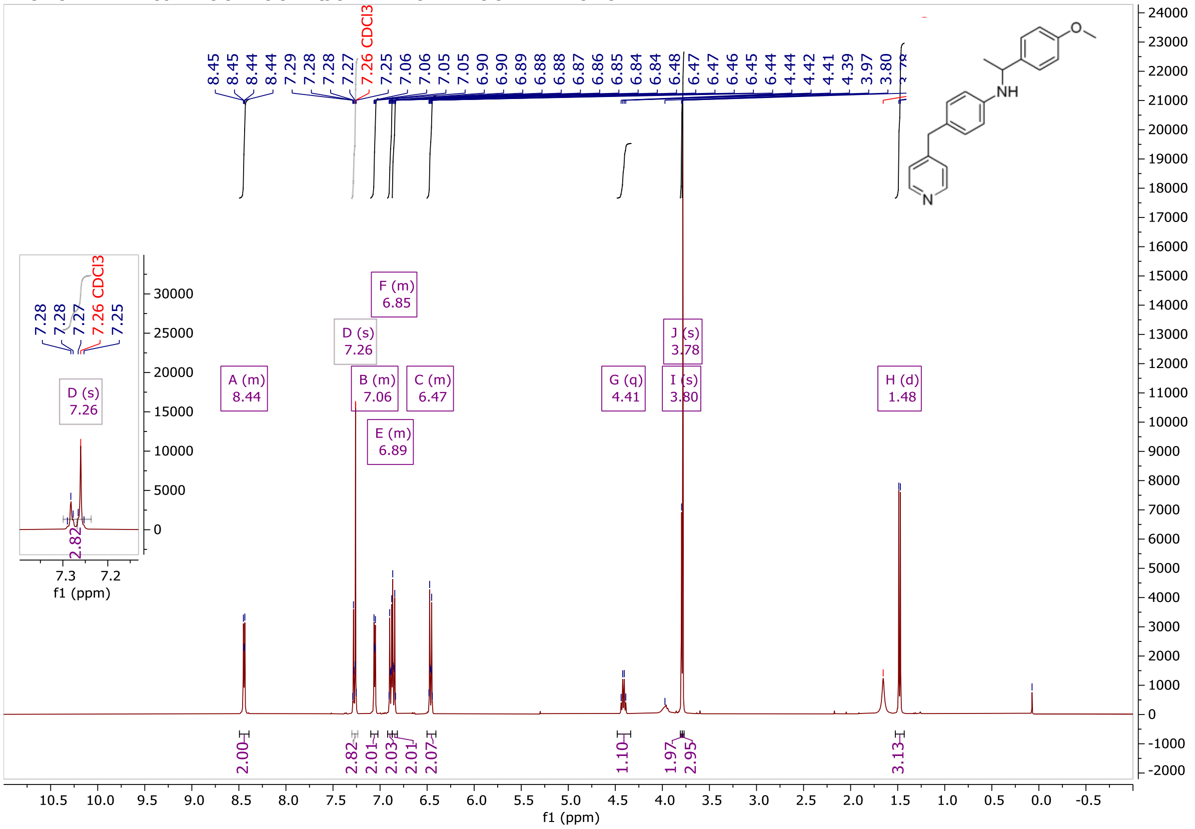


N-(1-(4-Methoxyphenyl)ethyl)-4-(pyridin-4-ylmethyl)aniline, (5o), 13C-NMR, 100 MHz, CDCl3

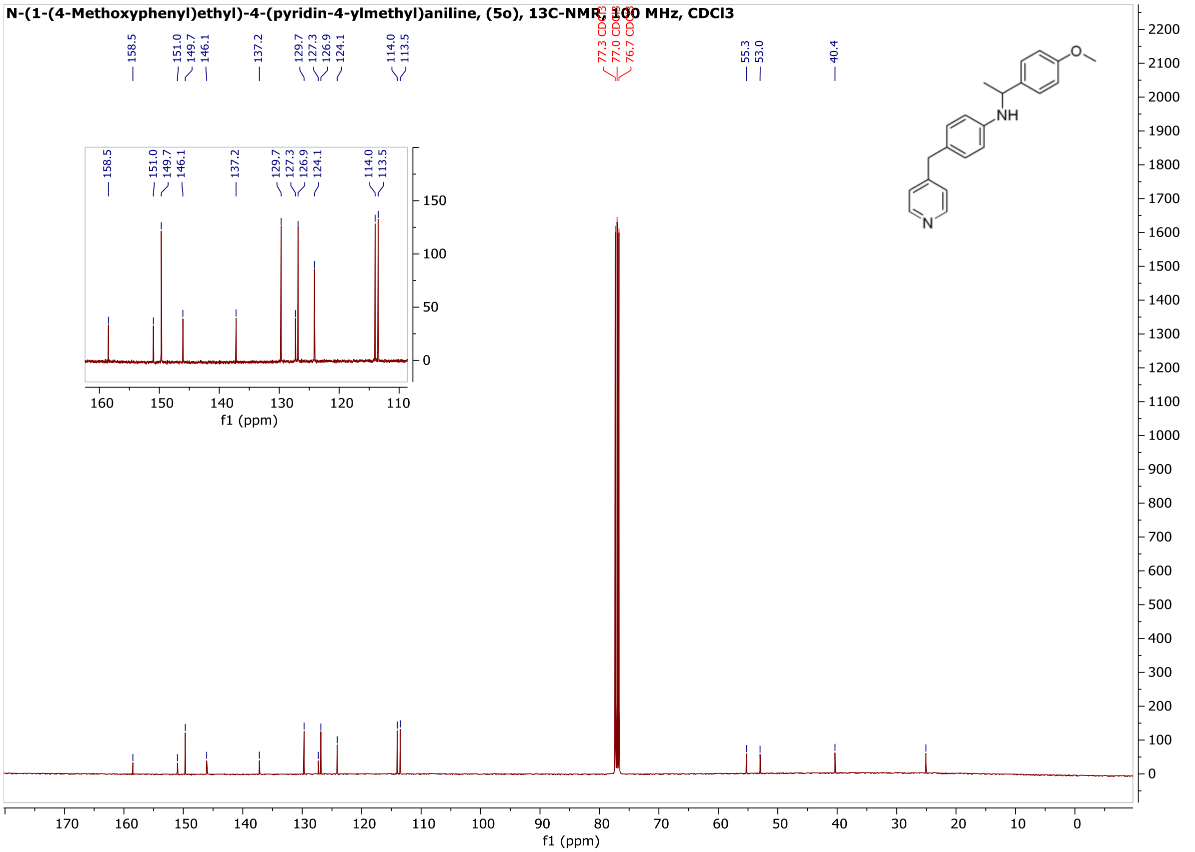

**N-(3-(Methylsulfonyl)benzyl)-4-(pyridin-4-ylmethyl)aniline, (5p), 500 MHz, CDCl3**

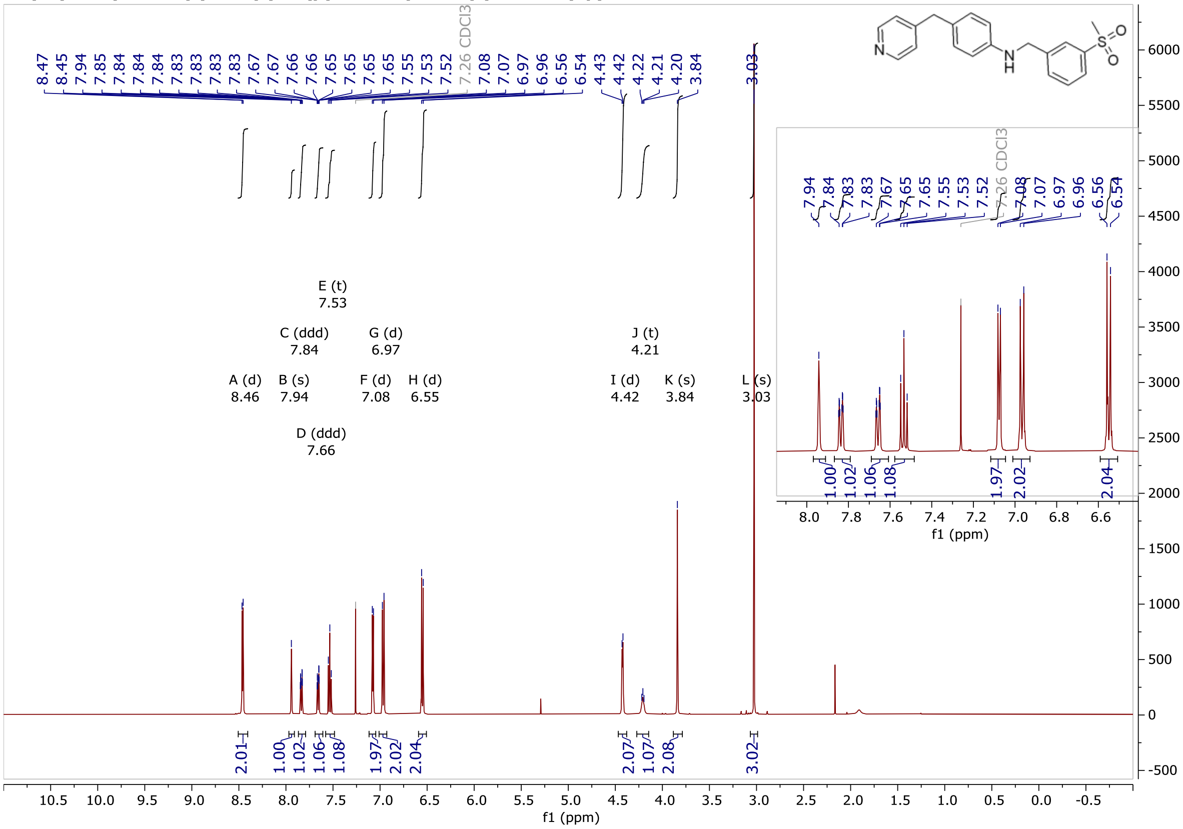

**N-(3-(Methylsulfonyl)benzyl)-4-(pyridin-4-ylmethyl)aniline, (5p), 125 MHz, CDCl3**

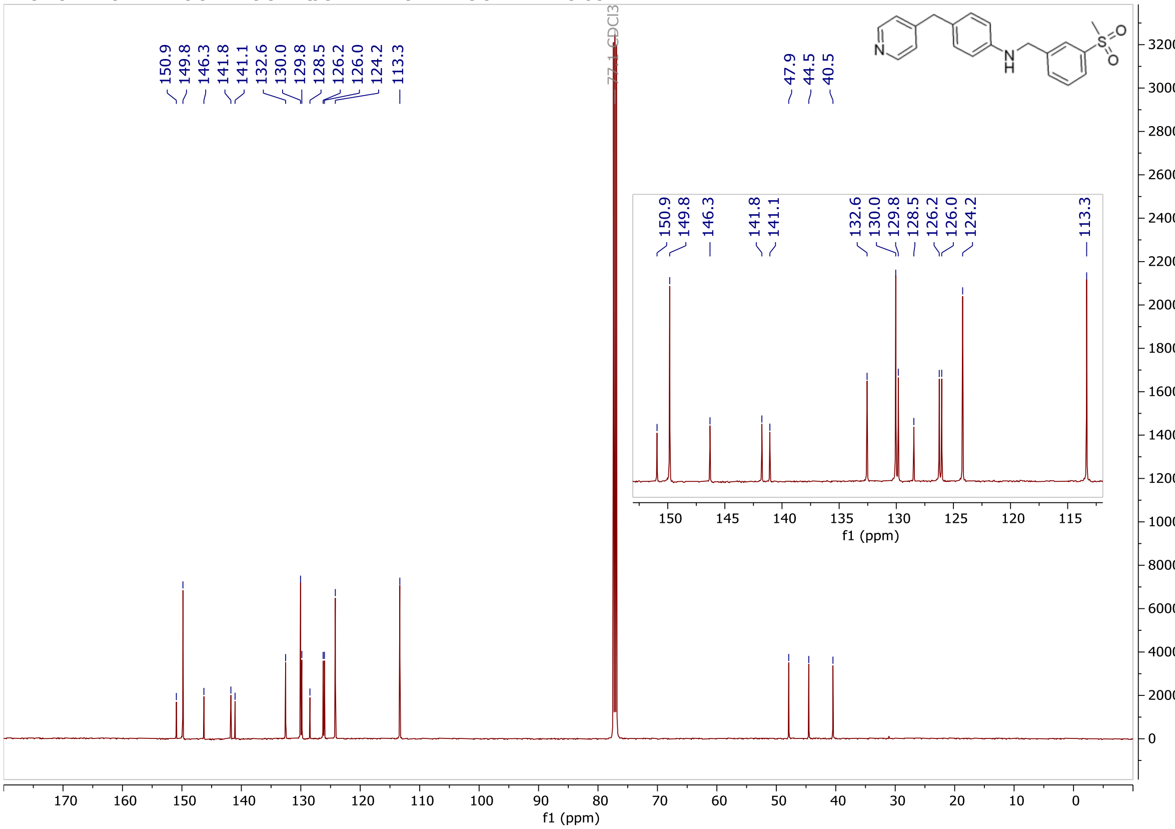
