## Supporting Information for "Fragment-based development of small molecule inhibitors targeting *Mycobacterium tuberculosis* cholesterol metabolism"

[1]Yusuf Hamied Department of Chemistry, University of Cambridge, Lensfield Road, Cambridge, CB2 1EW, UK. [2]Centre for Synthetic Biology of Fine and Specialty Chemicals (SYNBIOCHEM), Manchester Institute of Biotechnology, University of Manchester, 131 Princess Street, Manchester, M1 7DN, UK. [3]Tuberculosis Research Section, Laboratory of Clinical Infectious Diseases, National Institute of Allergy and Infectious Disease, National Institutes of Health, Bethesda, Maryland, USA. [4]Global Health R&D, GSK, Severo Ochoa, 2, 28760 Tres Cantos, Spain. [5]Manchester Protein Structure Facility (MPSF), Manchester Institute of Biotechnology, University of Manchester, Manchester, M1 7DN, UK. [6]Department of Chemistry, Manchester Institute of Biotechnology, University of Manchester, 131 Princess Street, Manchester, M1 7DN, UK

[#]corresponding authors

**Table S1.** Screen of focused library of heme-binding fragments against purified *Mtb* P450s by UV-vis spectroscopy. Interaction between fragment (1 mM) and P450 (4-6 µM) was quantified from the shift in the Soret band ($\Delta\lambda_{max}$, nm) of each enzyme's absorbance spectrum, relative to DMSO. NB: All P450s tested are predominantly low-spin at resting state ($\lambda_{max}$ (DMSO) ~ 416 – 421 nm), except CYP125, which is predominantly high spin ($\lambda_{max}$ (DMSO) = 393 nm). Consequently, fragments that bind and stabilize the P450 low-spin state induce a significantly larger $\Delta\lambda_{max}$ in the spectrum of CYP125 compared to the other P450s analyzed. ND: not determined, due to optical interference, or insolubility in P450-optimized buffer.

| Fragment | SMILES | CYP125 | CYP142 | CYP124 | CYP121 | CYP126 | CYP143 | CYP144 |
|---|---|---|---|---|---|---|---|---|
| | Soret $\lambda_{max}$ (nm) - DMSO | 393 | 418.5 | 419 | 417 | 418 | 416 | 421 |
| 1 | OC(=O)c1ccnnc1 | 0 | 0 | 0 | -1 | 0 | 0.5 | 0 |
| 2 | C1(C2=CC=NN2)=CC=CC=N1 | 0 | 0 | 0 | 0 | 0 | 0 | 0 |
| 3 | NC1=NC2C(C=CC=C2)=N1 | 0 | 0 | 0 | 0 | 0 | 0.5 | 0 |
| 4 | C1(C2=CC=CC=C2)=NC=CN1 | 0 | 0 | 0 | 0 | 0 | 0 | 0 |
| 5 | c1ncnn1Cc1cc(ccc1)N | 0 | 1.5 | 1 | 2 | 0 | 1 | 0 |
| 6 | Nc1n(c2ccccc2)nc(C)c1 | 0 | 0 | 0 | 0 | 0 | 0.5 | 0 |
| 7 | CC(C=C1)=NNC1=O | 0 | 0 | 0 | 0 | 0 | 0 | 0 |
| 8 | c1(c2ccncc2)ccc(cc1)CO | 25 | 0 | 1 | 1 | 3 | 1 | 0 |
| 9 | n1c(cc(n1[H])c1occc1)CO | 0 | 0 | -1 | 0 | 0 | 0.5 | 0 |
| 10 | n1(c(ncc1)CO)Cc1ccccc1 | 0 | 3 | 1 | 0 | 0 | 1 | 0 |
| 11 | n1n(c(cc1c1ccccc1)N)C | 0 | 0 | 0 | 0 | 0 | -0.5 | 0 |
| 12 | NC1=CC(C2=CC=C(C=C2)Cl)=NN1 | -2 | 0 | -1 | 2 | 1 | 0 | 0 |
| 13 | N1(/N=C(\CC1=O)/N)c1ccccc1 | 0 | 0 | -1 | 0 | 0 | 0 | 0 |
| 14 | CC1=C(C=NN1)C2=CC=CC=C2 | -2 | 1.5 | 3 | 0 | 0 | 0 | 0 |
| 15 | c1cnc(cc1)c1cccc(c1)C(O)=O | 0 | 0 | 0 | -1 | 0 | 0 | 0 |
| 16 | NC1=CC=C(N2N=C(C)C=C2C)C=C1 | 0 | 0 | 0 | -1 | 0 | 0.5 | 0 |
| 17 | s1c(ccc1c1ncccc1)C(=O)O | 0 | 0 | 0 | -1 | 0 | 0.5 | 0 |
| 18 | n1(nc(c(n1)C)C(=O)O)c1ccccc1 | 0 | 0 | 0 | 0 | 0 | 0 | 0 |
| 19 | Clc1cncc(Cl)c1 | 0 | 0 | 0 | 0 | 0 | 0 | 0 |
| 20 | c1ccc(cc1)Cc1ccncc1 | 28 | 3 | 1 | 2 | 1 | 2 | 0 |
| 21 | CNCc1cccc(c1)c1cccnc1 | 25 | 3 | -1 | 0 | 1 | 4 | 0 |
| 22 | OC(=O)c1ccccc1n1cccn1 | 0 | 0 | 0 | 0 | 0 | 0 | 0 |
| 23 | Nc1ccc(Br)cn1 | 0 | 0 | 0 | -1 | 0 | 0.5 | 0 |
| 24 | Oc1nnc(O)c(C)c1 | 0 | 0 | 0 | -1 | 0 | 0 | 0 |
| 25 | COC(=O)c1ccnc(N)c1 | 0 | 0 | 0 | 0 | 0 | 0.5 | 0 |
| 26 | COC(=O)c1ncc2ccccc2c1 | 0 | 0 | 0 | 0 | 0 | 1 | 0 |
| 27 | Clc1ccc(Cl)nn1 | 0 | 0 | 0 | 0 | 0 | 0.5 | 0 |
| 28 | Nc1cc(Cl)nc(Cl)c1 | 0 | 0 | 0 | 0 | 0 | 0 | 0 |
| 29 | OCc1ccc(Br)cn1 | 0 | 0 | 0 | 0 | 0 | 0.5 | 0 |
| 30 | Oc1cccc(n1)C(=O)O | 0 | 0 | 0 | 0 | 0 | 1 | 0 |
| 31 | COc1ccc(cn1)C(=O)C | 0 | 0 | 0 | 0 | 0 | 0.5 | 0 |
| 32 | Cc1cc(C(=O)O)[nH]n1 | 0 | 0 | 0 | 0 | 0 | -0.5 | 0 |
| 33 | OB(O)c1cccnc1 | 0 | 0 | 0 | 0 | 0 | 4 | 0 |
| 34 | OC(=O)CCc1nc2ccccc2[nH]1 | 0 | 0 | 0 | 0 | 0 | 0.5 | 0 |
| 35 | c1ccc(cc1)c1n[nH]cc1 | -2 | 0 | -1 | 0 | 0 | 1 | 0 |
| 36 | OC(=O)CCc1cccnc1 | 0 | 0 | 0 | 0 | 2 | 1 | 0 |
| 37 | Brc1ccc2ccnc(Cl)c2c1 | 0 | 0 | 0 | -3 | 0 | ND | 0 |

| | | | | | | | | |
|---|---|---|---|---|---|---|---|---|
| 38 | Brc1c[nH]nc1C | 0 | 1.5 | 0 | 0 | 0 | 0 | 0 |
| 39 | N1CCc2ccccc2C1 | 0 | 1.5 | 2 | 0 | 0 | 1 | 0 |
| 40 | N#Cc1ccccn1 | 0 | 0 | 0 | 0 | 0 | 0.5 | 0 |
| 41 | N#Cc1ccncc1 | 0 | 0 | 0 | 0 | 0 | -0.5 | 0 |
| 42 | Oc1ncccc1O | 0 | 0 | -1 | -1 | 0 | 0.5 | 0 |
| 43 | Nc1ccccn1 | 0 | 0 | 0 | 0 | 0 | 1 | 0 |
| 44 | Oc1ccccn1 | 0 | 0 | 0 | 0 | 0 | -0.5 | 0 |
| 45 | Nc1cccnc1O | 0 | 0 | 0 | ND | 0 | 0 | 0 |
| 46 | CC(C)c1n[nH]c(c1)C(=O)O | 0 | 0 | 0 | 0 | 0 | -0.5 | 0 |
| 47 | Nc1ccncc1 | 0 | 1.5 | 0 | 0 | 0 | 0 | 0 |
| 48 | Nc1ccc(nc1)Oc1ccccc1 | 0 | 1.5 | -1 | 1 | 2 | 0.5 | 0 |
| 49 | Nc1ccc(cc1)n1cccn1 | 0 | 0 | 0 | 0 | 2 | 1 | 0 |
| 50 | COc1ncccc1N | 0 | 0 | 0 | 0 | 0 | 2.5 | 0 |
| 51 | OC(=O)c1ccnn1c1ccccc1 | 0 | 0 | 0 | -1 | 0 | 0.5 | 0 |
| 52 | CNCc1ccc(cc1)c1ccncc1 | 28.5 | 1.5 | 2 | -1 | 0 | 4 | 0 |
| 53 | OC(=O)c1ccc(cc1)c1cccnc1 | 0 | 0 | 0 | 0 | 0 | 1 | 0 |
| 54 | OC(=O)c1ccc(cc1)n1cccn1 | 0 | 0 | 0 | 0 | -1 | -0.5 | 0 |
| 55 | [O-][N+](=O)c1cccnc1N | ND | 0 | 0 | ND | 0 | ND | 0 |
| 56 | Clc1nccc2ccccc12 | 0 | 0 | 0 | -2 | 0 | -0.5 | 0 |
| 57 | Brc1cccnc1Cl | 0 | 0 | 0 | -1 | 0 | 0.5 | 0 |
| 58 | Brc1ccc(=O)[nH]c1 | 0 | 0 | 0 | 0 | -1 | 0.5 | 0 |
| 59 | [O-][N+](=O)c1ccc(Br)c2ccncc12 | 0 | 0 | 1 | ND | 0 | ND | 0 |
| 60 | OC(=O)c1cc(F)cnc1 | 0 | 0 | 0 | 0 | 0 | 1 | 0 |
| 61 | Clc1ncccc1Cl | ND | 0 | 0 | 0 | 0 | 0 | 0 |
| 62 | OC(=O)c1cccnc1N | ND | 0 | 0 | ND | 0 | 4 | 0 |
| 63 | c1ccc(nc1)N1CCOCC1 | 0 | 0 | 0 | 0 | 0 | 1 | 0 |
| 64 | O=C1CCN(N1)c1ccccc1 | 0 | 1.5 | -1 | 0 | 0 | 1 | 0 |
| 65 | COc1ncccc1C(=O)O | ND | 0 | 0 | 0 | 0 | 1 | 0 |
| 66 | Nc1ncccc1N | ND | 0 | 0 | ND | 0 | 0 | 0 |
| 67 | CCc1[nH]cc(C)n1 | 0 | 0 | 0 | -1 | 0 | 0 | 0 |
| 68 | c1ccc(cc1)c1c[nH]cn1 | 0 | 3 | 4 | 0 | 1 | 2 | 0 |
| 69 | OCc1ccc(cc1)Cn1ncnc1 | 0 | 0 | 0 | 0 | 0 | 0 | 0 |
| 70 | OC(=O)c1cc(c2ccccc2)n(C)n1 | 0 | 0 | 2 | 0 | 0 | 0.5 | 0 |
| 71 | N#Cc1cccc(c1)n1cncc1 | 0 | 3 | 1 | 0 | 0 | 2.5 | 0 |
| 72 | N#Cc1ncccc1F | 0 | 0 | 0 | 0 | 0 | 0 | 0 |
| 73 | Nc1ccc(Cl)nn1 | 0 | 0 | 0 | 0 | 0 | 0 | 0 |
| 74 | OC(=O)c1cccc(c1)c1cccnc1 | 0 | 0 | 0 | 0 | 0 | 1 | 0 |
| 75 | C#Cc1cccnc1 | 0 | 0 | 0 | 0 | 0 | 0 | 0 |
| 76 | OC(=O)c1cc(O)c2ccccc2n1 | 0 | 0 | 0 | 0 | 0 | 0.5 | 0 |
| 77 | Brc1c[nH]cn1 | 0 | 3 | 0 | -1 | 0 | 0.5 | 0 |
| 78 | OS(=O)(=O)c1ccccn1 | 0 | 0 | 0 | 0 | 0 | 1 | 0 |
| 79 | OC(=O)[C@H]1NCc2ccccc2C1 | -2 | 0 | 0 | 1 | 0 | 0 | 0 |
| 80 | Cc1ccc(nc1)S(=O)(=O)N | 0 | 0 | 0 | 0 | 0 | 0 | 0 |

**Table S2.** Screen of benzylpyridine **1a** analogues to establish CYP125 and CYP142 structure-activity relationships (SARs). Compound (100 µM) binding to purified P450 (5 µM) was quantified from the shift in the Soret band ($\Delta\lambda_{max}$, nm) of each enzyme's absorbance spectrum, relative to DMSO. [a]CYP125 is prominently high spin (HS) at resting state ($\lambda_{max}$ ~ 393 nm), so $\Delta\lambda_{max}$ values was calculated for the $\Delta\lambda_{max}$ of both the HS and low spin (LS) enzyme populations represented in the absorbance spectrum. [b]The ratio of LS/HS CYP125 provides additional indication of the extent to which compound binding stabilizes the LS, presumably inactive, state. [c]No LS or no HS maxima was present in the enzyme spectrum. ND – value not determined.

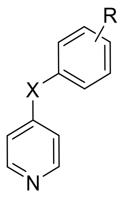


| Compound | X | R | CYP125 $\Delta\lambda_{max}$ (nm) HS[a] | CYP125 $\Delta\lambda_{max}$ (nm) LS | CYP125 LS/HS[b] | CYP142 $\Delta\lambda_{max}$ (nm) |
|---|---|---|---|---|---|---|
| **1a** | $CH_2$ | H | 2 | 27 | 1.1 | 2 |
| **2a** | NH | H | 0 | -[c] | ND | 4 |
| **2b** | NHMe | H | 1 | 25 | 0.9 | 4 |
| **2c** | NHPh | H | 2 | 28 | 1 | 4 |
| **2d** | O | H | 1 | 25 | 1 | 2 |
| **2e** | $SO_2$ | H | 0 | - | ND | 0 |
| **2f** | C=O | H | 0 | - | ND | 0 |
| **2g** | *E*-CH=CH | H | - | 29 | 1.4 | 3 |
| **2h** | $CH_2CH_2$ | H | 2 | 26 | 1 | 2 |
| **3a** | $CH_2$ | 3-OMe | 1 | 27 | 1.1 | 2 |
| **3b** | $CH_2$ | 4-OMe | 1 | 25 | 1 | 2 |
| **3d** | $CH_2$ | 3-$NH_2$ | 3 | 29 | 1.3 | 2 |
| **3e** | $CH_2$ | 4-$NH_2$ | 3 | 29 | 1.7 | 3 |
| **3f** | $CH_2$ | 4-NHCOMe | - | 13 | ND | 2 |
| **3g** | $CH_2$ | 4-$NHSO_2Me$ | - | 30 | 2.7 | 0 |
| **4a** | NH | 3-$NH_2$ | 1 | 22 | 0.9 | 3 |
| **4b** | NH | 4-$NH_2$ | 1 | 31 | 1.3 | 3 |
| **4c** | NH | 3-$CO_2Me$ | 1 | 27 | 1 | 3 |
| **4d** | NH | 4-$CO_2Me$ | 2 | 26 | 1 | 3 |
| **4e** | NH | 3-$CO_2H$ | 0 | - | ND | 0 |
| **4f** | NH | 4-$CO_2H$ | 1 | - | ND | 0 |
| **4g** | NH | 3-$CH_2OH$ | 0 | - | ND | 1 |
| **4h** | NH | 3-Br | - | 31 | 1.4 | 4 |
| **4i** | NH | 4-Br | 2 | 29 | 1.2 | 4 |

**Table S3**. X-ray crystallography data and refinement statistics.

| | **CYP125-5m (PDB 7ZIC)** | **CYP125-5j (PDB 7ZGL)** | **CYP125-5g (PDB 8S4M)** | **CYP142-1a (PDB 8S53)** | **CYP142-5m (PDB 7P5T)** | **CYP142 -5j (PDB 7QQ7)** |
|---|---|---|---|---|---|---|
| **Data collection** | | | | | | |
| Space group | C 1 2 1 | C 1 2 1 | C1 2 1 | P $2_1 2_1 2_1$ | P $2_1 2_1 2_1$ | P $2_1 2_1 2_1$ |
| Cell dimensions | | | | | | |
| a, b, c (Å) | 136.23, 68.68, 144.47 | 136.31, 69.33, 144.89 | 136.90, 68.85, 144.12 | 55.46, 65.77, 130.66 | 55.73, 65.72, 129.07 | 55.31, 65.19, 128.41 |
| α, β, γ (°) | 90,00 93.97, 90.00 | 90.00, 94.41, 90.00 | 90.00, 93.92, 90.00 | 90.00, 90.00, 90.00 | 90.00, 90.00, 90.00 | 90.00, 90.00, 90.00 |
| Resolution (Å) | 1.90 | 2.50 | 2.1 | 1.6 | 1.30 | 1.60 |
| No. reflections (total) | 348123 (16169) | 156569 (15019) | 260955 (15185) | 490509 (12064) | 663389 (12279) | 410853 (21072) |
| No. reflections (unique) | 104289 (5050) | 46975 (4600) | 78300 (4463) | 62563 (2491) | 116304 (5347) | 62174 (3085) |
| $R_{merge}$ | 0.064 (0.898) | 0.148 (0.908) | 0.071 (0.861) | 0.062 (0.831) | 0.053 (0.705) | 0.072 (0.795) |
| I / σI | 9.2 (1.2) | 4.6 (1.1) | 9.1 (1.4) | 17 (1.3) | 11.4 (1.0) | 12.1 (2.2) |
| CC 1/2 | 0.996 (0.518) | 0.969 (0.599) | 0.995 (0.585) | 0.998 (0.657) | 0.998 (0.566) | 0.998 (0.841) |
| Completeness (%) | 99.4 (99.0) | 99.9 (99.9) | 99.9 (100) | 97.9 (79.9) | 99.28 (99.10) | 100.00 (100.00) |
| Multiplicity | 3.3 (3.2) | 3.3 (3.3) | 3.3 (3.4) | 7.8 (4.8) | 5.7 (2.3) | 6.6 (6.5) |
| **Refinement** | | | | | | |
| Rwork / Rfree | 0.188 / 0.224 | 0.210 / 0.251 | 0.212 / 0.246 | 0.156/0.175 | 0.137 / 0.162 | 0.164 / 0.183 |
| R.m.s. deviations | | | | | | |
| Bond lengths (Å) | 0.007 | 0.004 | 0.003 | 0.012 | 0.009 | 0.009 |
| Bond angles (°) | 0.801 | 0.651 | 0.619 | 1.154 | 1.41 | 1.072 |

**Table S4.** Inhibition of CYP125 and CYP142 catalytic activity in vitro. The concentration of compound to inhibit 50% of CYP125 (0.5 µM) or CYP142 (1 µM) catalyzed turnover of cholest-4-en-3-one (5 µM) ($IC_{50}$ value) was quantified by LC-MS. Inhibition equilibrium constants ($K_I$) were estimated by Cheng-Prusoff method using cholest-4-en-3-one $K_m$ CYP125 = 2.1 µM, CYP142 = 0.36 µM. "-" – not determined. Compound structures shown below.

| | **CYP125A1** | | **CYP142A1** | |
|---|---|---|---|---|
| **Compound** | **$IC_{50}$ (µM)** | **$K_I$ (µM)** | **$IC_{50}$ (µM)** | **$K_I$ (µM)** |
| **1a** | - | - | - | - |
| **2a** | - | - | - | - |
| **3f** | - | - | - | - |
| **5a** | 4.3 ± 0.5 | 1.3 | - | - |
| **5b** | - | - | - | - |
| **5c** | 26 ± 2.4 | 7.7 | - | - |
| **5d** | 0.79 ± 0.09 | 0.23 | 2.5 ± 2.4 | 0.17 |
| **5e** | 9.2 ± 0.89 | 2.7 | 33 ± 3.4 | 2.2 |
| **5f** | 3.6 ± 0.41 | 1.1 | - | - |
| **5g** | 25 ± 2.4 | 7.1 | 34 ± 3.4 | 2.3 |
| **5i** | 33 ± 3.7 | 9.8 | - | - |
| **3g** | - | - | - | - |
| **5j** | 1.5 ± 0.16 | 0.44 | 2.3 ± 0.30 | 0.15 |
| **5k** | 0.91 ± 0.22 | 0.27 | 3.6 ± 0.36 | 0.24 |
| **5l** | 18 ± 1.9 | 5.4 | 16 ± 1.6 | 1.1 |
| **5m** | 0.35 ± 0.04 | 0.10 | 0.67 ± 0.07 | 0.05 |
| **5n** | 12 ± 1.2 | 3.6 | - | - |
| **5o** | 4.2 ± 0.48 | 1.2 | 6.1 ± 0.60 | 0.41 |
| **5p** | 10 ± 0.95 | 3.1 | 22 ± 2.4 | 1.5 |

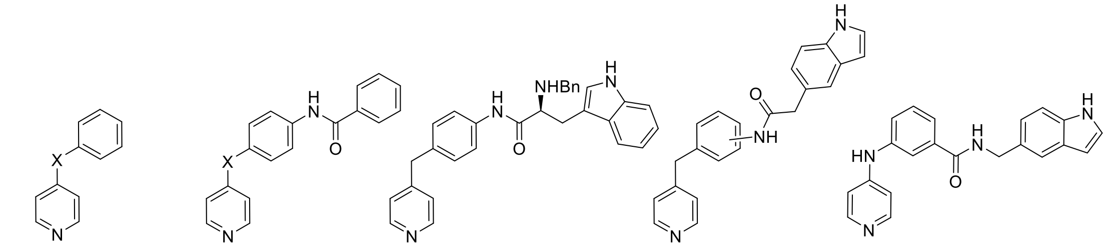


**1a** X = $CH_2$
**2a** X = NH

**5a** X = $CH_2$
**5b** X = NH
**5c** X = NMe

**5d**

**5e** 3-C
**5f** 4-C

**5g**

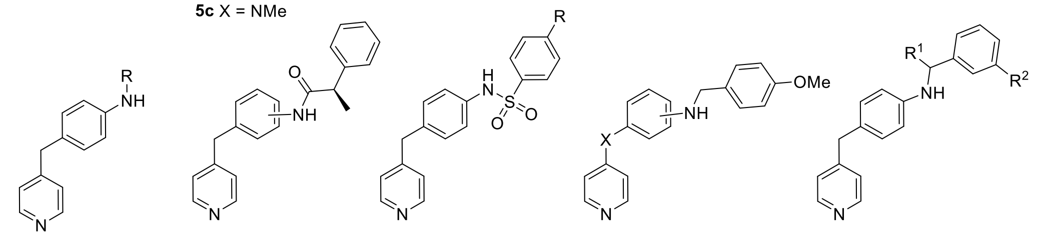


**3f** R = COMe
**3g** R = $SO_2Me$

**5h** 3-C
**5i** 4-C

**5j** R = H
**5k** R = $CF_3$

**5l** 3-C, X = $CH_2$
**5m** 4-C, X = $CH_2$
**5n** 4-C, X = NMe

**5o** $R^1$ = Me, $R^2$ = H
**5p** $R^1$ = H, $R^2$ = $SO_2Me$

**Table S5.** CYP125/142 ligands inhibit the growth of extracellular *Mtb* (H37Rv) on cholesterol. *Mtb* (H37Rv) was cultured in media that contained cholesterol as the only source of carbon and treated with compounds (0-50 µM) for 1-2 weeks. Growth inhibition was quantified as the concentration of compound required to reduce resazurin reduction (MABA) by 99% relative to DMSO control ($MIC_{99}$), or ATP-dependent luminescence (ATP) by 50% relative to DMSO treated controls ($IC_{50}$) in independent replicate experiments. "-" – not determined.

| | **MABA-$MIC_{99}$ (µM)** | **ATP-$IC_{50}$ (µM)** | |
|---|---|---|---|
| **Compound** | **W2** | **W1** | **W2** |
| **1a** | - | - | - |
| **2a** | - | - | - |
| **3f** | - | - | - |
| **5a** | 25 | 19 | 38 |
| **5b** | - | - | - |
| **5c** | >50 | 9.4 | 19 |
| **5d** | 25 | 4.7 | 4.7 |
| **5e** | 13 | 2.3 | 19 |
| **5f** | >50 | >50 | >50 |
| **5g** | 13 | 4.7 | 9.4 |
| **5h** | 25 | 9.4 | 19 |
| **5i** | 25 | 9.4 | 4.7 |
| **3g** | - | - | - |
| **5j** | 50 | 38 | 50 |
| **5k** | 25 | 2.3 | 2.3 |
| **5l** | 19 | 0.59 | 2.3 |
| **5m** | 1.5 | 0.15 | 1.2 |
| **5n** | 50 | 4.7 | 9.4 |
| **5o** | 19 | 0.15 | 0.59 |
| **5p** | 13 | 1.2 | 4.7 |
| **p-AS** | 0.19 | 0.04 | 0.29 |
| **Isoniazid** | <0.1 | <0.1 | <0.1 |

**Table S6.** Activity of CYP125/142 inhibitors against multi-drug resistant *Mtb.* Inhibition of drug susceptible H37Rv *Mtb* or isoniazid and rifampicin resistant MDR-TB (K26b00MR 113) growth by 90% ($MIC_{90}$) was calculated from the difference in resazurin reduction (MABA) relative to DMSO-treated controls 1- and 2-weeks post-compound treatment in replicate experiments.

| | **Week 1** | | **Week 2** | |
|---|---|---|---|---|
| **Compound** | **H37Rv** | **MDR-TB** | **H37Rv** | **MDR-TB** |
| **5m** | 0.78 | 0.39 | 6.25 | 12.5 |
| Isoniazid | 0.19-0.39 | 12.5-25 | 0.39 | 50 |

**Table S7.** Antimicrobial activity of CYP125/142 inhibitors on glucose and activity of lead compound **5m** against H37Rv *Mtb* cultured on glycerol. The concentration of compound required to inhibit the growth of H37Rv *Mtb* on media that contained either glucose or glycerol as the sole source of carbon by 50% ($IC_{50}$) was determined at 2 time points post-compound treatment from either the relative reduction in ATP-dependent luminescence or resazurin reduction (MABA) relative to DMSO treated controls. "**-**" – not determined.

| | **Glucose Media** | | **Glycerol** | |
|---|---|---|---|---|
| | **ATP $IC_{50}$ (μM)** | | **MABA $IC_{50}$ (μM)** | |
| **Compound** | **Week 1** | **Week 2** | **10-days** | **21 days** |
| **5a** | 19 | 50 | - | - |
| **5c** | 19 | >50 | - | - |
| **5d** | 4.7 | 2.3 | - | - |
| **5e** | 1.2 | 9.4 | - | - |
| **5g** | 2.3 | 4.7 | - | - |
| **5h** | 1.2 | 19 | - | - |
| **5i** | 9.4 | 19 | - | - |
| **5j** | 38 | 19 | - | - |
| **5k** | 19 | 4.7 | - | - |
| **5l** | 9.4 | 2.3 | - | - |
| **5m** | 1.2 | 2.3 | 3.5 | 19 |
| **5n** | 9.4 | 19 | - | - |
| **5o** | 2.3 | 2.3 | - | - |
| **5p** | 2.3 | 4.7 | - | - |
| p-AS | <0.04 | 0.07 | - | - |
| Isoniazid | <0.1 | 0.24 | - | - |
| Pyrazinamide | - | - | 0.9 | 2.8 |

**Table S8.** Inhibition of human liver microsomal P450s. Inhibition constants ($IC_{50}$ values, µM) of CYP125/142 inhibitors **5d**, **5k**, and **5m** were determined for select human P450 isoforms using the following substrates: CYP1A-ethoxyresorufin, CYP2C19- mephenytoin, CYP2C9-tolbutamide, CYP2D6-dextromethorphan, CYP3A4-midazolam/testosterone.

| Compound | **CYP1A** | **CYP2C19** | **CYP2C9** | **CYP2D6** | **CYP3A4** |
|---|---|---|---|---|---|
| **5d** | 0.11 | <0.10 | 0.14 | 0.46 | 0.52/1.49 |
| **5k** | 5.95 | 3.1 | 5.27 | 0.18 | 0.26/1.25 |
| **5m** | 0.82 | 0.44 | 1.86 | 0.33 | 0.52/1.82 |

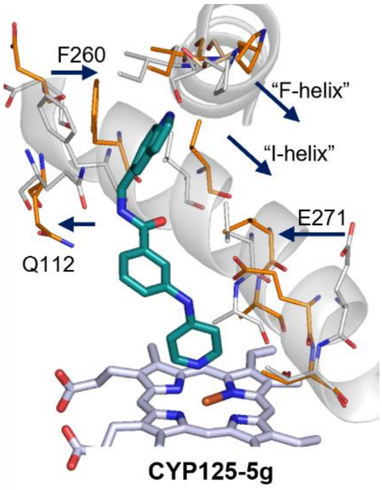


**Figure S1.** Compound 5g induces “substrate-like” shift in the CYP125 active site. (A) Aligned X-ray crystal structure of **5g-**CYP125 (aqua ligand, orange residues) and **5m-**CYP125 (grey residues, ligand not shown). Arrows highlight amino acids in the F- and I-helices that move inwards into a substrate-like conformation in the **5g-**CYP125 structure, and the significantly re-orientation of E271 to hydrogen bond with the benzylic amine.

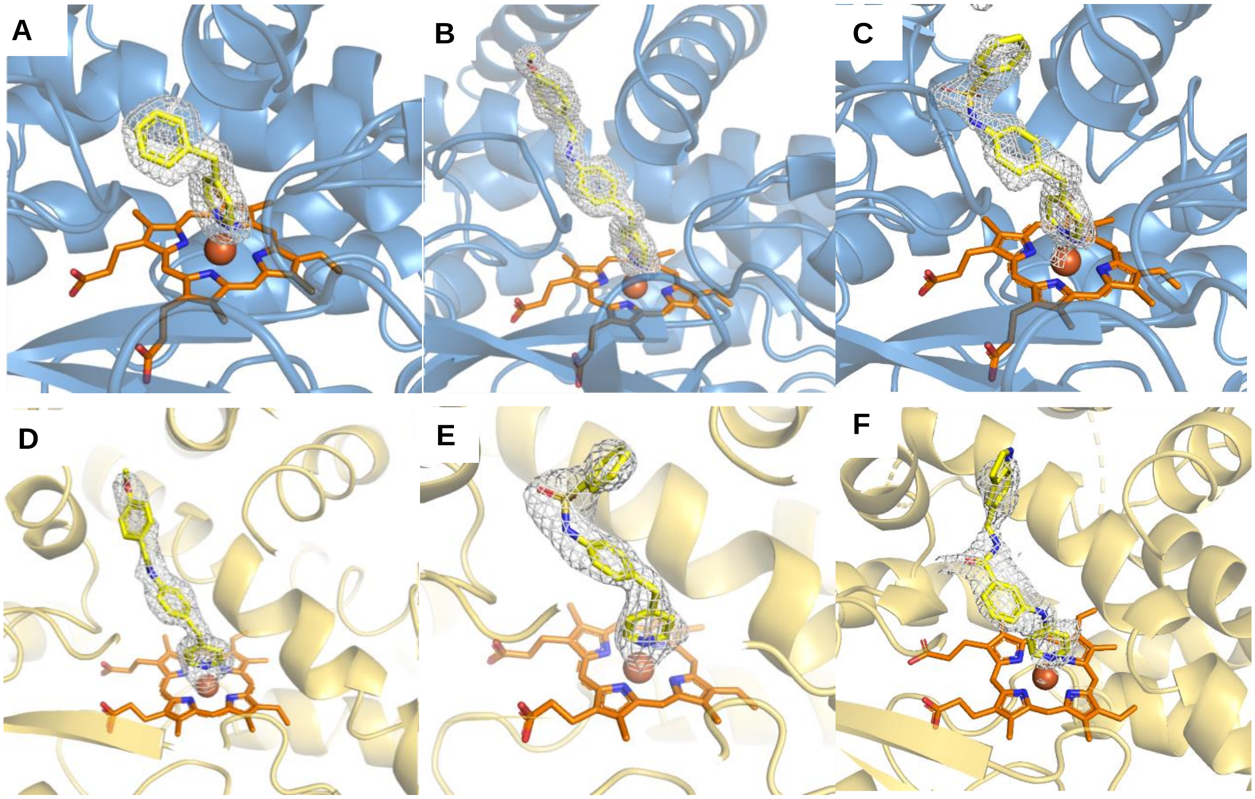


**Figure S2.** Electron density maps for ligands shown in Figures 1a and 3a-d. Polder maps (grey mesh) are contoured to 3σ, ligands are shown as yellow sticks, in complex with either CYP125 (yellow cartoon), or CYP142 (blue cartoon). (A) (CYP142-**1a-i**), (B) CYP142-**5m**; (C0 CYP142-**5j**; (D) CYP125-**5m**; (E) CYP125-**5j**; (F) CYP125-**5g**.

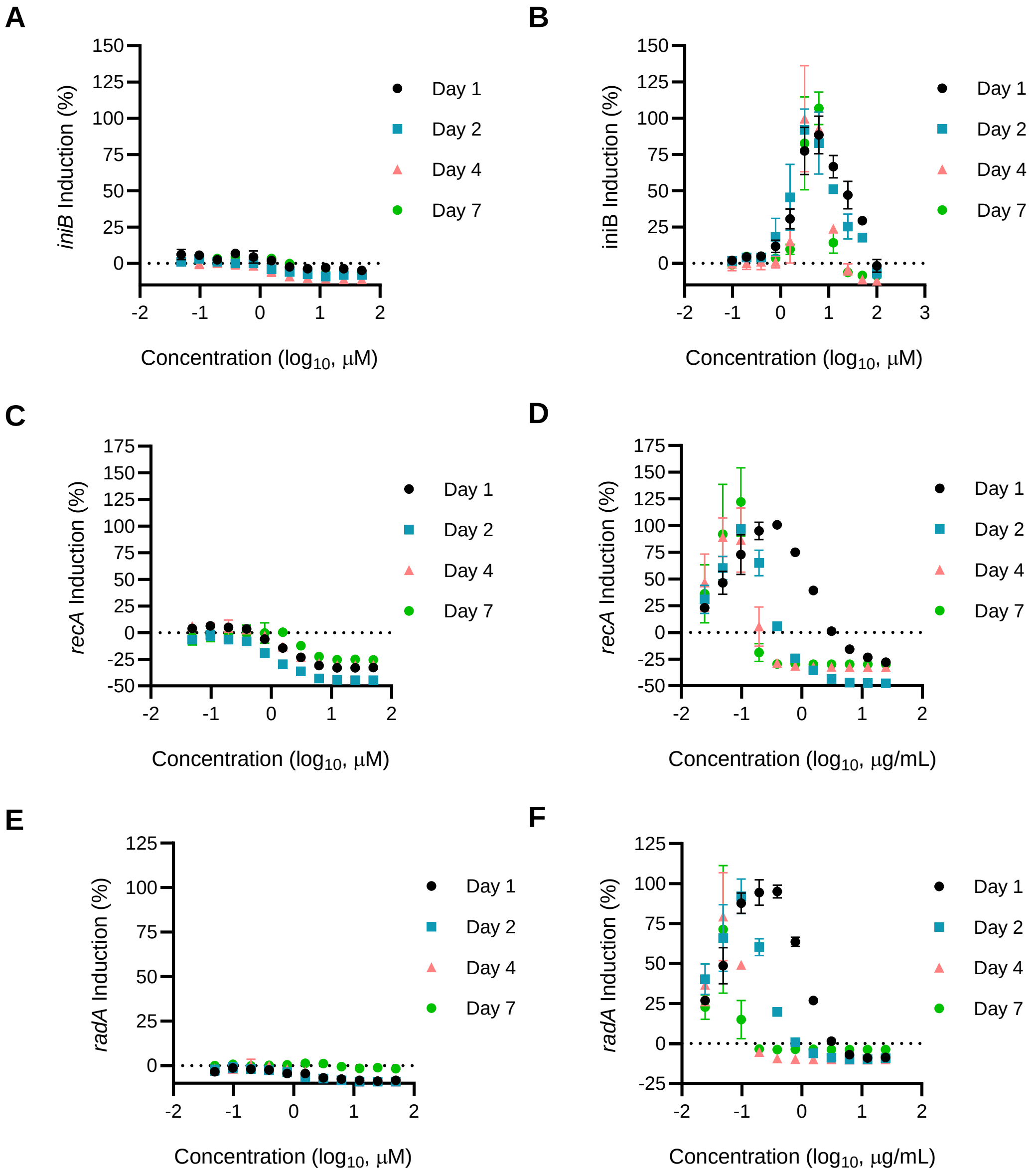


**Figure S3.** Transcriptional reporter assays to determine **5m** mechanisms of action. Induction of bioluminescent reporter for cell wall damage response (*iniB*) or DNA damage response (*recA* or *radA)* after *Mtb* were treated with compound 5m (A, C, E) or positive control compound SQ109 (B) or moxifloxacin (D, F). Signal intensity of the reporter was adjusted as a %max signal induced by positive control, and data are plotted as mean values ± SD of n=2 replicates.

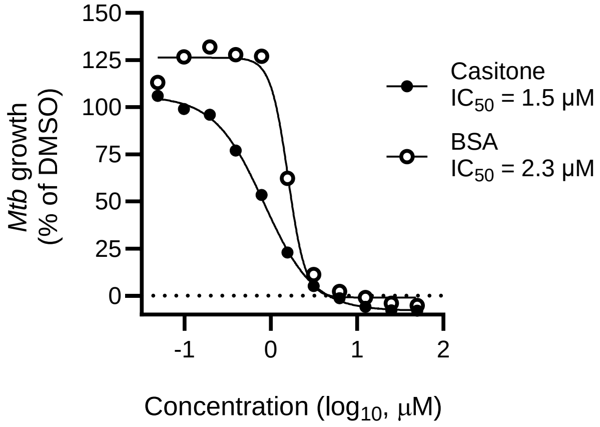


**Figure S4.** Non-specific protein binding reduces extracellular antitubercular activity of CYP125-142 inhibitors. Dose-response curves showing the inhibition of H37Rv *Mtb* growth by compound **5m** when bacteria are cultured on cholesterol media supplemented with either casitone or bovine serum albumin (BSA). Bacterial growth quantified by ATP-luminescence 2-weeks post-compound treatment and are reported as a percent of the DMSO-treated control.

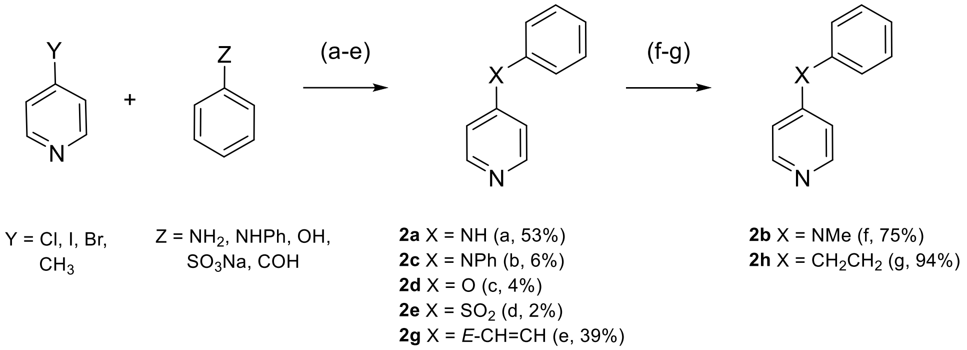


**Scheme S1.** Synthesis of **1a-i** analogue to explore SAR of benzylic position. *Reagents and conditions:* (a) Y=Cl.HCl, Z=$NH_2$, HCl (37%), EtOH, 90 °C, 20 h; (b) Y=Br.HCl, Z=NHPh, $K^tOBu$, $Pd(OAc)_2$, *rac*-BINAP, toluene, 70 °C, 16 h; (c) Y=Cl.HCl, Z=OH, Cu(s) powder, $Cs_2CO_3$, DMF, 100 °C, 18 h; (d) Y=I, Z=$SO_3Na$, *L*-proline sodium salt, CuI, DMSO, 80 °C, 44 h; (e) Y=$CH_3$, Z=COH, $Ac_2O$, 140 °C, 24 h; (f) **2a**, MeI, $K^tOBu$, DMF, r.t. 19 h; (g) **2g**, $H_2$(g), Pd/C, EtOH, r.t., 20 h.

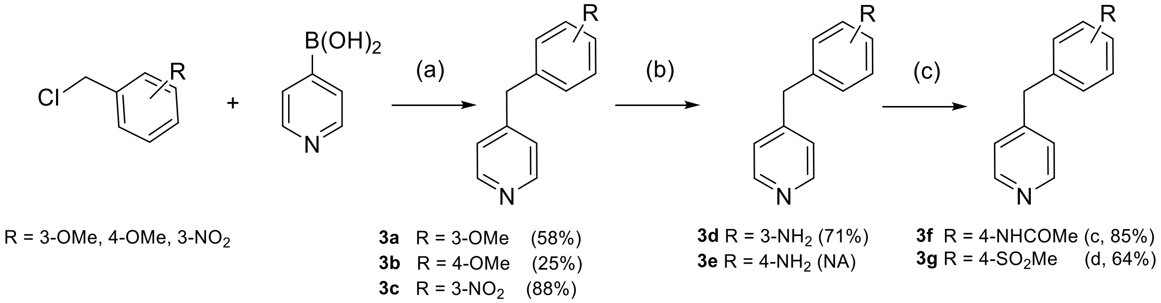


**Scheme S2.** Synthesis of **1a-i** analogues to explore SAR of "linker". *Reagents and conditions:* (a) $Pd(PPh_3)_4$, $Na_2CO_3$, DME:$H_2O$ (2:1), 100 °C, 4 h; (b) **3c**, Pd/C, $N_2H_4.xH_2O$, EtOH, 90 °C, 2 h; (c) **3f** (d) **3g**. N/A – obtained from commercial sources.

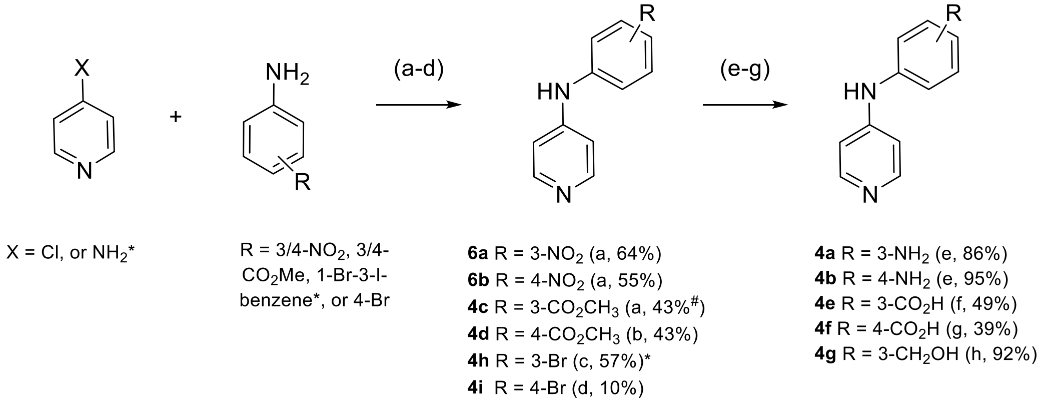


**Scheme S3.** Synthesis of **2a** analogues to explore SAR of "*linker*"*:* (a) X = Cl.xHCl, HCl (37%), EtOH, 90 °C, 20 h; (b) X = Cl.xHCl, R=4-$CO_2Me$, AcOH, $LiOH.H_2O$, MeOH:$H_2O$:THF, r.t., 4 h; (c) X = $NH_2$, 1-bromo-3-iodobenzene, $Pd_2(dba)_3$, DPPF, $Na^tOBu$, toluene, 115 °C, 24 h; (d) $Pd_2(dba)_3$, IPr.HCl, $K^tOBu$, 1,4-dioxane, 100 °C, 21 h; (e) **6a/b**, $SnCl_2.2H_2O$, HCl (37%), EtOH, 0-80 °C, 1−3 h; (f) **4c**, $LiOH.H_2O$,

$MeOH:H_2O:THF$, r.t., 4 h; (g) **4d**, KOH(aq), EtOH, reflux, 2 h; (h) **4c**, $LiAlH_4$, THF, 0 °C-r.t., 20 h.
#impure mixture of methyl/ethyl ester.
